# Contractile reprogramming of cardiac pericytes by MEK inhibition promotes arteriologenesis of the ischemic heart

**DOI:** 10.1101/2020.11.10.376772

**Authors:** Elisa Avolio, Rajesh Katare, Anita C Thomas, Andrea Caporali, Daryl Schwenke, Marco Meloni, Massimo Caputo, Paolo Madeddu

## Abstract

**Background:** The development of collateral arteries after a myocardial infarction (MI) was intensively studied, while the mechanism by which pericytes (PCs) contribute to arteriologenesis remains unexplored. This study aimed to 1) investigate if cardiac PCs gain functional features of contractile vascular smooth muscle cells (VSMCs) *in vitro*, and 2) determine if this potential can be evoked pharmacologically to encourage heart arteriologenesis *in vivo*.

**Methods:** PCs were immunosorted as CD31^neg^/CD34^pos^ cells from human and mouse hearts. Contractile reprogramming was induced by either depletion of growth factors or addition of PD0325901, a clinically available MEK inhibitor. Next generation RNA-Sequencing was performed in naïve and differentiated human PCs to assess the whole-transcriptome profile. Three *in vivo* studies were conducted in C57BL6/J mice to determine: 1) the ability of human PCs to promote arteriole formation when implanted subcutaneously within PD0325901-containing Matrigel plugs, 2) the effect of orally administered PD0325901 on the arteriole density of normoperfused hearts, and 3) the possibility of promoting capillary formation and muscularization of the infarcted heart through the same pharmacological approach.

**Results:** Removal of epidermal growth factor (EGF) and basic fibroblast growth factor (bFGF) from the culture medium induced the differentiation of PCs into contractile VSMC-like cells. Because both growth factors induce the extracellular signal-regulated kinase 1/2 (ERK1/2) signalling, we attempted to induce PC differentiation *in vitro* and *in vivo* using PD0325901. RNA-sequencing revealed that differentiated PCs were enriched in transcripts associated with smooth muscle contraction and biological function. PD0325901-treated PCs rapidly acquired antigenic and functional features of contractile VSMCs *in vitro*. Moreover, human PCs formed new arterioles when implanted subcutaneously within PD0325901-containing Matrigel plugs in mice. Oral administration of PD0325901 for two weeks increased the density and expression of contractile proteins in small-calibre arterioles of the murine heart, thereby increasing myocardial perfusion. Similarly, PD0325901 induced reparative arteriologenesis and capillarization, reduced the scar, and improved left ventricular performance in a murine model of MI.

**Conclusion:** We propose a novel method to promote the heart vascularization through the pharmacological modulation of resident mural cells. This novel approach could have an immediate impact on the treatment of coronary artery disease.

**Clinical perspective:** *What is new?:*

- Human myocardial pericytes have intrinsic vascular plasticity that can be pharmacologically evoked using PD0325901, a clinically available MEK inhibitor.
- In mice, the pharmacological inhibition of ERK1/2 signalling, by the oral administration of PD0325901 for 2 weeks, encouraged the heart arteriologenesis through pericyte differentiation.
- In a preclinical mouse model of myocardial infarction, the oral administration of PD0325901 for 2 weeks induced reparative arteriologenesis and capillarization, reduced the scar, and improved left ventricular performance.

*What are the clinical implications?:*

- This novel drug-based therapeutic approach is readily available to all patients.
- Therefore, it could have an immediate clinical impact for the treatment of coronary artery disease and other heart conditions associated with deficient coronary vascularization.

## Introduction

Cardiac repair after myocardial infarction (MI) results from a finely orchestrated series of events, including the appropriate and timely growth of pre-existent collateral arteries.^1^ Vascularization is also supported by angiogenesis, i.e. the formation of new capillaries, which eventually mature into new arterioles through the recruitment of mural cells.^2^ Patients capable of developing a good coronary circulation after an MI have a significantly reduced mortality risk compared with patients who have a poor coronary circulation.^3^ Therefore, there is a tremendous interest in deploying new therapies to reprogramming the vascularization potential of resident cardiac cells.

Pericytes (PCs) are abundantly associated with capillaries and microvessels in the heart.^4^ They are considered mesodermal precursors, which share some antigenic markers with other stromal cells, such as myofibroblasts, but have distinct functional roles in vascular stabilization and remodeling.^4-9^ A lineage tracing study showed that epicardial PCs are the ancestors of coronary vascular smooth muscle cells (VSMCs) in the developing murine heart.^10^ Moreover, PCs participate in the “no-reflow” coronary vasoconstriction during post-ischemic reperfusion.^11^ Nonetheless, it remains unknown if cardiac PCs can be pharmacologically reprogrammed to acquire a contractile phenotype, thereby enhancing the muscularization of nascent arterioles in the infarcted heart.

The present study aimed to determine the contribution of PCs to myocardial arteriologenesis. Results indicate that cultured cardiac PCs can switch to a VSMC-like phenotype following pharmacological inhibition of the extracellular signal-regulated kinase 1/2 (ERK1/2). The same pharmacological approach induced myocardial arteriologenesis in healthy and infarcted mice, which was combined with increased capillarization, reduced scars and improved ventricular performance in the latter. These findings open new therapeutic opportunities to treat coronary artery disease.

## Methods

Detailed procedures are described in the **Supplemental material online**.

### Ethics

This study complies with the guidelines of the Declaration of Helsinki. Discarded material from congenital heart defect surgery was obtained with adult and paediatric patients’ custodians’ informed consent (ethical approval 15/LO/1064 from the North Somerset and South Bristol Research Ethics Committee). Donors and samples characteristics are described in **Supplementary Table 1**.

Animals studies were covered by licenses from the British Home Office (30/3373 and PFF7D0506) and the University of Otago, New Zealand (AEC10/14), and complied with the EU Directive 2010/63/EU. Procedures were carried out according to the principles stated in the Guide for the Care and Use of Laboratory Animals (The Institute of Laboratory Animal Resources, 1996). Report of results is in line with the ARRIVE guidelines.

### Derivation of primary cultures of cardiac PCs

Human and mouse PCs were immunosorted as CD31^neg^/CD34^pos^ cells from human and mouse myocardial samples. Cells were expanded in full ECGM2 medium (PromoCell).

### *In vitro* studies

Differentiation of PCs and coronary artery VSMCs (CASMCs) employed either growth factors (GFs) depletion or PD0325901 (250 nM) treatment. Functional *in vitro* assays included antigenic profile (by real-time qPCR, immunocytochemistry and western blotting), contraction (embedding of cells in collagen gels), migration (wound healing), angiogenesis (2D-matrigel), calcium flux (Fluo-4 dye-based visualisation of calcium) and production of extracellular matrix (ECM). When required, cells were stimulated with the vasoconstrictor Endothelin-1 (ET-1). Antibodies for immunofluorescence in tissues and cells, and western blotting are reported in **Supplementary Tables 2&3**. Primers are listed in **Supplementary Table 4**.

### Next-generation RNA-Sequencing

Strand-specific RNA-sequencing was carried out on total RNA in naïve and PD0325901-differentiated PCs, and CASMCs as control, using an Illumina HiSeq platform, with a 2×150bp configuration, ∼20M reads per sample.

### *In vivo* studies

Three randomized, controlled experiments were conducted in mice.

#### Study 1

Male C57BL6/J mice (n=4/group) were injected subcutaneously, on both flanks, with Matrigel plugs containing human PCs and either PD0325901 (500 nM) or vehicle, and sacrificed seven days later for histological study of PC plasticity.

#### Study 2

Female C57BL6/J mice received either PD0325901 (orally, 10 mg/kg/day) or vehicle for fourteen days (n=11/group). Endpoints were myocardial perfusion (assessed using fluorescent microspheres), left ventricular (LV) performance (echocardiography), and vascularization (immunohistochemistry).

##### Study 3

Female C57BL6/J mice underwent permanent ligation of the left anterior descending (LAD) coronary artery, followed by oral administration of PD0325901 (orally, 10 mg/kg/day) or vehicle for fourteen days (n=12/group). Endpoints included LV performance, vascularization, and scar size.

#### Statistics

Data were analyzed using Prism version 6.0 or 8.0 and expressed as means ± standard error of the mean or standard deviation. Statistical differences were determined using T-tests and ANOVAs, unless otherwise stated. Analyses of *in vivo* outcomes followed the intention-to-treat principle.

## Results

### Human cardiac PC characterization

As previously reported in neonatal hearts,^12^ CD31^neg^ alpha-Smooth Muscle Actin (αSMA)^neg^ CD34^pos^ Platelet-Derived Growth Factor Receptor Beta (PDGFRβ)^pos^ PCs were identified around capillaries and within the adventitia of arteries in adult human hearts using immunofluorescence microscopy (**Figure 1A-C**). CD31^neg^ CD34^pos^ sorted PCs grew in culture, either as a bulk population or single cell-derived clones, showing a spindle-shaped morphology and typical antigenic profile (**Figure 1D&E** and **Supplementary Figure 1**).^12^ Compared with cardiac fibroblasts, PCs express remarkably lower levels of PDGFRα (<10% of positive cells, **Figure 1E**) and transcription factor 21 (*TCF21*) (**Figure 1F&G**), thus confirming the difference between the two populations.^13-15^ Cardiac PCs secrete angiogenic factors Hepatocyte Growth Factor (HGF), Angiopoietin (ANGPT)-1 and −2 and Vascular Endothelial Growth Factor (VEGF) (**Figure 1H**), and promote the formation and stabilization of coronary artery EC (CAEC) networks in Matrigel assays *in vitro* through physical interaction with ECs (**Figure 1I**). Controls for the above experiments are supplied in **Supplementary Figure 2**.

**Figure 1.**
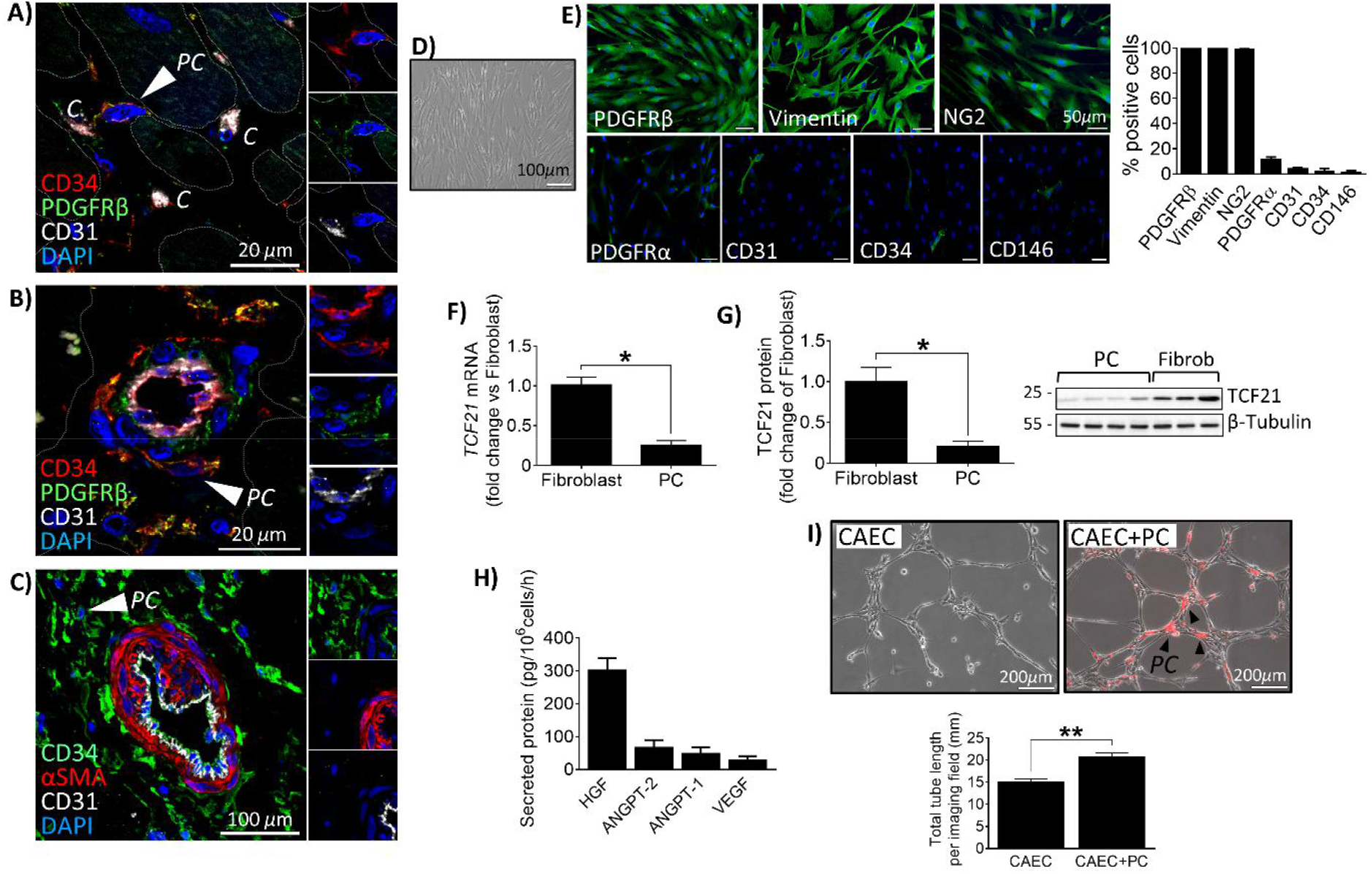
Human cardiac PC characterization. **(A-C)** Confocal immunofluorescence images of human hearts. Arrows point to CD31^neg^ αSMA^neg^ CD34^pos^ PDGFRβ^pos^ PCs around capillaries (A) and arterioles (B&C). *C*: capillary. **(D)** Brightfield microphotograph of expanded PCs. **(E)** Immunofluorescence images and bar graphs showing PC antigenic profile at passage 5 of culture. N=4. **(F&G)** Expression of TCF21 in cardiac PCs and fibroblasts evaluated by RT-qPCR (F) and western blot (G). N=3 fibroblasts and N=5 PCs. **(H)** Pro-angiogenic factors secreted by cardiac PCs. N=4. **(I)** 2D-Matrigel assay with human coronary artery ECs (CAEC) alone or in co-culture with cardiac PCs. PCs were labelled with dil (red fluorescence). Black arrows point to examples of PCs. N=5. All values are means ± SEM. * P<0.05, ** P<0.01.

### EGF and bFGF restrain cardiac PCs from differentiation into VSMC-like cells

PCs retained their original phenotype during culture expansion in GF-enriched medium (ECGM2) (*All GFs*, **Figure 2**). Using immunocytochemistry (**Figure 2A**), qPCR (**Figure 2B**) and western blotting (**Figure 2C**), we showed that GF depletion for ten days induced PCs to acquire intermediate and late-stage contractile proteins (*No GFs*). Supplementation of epidermal growth factor (EGF) and basic fibroblast growth factor (bFGF), alone and even more in combination, to basal medium, prevented the expression of smooth muscle (SM)-specific markers (*+EGF/bFGF*, **Figure 2** and **Supplementary Figure 3**). Conversely, the presence of VEGF and insulin-like growth factor 1 (IGF1) alone or together, in the absence of EGF and bFGF, induced SM-markers (*-EGF/bFGF*, **Figure 2** and **Supplementary Figure 4**). VSMC markers used for these analyses were smooth muscle myosin heavy chain (SM-MHC, gene *MYH11*), non-muscle myosin IIB (NM-MyoIIB, *MYH10)*, Smoothelin B (*SMTN*), SM-Calponin (CALP, *CNN1*), SM alpha-actin (αSMA, *ACTA2*), and Smooth Muscle Protein 22-Alpha (SM22α, *TAGLN*) (described in **Supplementary Table 5**). Both PCs and PC-derived VSMC-like cells retained the expression of Neuron-Glial antigen 2 (NG2 - *CSPG4*) and PDGFRβ (*PDGFRB*), antigens shared by PCs and VSMCs (**Supplementary Figure 5**). We confirmed the GF-modulated plasticity in single PC-derived clones (**Supplementary Figure 6**).

Human CASMCs were also responsive to GF depletion, acquiring the contractile marker SM-MHC along with upregulation of other SM-markers (**Supplementary Figure 7**).

**Figure 2.**
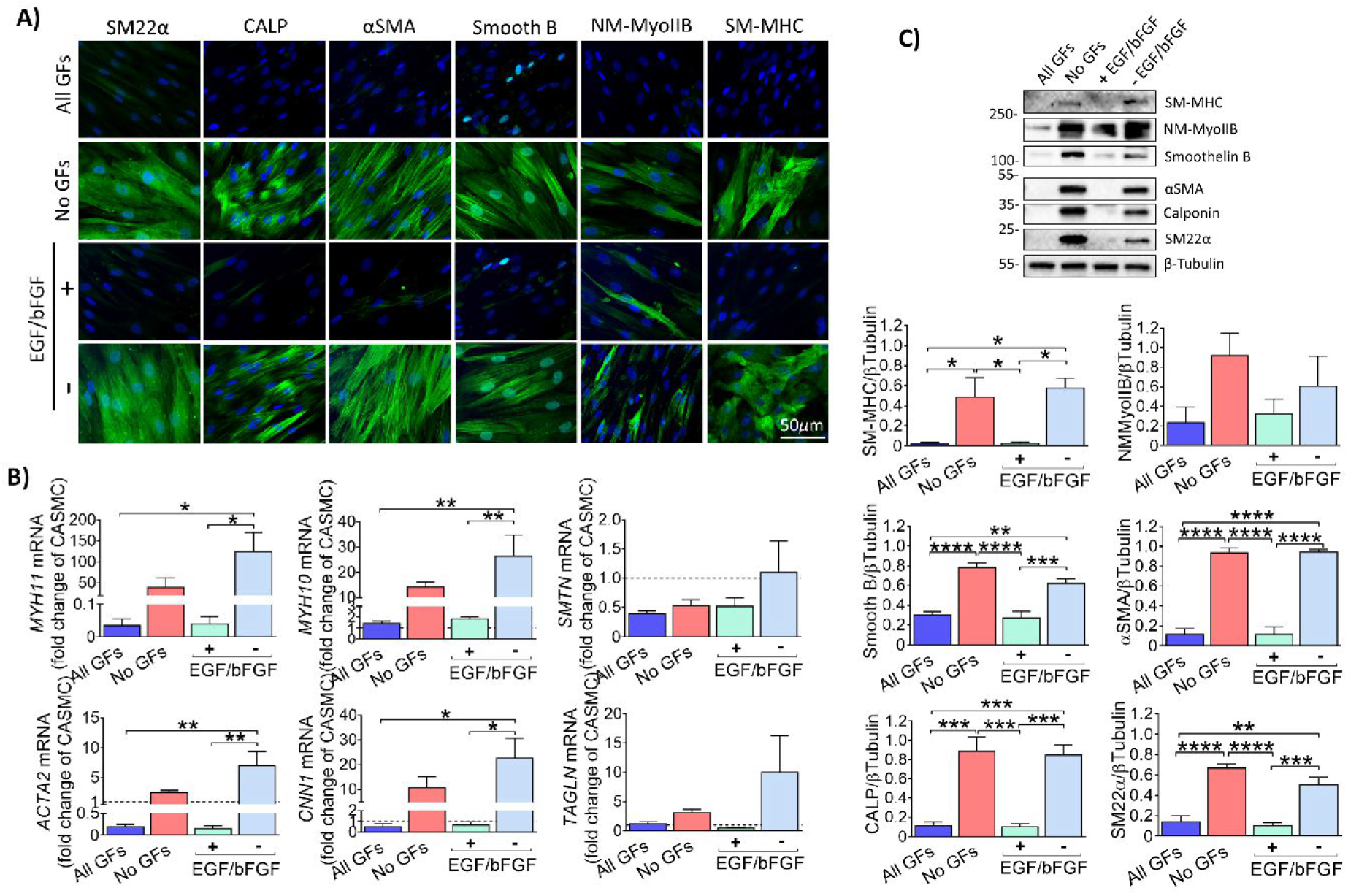
EGF and bFGF control the VSMC potential of human cardiac PCs. **(A)** Immunofluorescence images showing expression of VSMC markers by naïve and differentiated PCs when cultured with different combinations of GFs for ten days. ***All GFs***: VEGF, IGF-1, EGF, bFGF. ***No GFs***: depletion of all GFs. ***-EGF/bFGF***: only VEGF and IGF1. ***+ EGF/bFGF***: only EGF and bFGF. **(B)** Transcriptional analysis of SM-genes. mRNA data are expressed as fold change of CASMCs used as reference (dashed line at y=1). **(C)** Western blotting analysis of VSMC markers. Blots show a representative PC line. Graphs report blots densitometry. N=5. Values are means ± SEM. * P<0.05, ** P<0.01, *** P<0.001, **** P<0.0001.

### GF-depleted PCs display functional properties of contractile VSMCs

We next assessed key functional characteristics of PCs differentiated through GF depletion, namely contraction, calcium flux, migratory ability, and production of ECM (**Figure 3**). When embedded into collagen gels, differentiated PCs, but not *naïve* PCs, contracted in response to vasoconstrictor ET-1, this response being prevented by butanedione monoxime, a myosin ATPase inhibitor (**Figure 3A**). Endothelin-induced contraction is calcium-dependent; a Fluo-4 calcium flux assay showed that differentiated PCs were characterized by a greater and faster intracellular calcium mobilization in response to ET-1 compared with *naïve* PCs (**Figure 3B**). Differentiated PCs did not migrate in response to Platelet-Derived Growth Factor-BB (PDGF-BB), Stromal Derived Factor-1 alpha (SDF-1α) or VEGF-A, whereas *naïve* PCs did so (**Figure 3C**). Moreover, differentiated PCs produced lower amounts of fibronectin (*FN1*) but more elastin (*ELN*) (**Figure 3D)**.

**Figure 3.**
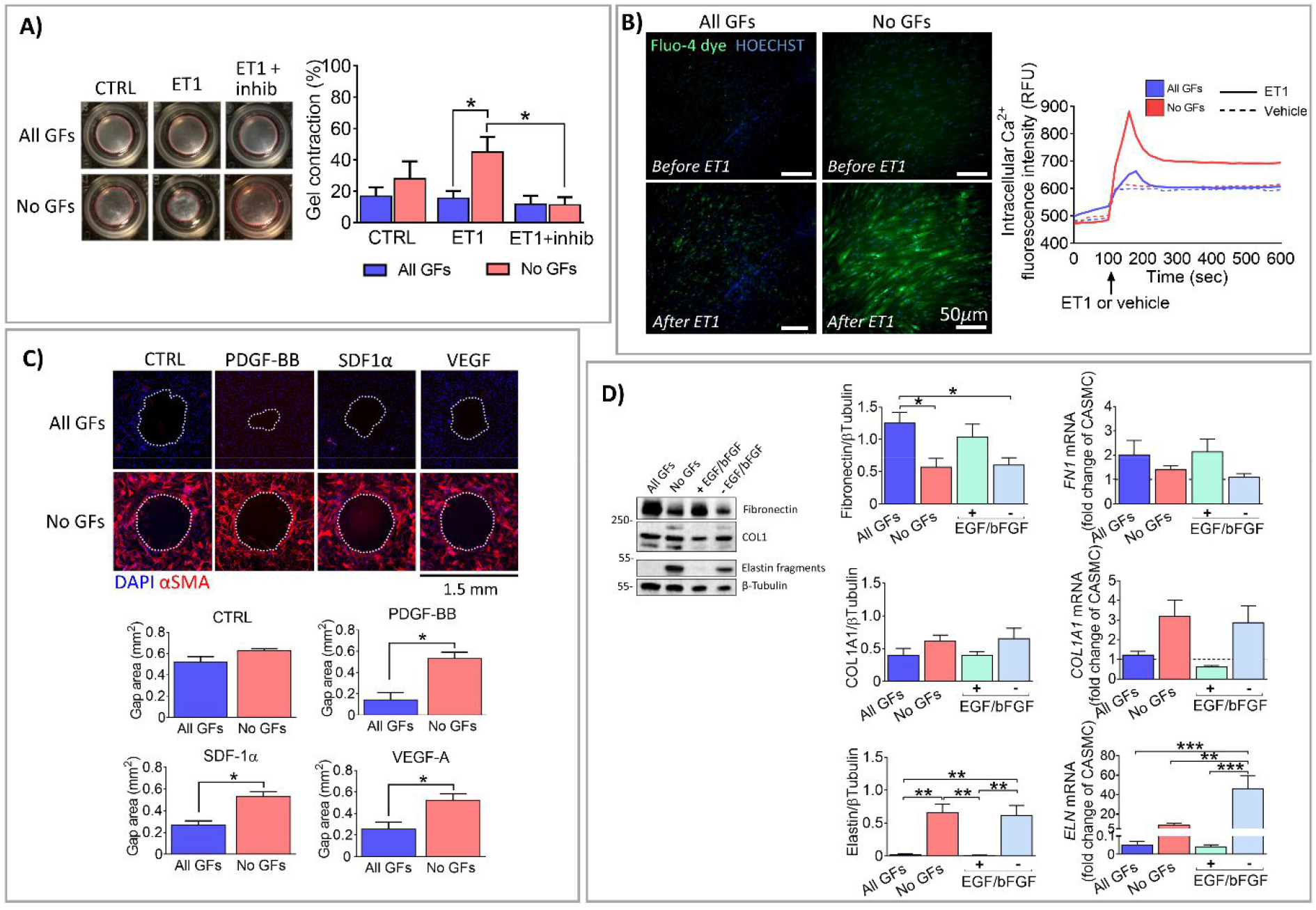
Functional assays confirm that differentiated human cardiac PCs gain properties of contractile VSMCs. Assays were performed at the end of a 10-day culture with or without GFs. **(A) Contraction assay**. Cells were embedded in collagen gels, treated with a contraction inhibitor and stimulated with Endothelin-1 (ET1). Bar graphs indicate the % of gel contraction after 24h. N=4. **(B) Fluo-4 calcium assay**. The Fluo-4 dye was loaded into the cells and these were stimulated with ET1 or vehicle. Images were recorded every 20 sec for 10 min. The intracellular calcium flux was measured as relative fluorescence units (RFU - green fluorescence of Fluo-4). N=2. The curves show 1 PC line. **(C) Gap closure migration assay**. Migration time was 24h. Absence of stimuli was the control - CTRL. Cells were stained for DAPI and αSMA at the end of the experiment. Bar graphs show the area of the final gap. N=3. **(D) Expression of extracellular matrix (ECM) markers**. Blots and bar graphs show the expression of ECM proteins and relative genes. Blots are representative of one PC line. mRNA data are expressed as fold change of CASMCs used as reference (dashed line at y=1). N=5. All values are means ± SEM. * P<0.05, ** P<0.01, *** P<0.001.

Like differentiated PCs, differentiated contractile CASMCs showed limited ability to migrate and secreted elastin (**Supplementary Figure 8**).

Together, these results indicate that GF-depleted PCs acquire not only antigenic but also typical functional features of contractile VSMCs.

### ERK1/2 signalling controls the PC switch to VSMC-like cells *in vitro*

We next analysed the signalling activated by EGF and bFGF in cardiac PCs. A phospho-kinase array screening of PCs cultured in the presence or absence of GFs showed that only EGF and bFGF activate ERK1/2 and its downstream targets (**Supplementary Figure 9A**). Western blotting confirmed the phosphorylation-activation of the EGF Receptor (EGFR)-FGFR-ERK1/2-ELK1 axis (**Supplementary Figure 9B**). Phosphorylation-activation of ELK1 (E26 transformation-specific (ETS) Like-1 protein) by ERK1/2 reportedly prevents SM genes transcription.^16, 17^ In view of a clinical application, we adopted a pharmacologic inhibitory approach based on the use of small bioactive molecules to investigate the role of this signalling in controlling PC differentiation. A pilot screening of four candidates (*Genistein*, EGFR and FGFR inhibitor; *U0126, PD98059* and *PD0325901*, dual specificity mitogen-activated protein kinase kinase (MEK)1/2 inhibitors (MEKi)), identified PD0325901 as the most potent inhibitor of MEK activity in cardiac PCs (data not shown). PD0325901 binds to an allosteric site in the MEK1/2 activation loop in an Adenosine Triphosphate (ATP)-non-competitive fashion, thereby preventing its activation and kinase activity.

Dose-response studies demonstrated that PD0325901, used at 250-500 nM, prevents ERK1/2 phosphorylation for at least 48 hrs without affecting PC viability when assessed up to 10 continuous days of culture (**Supplementary Figure 10**).

### Next-generation RNA-sequencing shows that differentiated PCs are enriched with contractile mRNA transcripts

We next performed a whole-transcriptome analysis of naïve and PD0325901-differentiated PCs (DPCs) to establish if the pharmacological treatment could shift the global cell transcriptome towards the VSMC lineage. CASMCs were used as the internal control. As shown in **Figure 4A**, the bioinformatic analysis indicated that a cluster of genes was upregulated in both DPCs and CASMCs as compared with naïve PCs. Zooming into this cluster unveiled several gene transcripts associated with SM contraction and differentiation (**Figure 4B**). Moreover, the number of genes co-expressed by DPCs and CASMCs was 3-fold higher than that shared by naïve PCs and CASMCs (**Figure 4C**).

**Figure 4.**
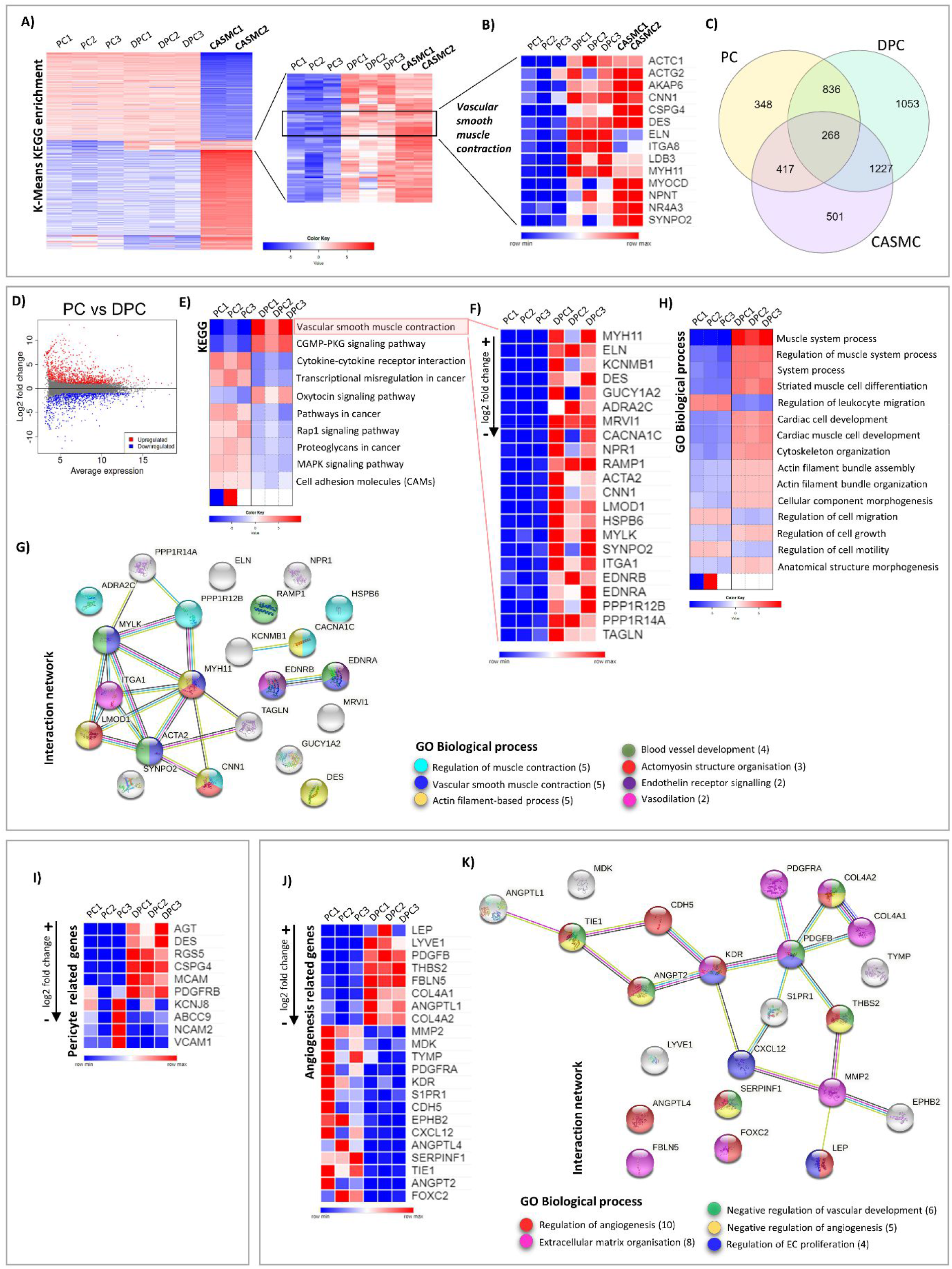
Next-generation RNA-sequencing analysis of naïve and differentiated human cardiac PCs. **(A)** K-means KEGG analysis of differentially expressed genes (DEGs) in PC, drug-differentiated PC (DPC) and CASMCs. **(B)** List of most predominant genes associated with the pathway *Vascular smooth muscle contraction*. **(C)** Venn diagram showing the number of genes expressed uniquely or shared by the three populations of cells. **(D-G)** Comparison of contractile VSCM-related genes in DPC vs naïve PC. **(D)** MA-plot representing genes differentially expressed by the two populations of cells. **(E)** List of most regulated KEGG pathways. **(F)** DEGs associated with *Vascular smooth muscle contraction*. **(G)** STRING protein-protein interaction analysis of genes in (F), and emerging gene ontology (GO) terms: Biological Process. **(H)** Main pathways resulting from the GO Biological Process analysis of DPC vs PC. **(I)** Differential analysis of pericyte related transcripts in DPC vs PC. **(J)** Analysis of angiogenesis-related DEGs in DPC vs PC. Genes were selected according to absolute log2 fold change > 1.5. **(K)** STRING network analysis of angiogenesis-related genes and emerging GO terms: Biological Process. In A, B, D, E, F, H, I and J, events upregulated are shown in red, downregulated in blue. N=3 PC, N=3 DPC, N=2 CASMC.

The contrast between DPCs vs PCs revealed 1,870 differentially expressed genes (DEGs, false discovery rate < 0.05 and absolute log2 fold change > 1), of which 1,037 upregulated and 833 downregulated (**Figure 4D**). The KEGG pathway *Vascular smooth muscle contraction* evidenced several genes upregulated in DPCs (**Figure 4E&F**, min log2 fold change +1.8, max +12.7). These genes were further analyzed in a STRING network, which showed that 13 proteins encoded by those genes have a strong biological connection (high confidence interaction score of 0.7 and protein-protein interaction (PPI) enrichment p-value < 1e-16) (**Figure 4G**). Among the main biological processes emerging from this analysis, there is *Vascular smooth muscle contraction* (*MYH11*, log2 fold change = +12.7; *ACTA2*, +4.5; endothelin A/B receptors, *EDNRA/B*, both +3.7; SM Myosin light chain kinase, *MYLK*, +3.4) and *Actomyosin structure organization (MYH11; CNN1, +4.4;* Leiomodin 1, *LMOD1, +4.4)*. Actomyosin is the molecular complex deriving by the association of actin and myosin, which ensures muscle cell contraction. The *Endothelin receptor signalling* is also instrumental in VSMC contraction. Besides, some hits of the network are involved in the control of the calcium flux and related to the *Regulation of muscle contraction* (Calcium channel, voltage-dependent, L type, alpha 1C, *CNCNA1C*, log2 fold change +5; Protein phosphatase 1 regulatory subunit 12B, *PPP1R12B*, +2.7). Other emerging processes are *Blood vessel development, Actin filament-based process* and *Vasodilation*, all related to blood vessels formation and function. As expected, the whole-genome contrast between DPCs vs PCs showed that the biological processes *Cytokine-cytokine receptor interaction* and *MAPK signalling* were downregulated in DPCs (**Figure 4H**). A schematic view of the two main regulated pathways is further illustrated in **Supplementary Figure 12**. Again, DPCs showed an upregulation of biological processes involving the cytoskeleton and actin filament organization (**Figure 4H**). Conversely, pathways controlling cell motility and migration were downregulated (**Figure 4H**).

We also analyzed how the expression of classical pericyte-associated transcripts changed in cardiac PCs after the MEKi treatment (**Figure 4I**). Five of these genes, reportedly shared by PCs and VSMCs, were upregulated in DPCs (Angiotensinogen, *AGT*, log2 fold change +10.4; Desmin, *DES, +*9.4; Regulator of G-protein signalling 5, *RGS5*, +7.5; *CSPG4*, +4.6; and *PDGFRB*, +1.4). As expected, three transcripts recently associated with pericytes but not VSMCs of the adult human heart,^7^ were downregulated in DPCs (Potassium inwardly-rectifying channel, *KCNJ8*, −1.1; ATP-binding cassette, sub-family C member 9, *ABCC9*, −1.3; Neural Cell Adhesion Molecule 2, *NCAM2*, −2.4).

Last, we performed an analysis of angiogenesis-related genes. As shown in (**Figure 4J**), 22 genes were differentially expressed between DPCs and PCs (cut-off absolute log2 fold change > 1.5). A STRING network analysis of the encoded proteins showed 15 genes are biologically connected with a confidence interaction score of 0.7 and PPI enrichment *p*-value < 1e-16 (**Figure 4K**). Among the most downregulated genes, there are *ANGPT2* (encoding ANGPT-2, log2 fold change −4.89) and *TIE1* (Tyrosine-Protein Kinase Receptor Tie-1, −4.33), whose proteins are associated with vascular regression, and *SERPINF1* (Serpin Family F Member 1, −3.42), a potent secreted inhibitor of angiogenesis. Among the most upregulated genes, we find two factors which are known to be secreted by PCs: *LEP* (Leptin, +7.55) and *PDGFB* (Platelet-Derived Growth Factor Subunit B, +3.13). Both factors induce EC proliferation. Moreover, the first protein was previously associated with a positive pro-angiogenic outcome of PCs, while the latter plays an essential role in mural cells survival and blood vessel development. Finally, eight genes are involved in the extracellular matrix organization, a process required to support the remodelling of the microvasculature.

Altogether, these findings indicate that, while retaining a transcriptional distinction from VSMCs, DPCs are characterized by the expression of a cluster of gene transcripts instrumental to contractile function and stationary behaviour. At the same time, cells downregulate typical pericyte transcripts and acquire a pro-angiogenic profile.

### PD0325901 induces the differentiation of cardiac PCs into functional VSMC-like cells *in vitro*

The next step was to validate the phenotypical and functional properties of DPCs. The analysis of mRNA transcripts in PD0325901-treated PCs confirmed the upregulation of the contractile VSMC genes *MYH11, ACTA2, CNN1* and *TAGLN* (**Supplementary Figure 12A**). Furthermore, PD0325901-treated cells retained the expression of the common pericyte/VSMC antigens NG2 (*CSPG4*) and PDGFRβ (*PDGFRB*) (**Supplementary Figure 12B&C**), while showing downregulation of *FN1* and upregulation of *ELN* genes compared with naïve PCs (**Supplementary Figure 12D**).

As shown in **Figure 5A-D**, the addition of PD0325901 to the GF-supplemented medium induced cardiac PCs to acquire the expression of contractile proteins that characterize the VSMC-like phenotype of GF-depleted PCs. In the absence of GFs, PD0325901 further enhanced the expression of SM-MHC, a specific marker of contractile VSMCs, compared with vehicle (**Figure 5C**, western blotting analysis). Intriguingly, PD0325901 was faster than GF depletion in inducing the expression of CALP, αSMA and SM22α (**Supplementary Figure 13**).

**Figure 5.**
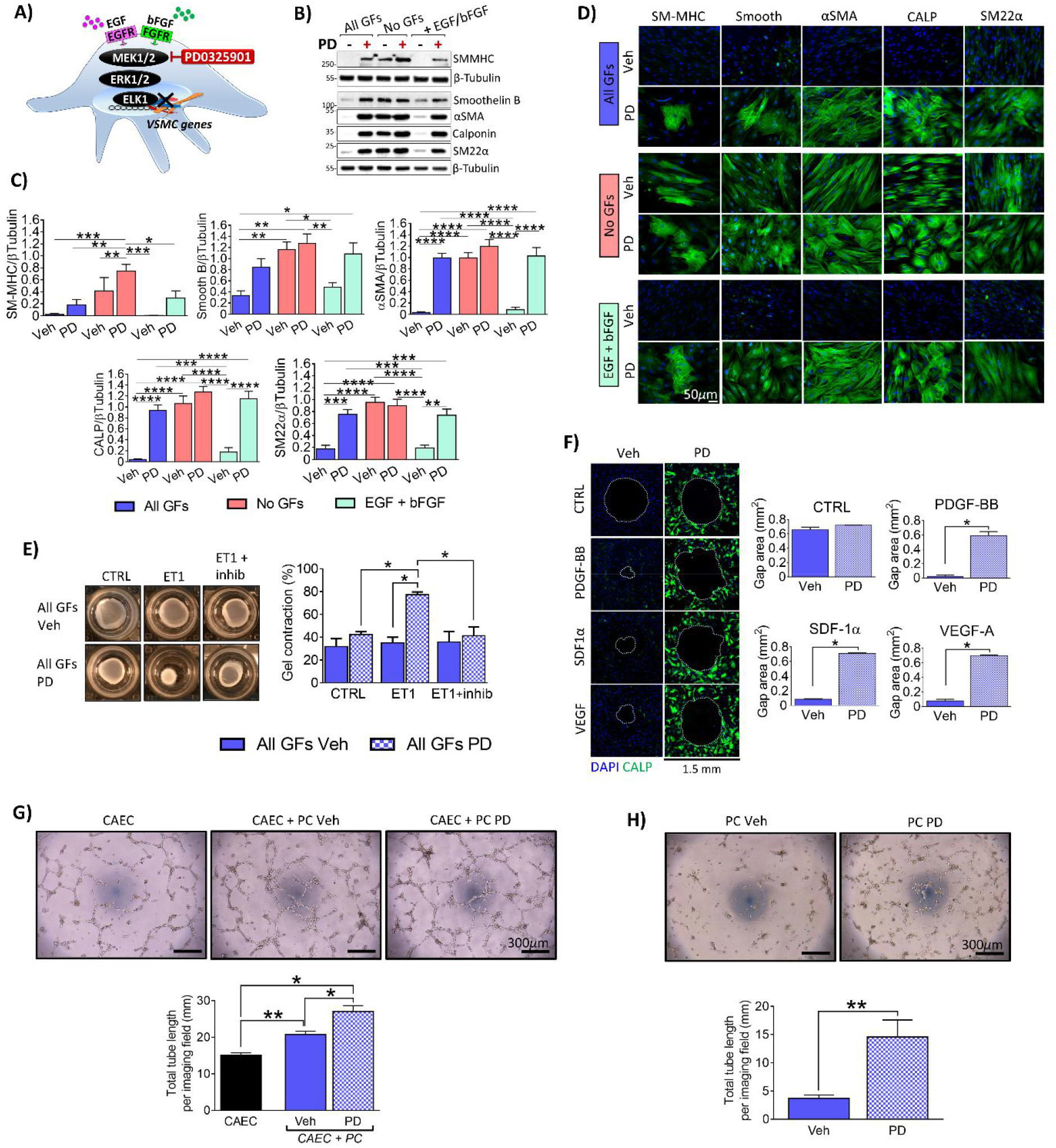
Inhibition of the MEK1/2 - ERK1/2 signalling induces the differentiation of human cardiac PCs into VSMC-like cells. **(A)** Schematic showing EGF and bFGF signalling. We have employed the MEK1/2 inhibitor PD0325901 to prevent ERK1/2 activation in PCs. Before using the cells for the functional assays, PCs were cultured for ten days with different media as indicated, and in the presence of PD0325901 (PD - 250 nM) or DMSO (Veh). (**B-C**) Protein expression analyses using western blotting. Representative blots from one patient and graphs reporting blots densitometry. N=5. Immunofluorescence images showing expression of VSMC markers. **(E) Contraction assay**. Cells were embedded in collagen gels, treated with a contraction inhibitor and stimulated with Endothelin-1 (ET1). Bar graphs indicate the % of gel contraction after 24h. N=4. **(F) Gap closure migration assay**. Migration time was 24h. Green fluorescence indicates Calponin, in blue DAPI. Bar graphs show the area of the final gap. N=4. **(G) 2D-Matrigel assay with human coronary artery ECs (CAEC) and PCs**. CAECs were used alone as control, or in co-cultures with cardiac PCs. N=3 to 5. **(H) 2D-Matrigel assay with PCs**. PCs alone were seeded on the top of Matrigel. N=6. In (G&H) bar graphs indicate the total tube length per imaging field. All values are means ± SEM. *P<0.05, **P<0.01, ***P<0.001, ****P<0.0001.

PCs differentiated with PD0325901 in the presence of all GFs contracted in response to ET-1 (**Figure 5E**), became unresponsive to pro-migratory stimuli (**Figure 5F**), and showed reduced proliferation, which are properties of contractile VSMCs (**Supplementary Figure 14**). Conversely, the cell cycle inhibitor hydroxyurea was unable to induce PC differentiation (**Supplementary Figure 15**).

PD0325901 upregulated contractile markers and inhibited proliferation also in control CASMCs (**Supplementary Figure 16**).

Interestingly, DPCs also showed a superior capacity to support angiogenesis in co-cultures with CAECs (**Figure 5G**), and gained the ability to assemble in tubular networks when seeded on the top of Matrigel (**Figure 5H**).

These findings indicate that ERK1/2 is a master controller of PC differentiation and that its inhibition can encourage both PCs and CASMCs towards a contractile phenotype.

### Human cardiac PCs differentiate into VSMC-like cells upon transplantation *in vivo*

We next investigated the potential of cardiac PCs to generate VSMC-like cells *in vivo*. Cardiac PCs were embedded in Matrigel containing either PD0325901 or vehicle and injected subcutaneously in C57BL6/J mice (**Figure 6A**). Plugs were harvested after seven days, and an antibody against the human Ku80-XRCC5 antigen was employed to recognize transplanted PCs (**Supplementary Figure 17A**). Matrigel was revealed using a secondary antibody anti-mouse (**Supplementary Figure 17B**). A small fraction of spindle-shaped PCs within the Matrigel stained positive for αSMA and CALP in the vehicle group (18.0% and 3.7% of total cells, respectively) these numbers being increased by PD0325901 (45% and 23% of total cells, respectively) (**Figure 6B&C**). Moreover, in the PD0325901 group, we could identify occasional SM-MHC-positive cells (**Figure 6D**). Intriguingly, the three-dimension (3D) reconstructions of 15 μm-thick Matrigel sections evidenced the presence of tubular-like structures only in PD0325901 plugs (**Figure 6E**). Immunohistochemistry quantification of CD45^pos^ cells documented a similar influx of immune/inflammatory cells within the vehicle-and PD0325901-plugs, ruling out host immune response was relevant for differences regarding implanted cells between the two groups (**Supplementary Figure 17C**).

**Figure 6.**
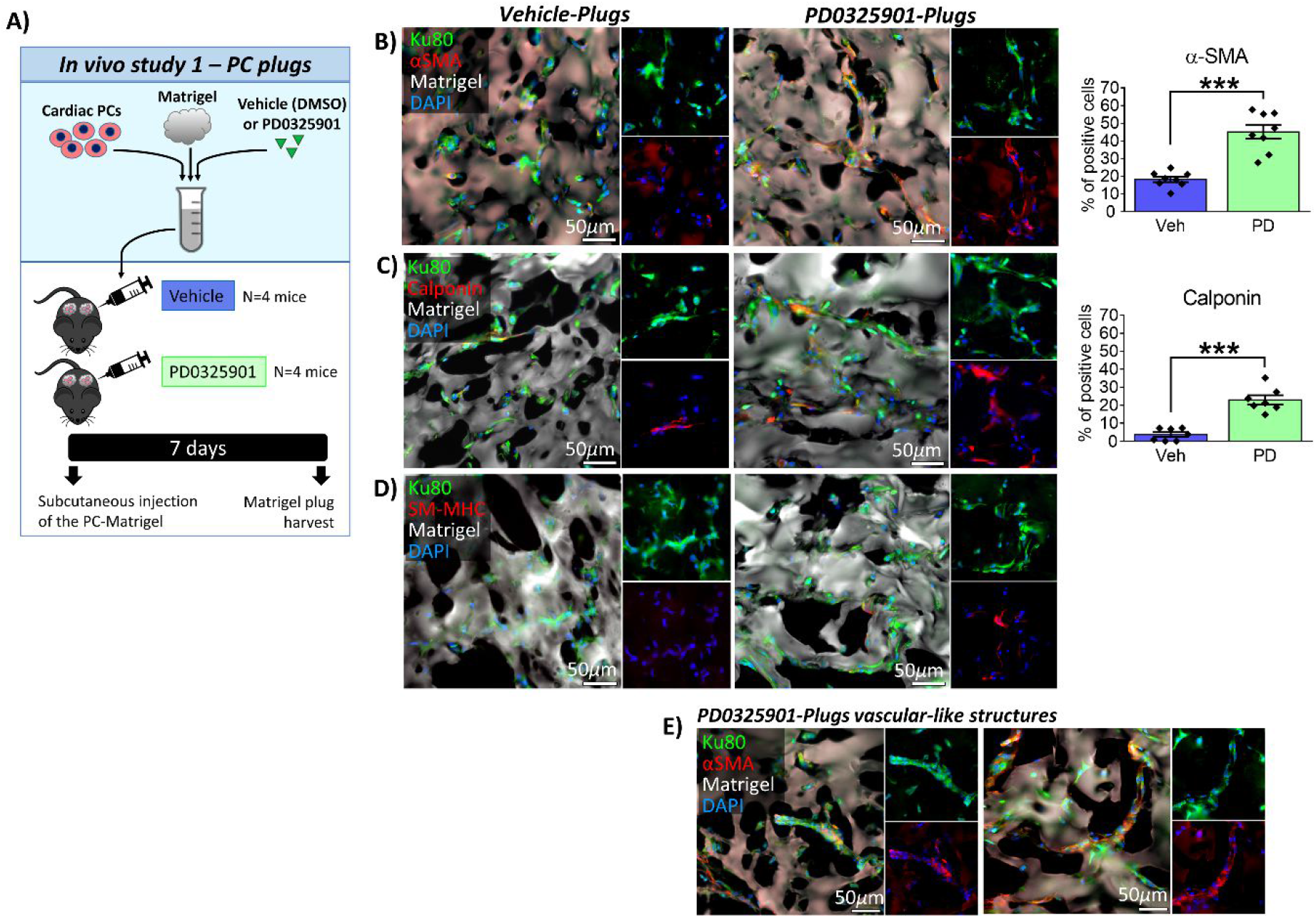
Human cardiac PCs differentiate into VSMC-like cells upon transplantation *in vivo*. **(A)** Experimental protocol of the Matrigel plug assay. **(B-D)** Immunofluorescence images of Vehicle (Veh) and PD0325901 (PD) Matrigel plugs showing that human cells express VSMC markers. Bar graphs report the percentage of human cells expressing α-SMA and Calponin. N=7-8/group. Values are means ± SEM. ***P<0.001. **(E)** Immunofluorescence images documenting the presence of α-SMA^pos^ vascular-*like* structures within the PD0325901-plugs.

### PD0325901 promotes myocardial arteriologenesis in healthy mice

We detected CD31^neg^ αSMA/CALP^neg^ CD34^pos^ PDGFRβ^pos^ cells around arterioles and capillaries in the mouse heart (**Supplementary Figure 18A&B**) and confirmed that *in vitro* cultures of murine PCs have a phenotype like the human PCs (**Supplementary Figure 18C&D**). Treatment of murine PCs with 250 nM PD0325901 increased the expression of contractile SM-proteins while halting cell proliferation (**Supplementary Figure 18E-G**).

A controlled, randomized study was then conducted in C57BL6/J mice receiving PD0325901 at 10 mg/kg/day or vehicle (dimethylsulfoxide - DMSO), orally for 14 days (**Figure 7A**). Baseline and endpoint indexes of LV function and dimension were similar between the experimental groups, as assessed by echocardiography (**Supplementary Table 6**). The inhibition of MEK1/2 activity in PD0325901-treated mice was confirmed by the absence of immunostaining for p-ERK1/2 in the heart (**Supplementary Figure 19A**) and by western blotting analysis of p-ERK1/2 in the liver (**Supplementary Figure 19B**). Moreover, safety was confirmed by verifying the treatment did not cause apoptosis in cardiomyocytes, vascular cells, and interstitial cells (terminal deoxynucleotidyl transferase dUTP nick end labelling - TUNEL assay) (**Supplementary Figure 19C**) or increase in plasmatic levels of cardiac troponin I (cTn-I) (**Supplementary Figure 19D**).

**Figure 7.**
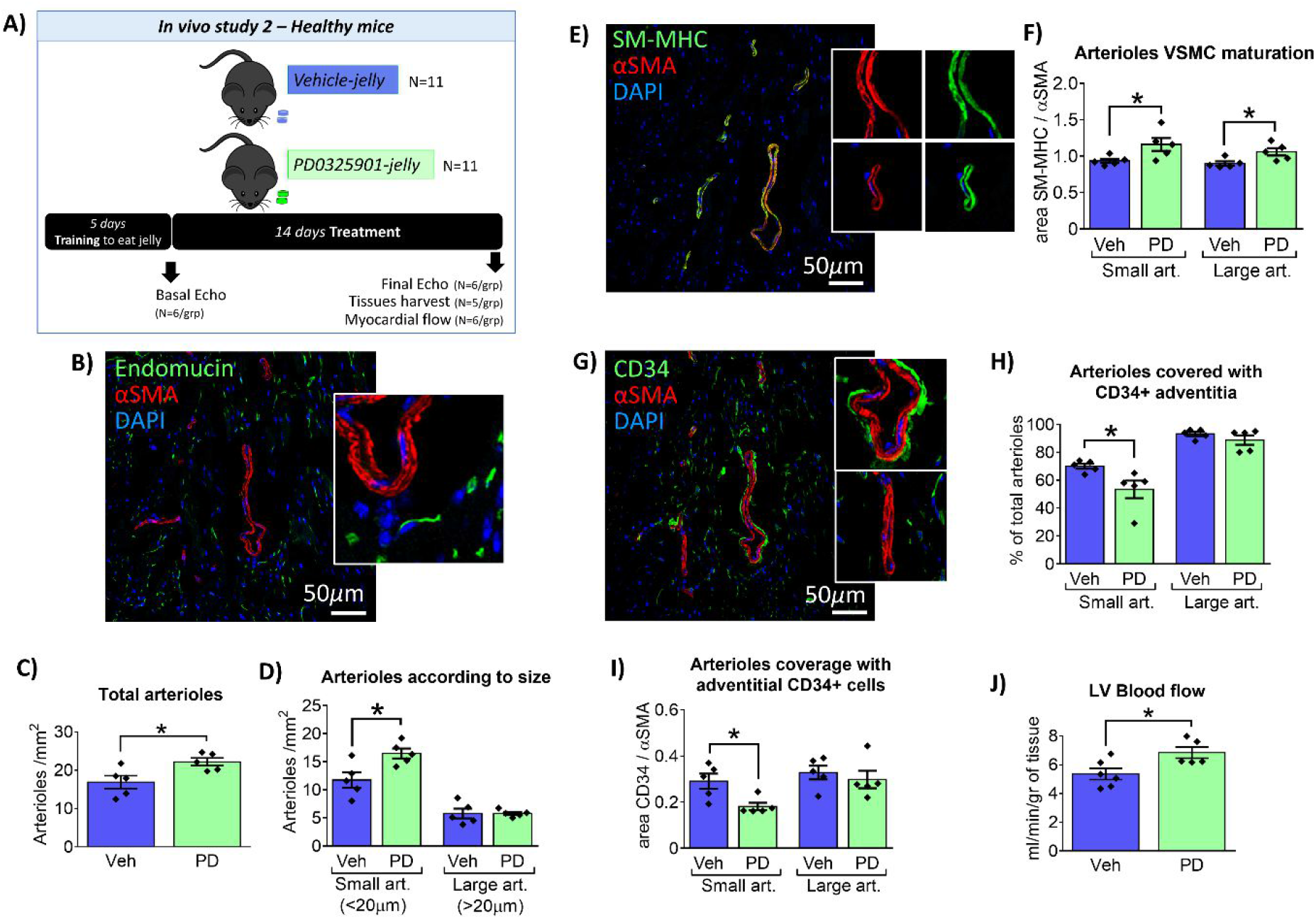
A 2-week treatment with PD0325901 promotes arteriologenesis and increased perfusion of the healthy mouse heart. **(A)** Experimental design. **(B-D)** Analysis of arteriole density. (B) Confocal immunofluorescence image of the LV showing examples of arterioles expressing αSMA but negative for Endomucin (marker of vein and capillary endothelium). Graphs showing the density of total arterioles (C) and arterioles categorized according to their lumen size (D). Data are expressed as the number of arterioles per mm^2^. **(E&F)** Analysis of arteriole maturation. (E) Confocal immunofluorescence image showing arterioles expressing αSMA and SM-MHC. (F) Graph reporting arterial VSMC maturation evaluated as the ratio between the areas occupied by SM-MHC and αSMA. **(G-I)** Analysis of perivascular CD34+ cells. (G) Confocal immunofluorescence image showing αSMA+ arterioles surrounded by adventitial CD34+ cells. (H) Graphs reporting arterioles surrounded by CD34+ cells expressed as a percentage of total arterioles. (I) Coverage of arterioles with CD34+ cells was evaluated as the ratio between the areas occupied by CD34 and αSMA. In (C-I) N=5/group. **(J)** Left ventricle blood flow. N=6 Veh, N=5 PD. For all graphs, individual values and means ± SEM. *p<0.05. *Veh*: vehicle. *PD*: PD0325901. *Art*: arteriole.

Histological analyses showed the density of arterioles was increased in the LV of PD0325901-treated mice compared with the vehicle group (**Figure 7B&C**), whereas cardiomyocyte cross-sectional area (CSA) and density were similar between the two groups (**Supplementary Figure 19E&F**). Interestingly, although the increase in vascular density could be attributed to small arterioles only (**Figure 7D**), the relative expression of mature vs. early contractile proteins in VSMCs of the vascular wall, was augmented by PD0325901 in the whole spectrum of arterioles (**Figure 7E&F**). Moreover, PD0325901 reduced the percentage of small arterioles covered with a CD34^pos^ layer (53% vs. 70% of total arterioles in the vehicle) (**Figure 7G&H**) as well as the ratio between the area occupied respectively by perivascular CD34^pos^ cells and αSMA^pos^ VSMCs (**Figure 7G&I**).

Venous vessels were recognized with the capillary- and vein-endothelium specific antigen Endomucin and excluded from the analysis (**Supplementary Figure 19G**). The PD0325901 treatment did not affect capillary density (**Supplementary Figure 19H**).

The observed structural changes were translated into an improvement of myocardial perfusion in the PD0325901 group (6.8 vs. 5.3 ml/min/g of LV tissue in vehicle) (**Figure 7J**).

### PD0325901 improves LV function and revascularization in a mouse MI model

Finally, a controlled, randomized study was conducted in C57BL6/J mice with MI. Three-days post-MI, mice were given PD0325901 at 10 mg/kg/day or vehicle (DMSO), orally for 14 days (**Figure 8A**). Echocardiography confirmed the proper induction of MI. Baseline indexes of LV function did not differ between groups.

**Figure 8.**
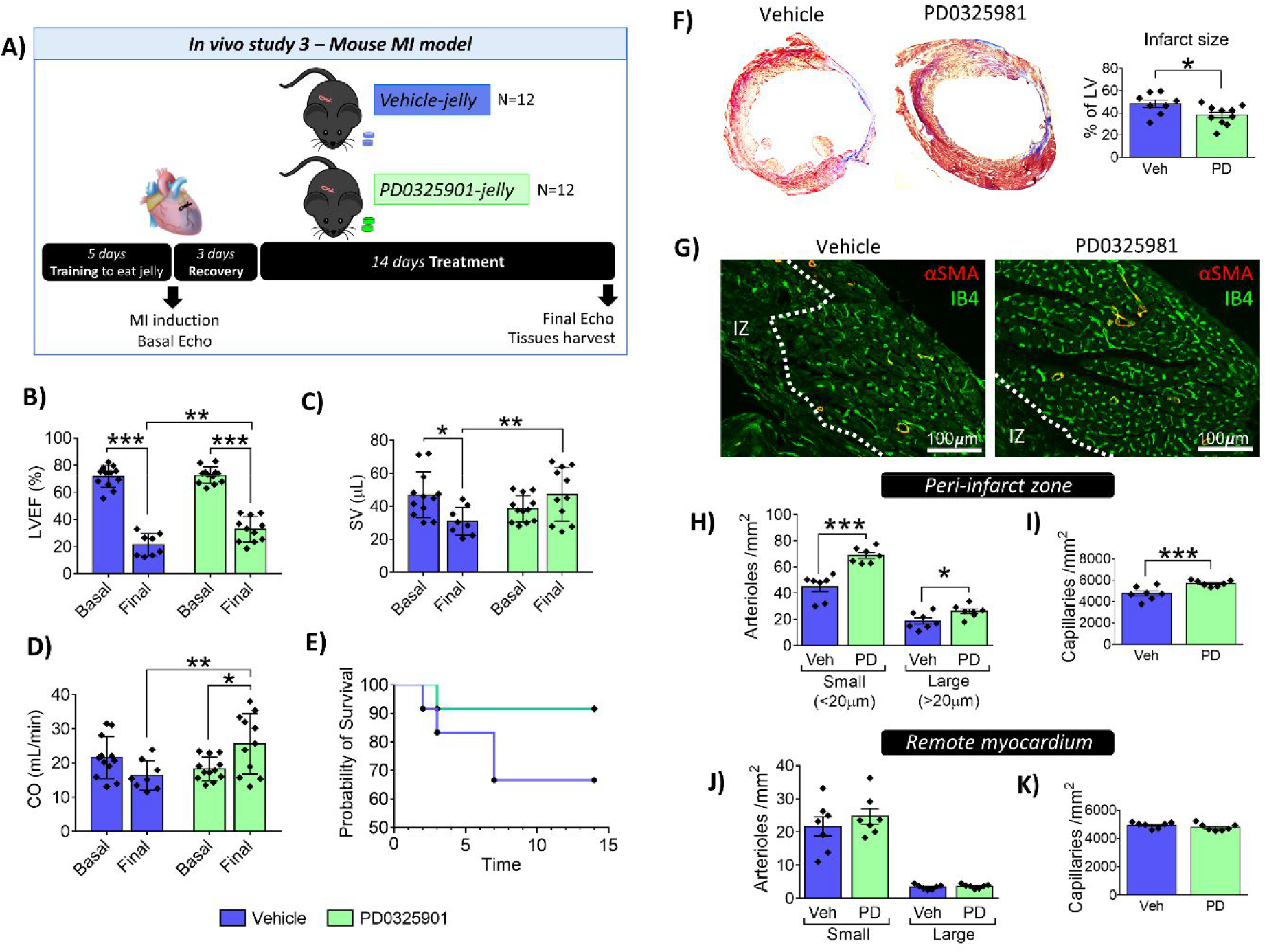
A 2-week treatment with PD0325901 improves LV function and vascularisation in a mouse MI model. **(A)** Experimental design. **(B-D)** Graphs showing basal and final echocardiography indexes. For Vehicle: N=12 basal, N=8 final; PD: N=12 basal, N=11 final. Individual values and means ± SD. **(E)** Graph reporting mice survival. **(F)** Representative images showing the Azan-Mallory staining of the LV and bar graphs indicating the quantification of the infarct size. N=8 Veh, N=10 PD. **(G)** Representative immunofluorescence images showing arterioles (αSMA, red fluorescence) and capillaries (Isolectin B4, green) in the peri-infarct myocardium of mice receiving vehicle or treatment. The dashed line defines the infarct zone (IZ). **(H-K)** Graphs reporting the quantification of arteriole and capillary density in the peri-infarct and remote LV myocardium of mice receiving vehicle or treatment. N=7 per group. In (F-K), individual values and means ± SEM. For all graphs, *p<0.05, **p<0.01, ***p<0.001 as indicated. Veh: vehicle. PD: PD0325901.

At the endpoint, PD0325901-treated mice showed reduced LV dilatation (**Supplementary Table 7**) and improved contractile function, as indicated by higher LV ejection fraction (LVEF, +11.4 percent points vs. vehicle), stroke volume (+52% vs. vehicle), and cardiac output (+56% vs. vehicle) (**Figure 8B-D**). Survival rate did not differ between groups (**Figure 8E**). However, the composite endpoint of survival and LVEF above 30% was significantly better in the PD0325901 group (Fisher exact test: p=0.0272) with an improved relative risk of 2.20 (95% CI = 1.23 to 4.80).

Histological examination evidenced that mice in the PD0325901-group had a smaller infarct size (38 vs. 48% of the LV in controls) (**Figure 8F**). Moreover, in the peri-infarct zone, PD0325901 induced a significant increase in the density of small (69 vs. 45 art/mm^2^ in controls) and large arterioles (26 vs. 19 art/mm^2^ in controls) (**Figure 8G&H**). Similarly, capillary density was higher in the PD0325901 group (5,700 vs. 4,700 cap/mm^2^ in controls) (**Figure 8G&I**). Conversely, PD0325901 did not modify vascularization in the remote myocardium (**Figure 8J&K**).

## Discussion

Here, we provide a new mechanistic understanding of cardiac PC potential in vascular remodelling. In the heart, PCs may represent an incremental cellular reservoir for fuelling arteriologenesis in conditions of ischemic injury. Importantly, we show that myocardial vascularization can be pharmacologically modulated *in vivo* using the selective MEK inhibitor PD0325901, although differences were observed between the normoperfused and ischemic murine heart. In the former, PD0325901 administration induced a generalized increase in small arterioles without affecting capillary density, whereas, when started at the early recovery stage from acute non-reperfused MI, the inhibitor potentiated both small and large arterioles selectively within the peri-infarct zone and also incited capillarization. Thus, the approach could have incremental therapeutic activities in specific areas of the heart depending on the underlying state of the coronary circulation and the occurrence of regional ischemia. Although additional investigation is needed, our findings raise the intriguing possibility of manipulating mural cells to generate a robust microvasculature in the adult heart.

We observed that DPCs became able to assemble in vascular-like tubes in an *in vitro* angiogenesis assay, along with forming more complex tubular networks in cooperation with ECs. Transcriptomic analysis further showed that DPCs acquired a pro-angiogenic profile whose most relevant elements are the downregulation of potent disruptors of angiogenesis, namely *ANGPT2, TIE1* and *SERPINF1*, paralleled by the induction of pro-angiogenic factors *LEP* and *PDGFB*. ANGPT2 antagonizes the pro-angiogenic ANGPT1/Tie2 signalling and was reported to be upregulated in the ischemic heart in mice and to cause abnormal vascular remodelling.^18^ Leptin is reportedly expressed by perivascular PDGFRβ^pos^ cells,^19^ and contributes to the pro-angiogenic activity of transplanted PCs in a mouse model of limb ischemia.^20^ Potentiation of angiogenesis and arteriologenesis is crucial in preserving cardiomyocyte viability in the area at risk, which may account for the finding of a smaller infarct scar documented in the PD0325901-treated group at sacrifice, 2 week after MI induction.

We discovered that EGF and bFGF play a primary role in the preservation of proliferative, undifferentiated cultures of cardiac PCs, through the activation of the ERK1/2 signalling. In contrast, upon GF depletion, PCs undertake a phenotypic switch to VSMCs. *In vitro*, differentiated PCs become stationary in migration assays, express elastin and contractile fibres typical of VSMCs, and also gain functional contractile features, as shown by the calcium inflow and ability to contract following ET-1 stimulation. It is known that bFGF inhibits transforming growth factor-beta 1 (TGFβ1)-driven activation of SM contractile gene expression in PCs and VSMCs.^21-23^ However, this is the first demonstration that cardiac PCs are intrinsically committed to a SM phenotypic switch upon removal of the inhibitory GFs. Both EGF and bFGF are upregulated in the acutely infarcted heart and unpublished data confirmed the elevation of the two GFs in peripheral blood of patients with recent MI (Katare et al, 2020). It is therefore plausible that PCs are initially restrained from differentiation into VSMC-like cells following an acute myocardial injury, participating in arteriologenesis upon GF downregulation during later recovery stages. A pharmacological approach impacting on the EGF-bFGF-ERK1/2 signalling could be a viable means to promote prompter arteriologenesis of the heart.

Several lines of evidence support this possibility. Administration of PD0325901, a targeted MEK1/2 inhibitor, evoked a SM phenotypic switch in human cardiac PCs cultured in the presence of GFs. This change was validated using RNA-Seq, which showed an upregulation of several clusters of genes associated with SM contraction and function in DPCs. Despite the whole transcriptome profiling indicated that DPCs were different from CASMCs, the upregulation of SM pathways in PCs was crucial to confer the cells with both phenotypic and functional properties of contractile VSMCs, as demonstrated by *in vitro* studies. We cannot exclude that this transformation is further magnified *in vivo*. We also observed that the MEKi induced the expression of mature contractile VSMC markers in proliferating CASMCs *in vitro*. These data support the intriguing possibility that different mural cell types can participate in muscular remodelling of the myocardial vasculature. PD0325901 was previously used to induce the complete differentiation of immature SMCs derived from human pluripotent stem cells.^24^ Interestingly, in our system, the effect of PD0325901 was faster and more potent than that observed by withdrawing GFs from the culture environment. Moreover, we have demonstrated the pro-arteriologenesis effect of PD0325901 *in vivo* using Matrigel plug-implanted PCs. In this assay, the acquisition of the late-stage differentiation marker SM-MHC was low, possibly because of the dispersion of the MEKi in the surrounding tissues. Therefore, we designed new experiments using daily, oral administration of the drug, to verify cardiac safety and efficacy in normoperfused and ischemic murine hearts. In both models, we confirmed that the treatment promotes myocardial vascularization. The increase in arteriologenesis might be attributed, at least in part, to the differentiation of PC-covered vascular structures into mature αSMA^pos^ arterioles. Additionally, angiogenic capillaries in the infarct border zone could be muscularized through the recruitment and differentiation of synthetic VSMCs and PCs. The contribution of cell proliferation cannot be excluded, although GF depletion and ERK1/2 inhibition exerted anti-proliferative activity on PCs.

### Clinical relevance and study limitations

This study documents that myocardial PCs are endowed with intrinsic vascular plasticity, which can be pharmacologically reprogrammed to encourage arteriologenesis and angiogenesis. Short-duration treatment with PD0325901 may aid the recovery from MI through enhanced vascularization of the area at risk. It is also worth to report that PD0325901 showed efficacy in reducing neointima formation in a mouse model of arterial stenosis,^25^ which increases the potential cardiac benefits of this class of compounds. Additional studies are warranted to demonstrate the benefit of repeated MEKi administration in ischemic heart failure and diseases characterized by arteriole regression, such as diabetic cardiomyopathy. In this respect, chronic treatment may require lowering the therapeutic dosage to avoid cardiotoxicity.^26, 27^ This was not a case in our study, where a short-duration treatment with a specific MEK1/2 inhibitor was applied.

Last, although this study focused on the analysis of vascular mural cells, we are fully aware that the direct beneficial effect of the MEKi on other cardiac cell types cannot be excluded and merits further investigation. This does not diminish the translational significance of presented findings, rather strengthens the concept that the heart has an intrinsic reparative potential worth to be aided after an acute ischemic injury.

## Authors contribution

**EA:** conception and design, data collection, analysis and interpretation, presentation and assembling of data; manuscript writing

**RK:** data collection, analysis, and interpretation

**ACT:** data collection and analysis

**AC:** data collection, analysis, and interpretation

**DS:** data collection and analysis

**MM:** data analysis

**MC:** recruitment of patients and provision of human samples

**PM:** conception and design, data analysis and interpretation, manuscript writing, provision of funding All Authors revised the article for relevant intellectual content and approved the final version for publication.

## Acknowledgements

We wish to acknowledge the Wolfson Bioimaging Facility for the access to confocal microscopes and expert technical advice; Professor Andrew Herman of the Flow Cytometry Facility for his help with the single-cell sorting, all from the University of Bristol. We also acknowledge the University of Edinburgh Bioresearch&Veterinary Services at Little France Facility for the support on the *in vivo* Matrigel experiment. EA is a postdoctoral fellow supported by the British Heart Foundation Centre for Vascular Regeneration. MC is a British Heart Foundation Professor of Cardiac Surgery.

## Data availability

The data underlying this article will be shared on reasonable request to the corresponding authors.

---

**Supplementary Figure 1.**
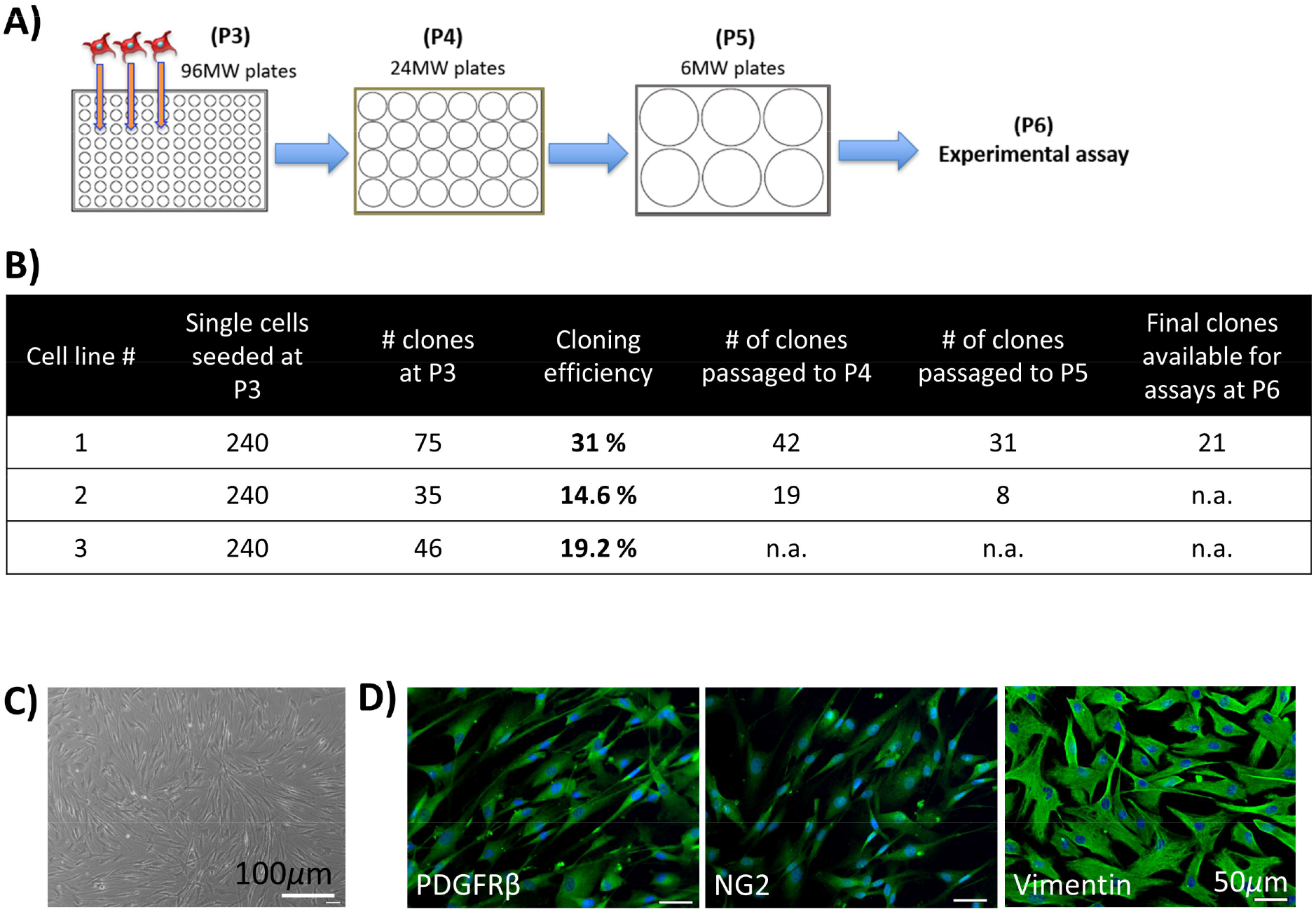
Human cardiac PCs are a clonogenic population. **(A)** Cartoon showing the experimental protocol for the single cell cloning. **(B)** Table summarising the results of the cloning assay with N=3 PCs. **(C)** Brightfield microphotograph of an expanded clone of PCs at passage 5 showing the characteristic spindle shape. **(D)** Representative immunofluorescence images showing the antigenic profile of a clone of PCs at passage 5 of culture. N=4 clones assayed, all belonging to cell line #1. **P** = passage; **n**.**a**. data not available (experiment not performed).

**Supplementary Figure 2.**
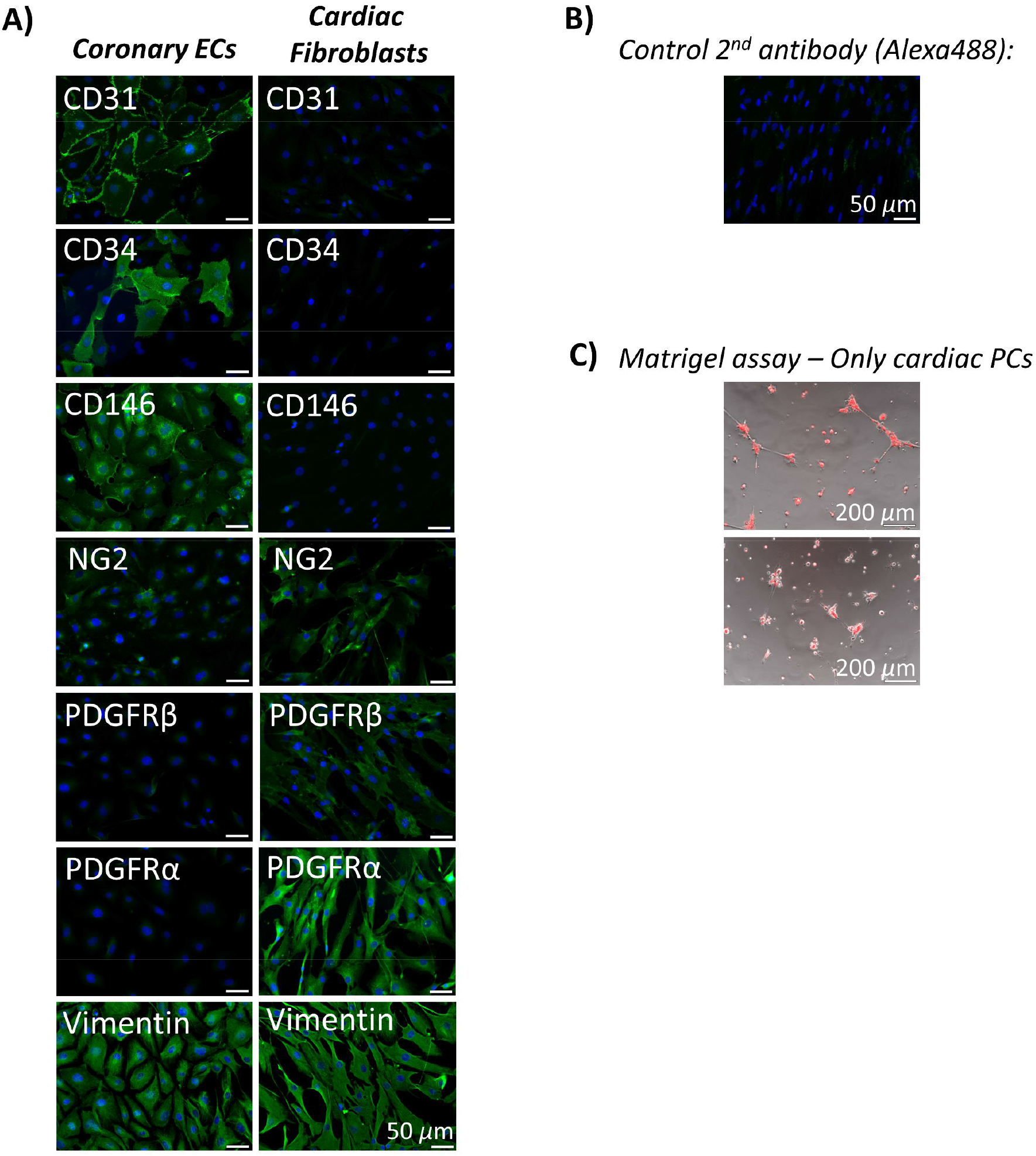
Controls for studies with human cardiac PCs. **(A)** Immunofluorescence microphotographs showing expression of the indicated markers in cardiac endothelial cells and fibroblasts used either as positive or negative control as appropriate. **(B)** Control human cardiac PCs stained only with secondary antibodies Alexa-488 conjugated to evidence the specificity of the antibodies and the low autofluorescence background of the cells. **(C)** 2D-Matrigel assay: control with only cardiac PCs labelled with the red fluorescent tracker dil. Pericytes alone fail to assemble in stable tubular networks.

**Supplementary Figure 3.**
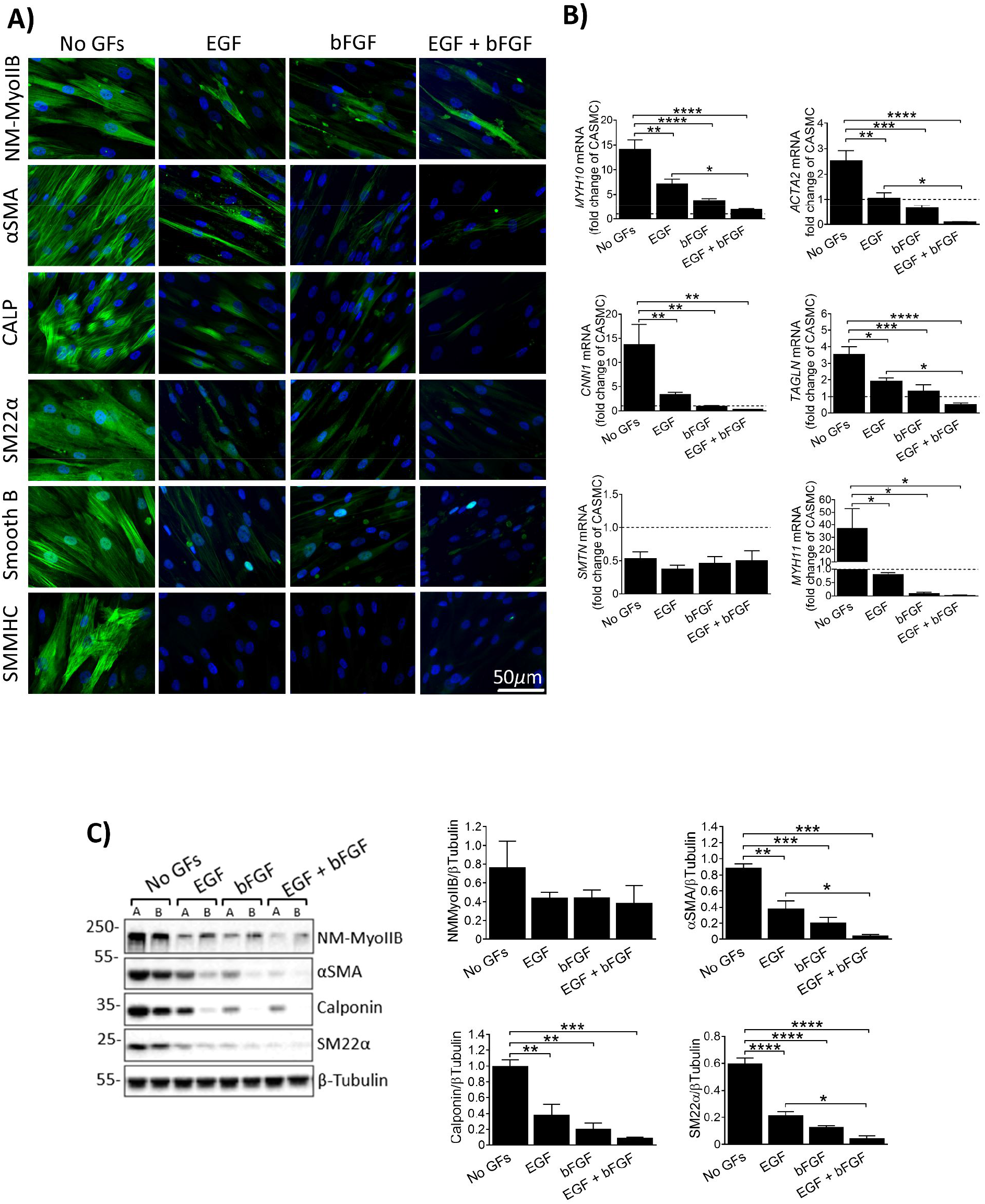
EGF and bFGF control the expression of contractile VSMC genes in human cardiac PCs. **(A)** Immunofluorescence images showing the expression of VSMC markers by PCs. Cells were cultured for 10 days. **(B)** Transcriptional analysis of VSMC genes in PCs and differentiated cells. Dashed line indicates coronary artery SMCs (CASMCs) used as reference cell line. Values are expressed as fold change of CASMCs. **(C)** Western blot analysis of VSMC proteins. A and B indicate cells from different donors. N=5 per group. Values are expressed as means ± SEM. * P<0.05, ** P<0.01, *** P<0.001, **** P<0.0001.

**Supplementary Figure 4.**
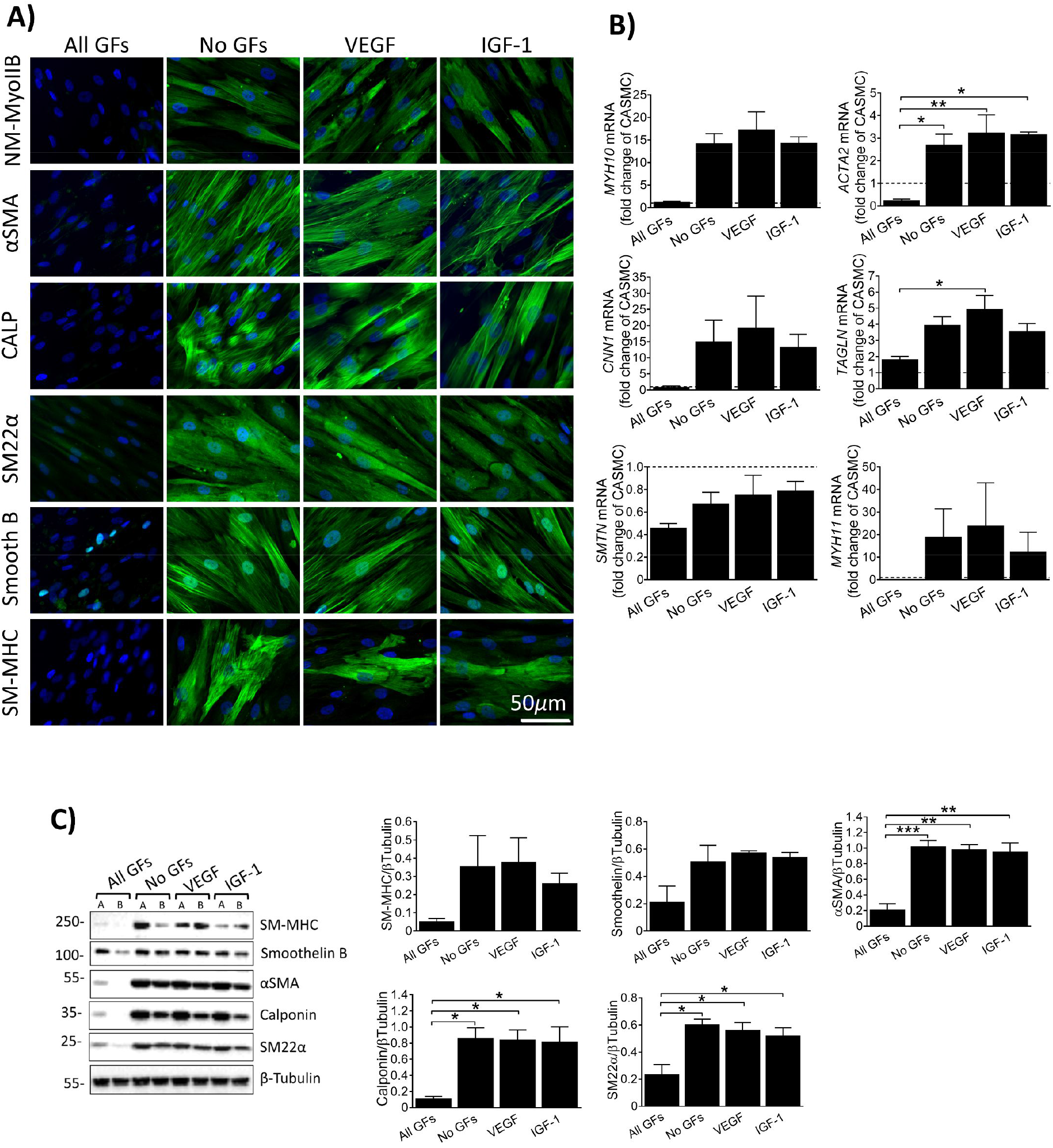
VEGF and IGF-1 promote the differentiation of human PC into VSMCs. Cells were cultured for 10 days with respective media. **(A)** Immunofluorescence images showing expression of VSMC markers by PCs and derived VSMC. **(B)** Transcriptional analysis of VSMC genes in PCs and differentiated cells. Dashed line indicates coronary artery SMCs (CASMCs) used as reference cell line. Values are expressed as fold change of CASMCs. **(C)** Western blot analysis of VSMC proteins. A and B indicate cells from different donors. N=3 per group. Values are expressed as means ± SEM. * P<0.05, ** P<0.01, *** P<0.001.

**Supplementary Figure 5.**
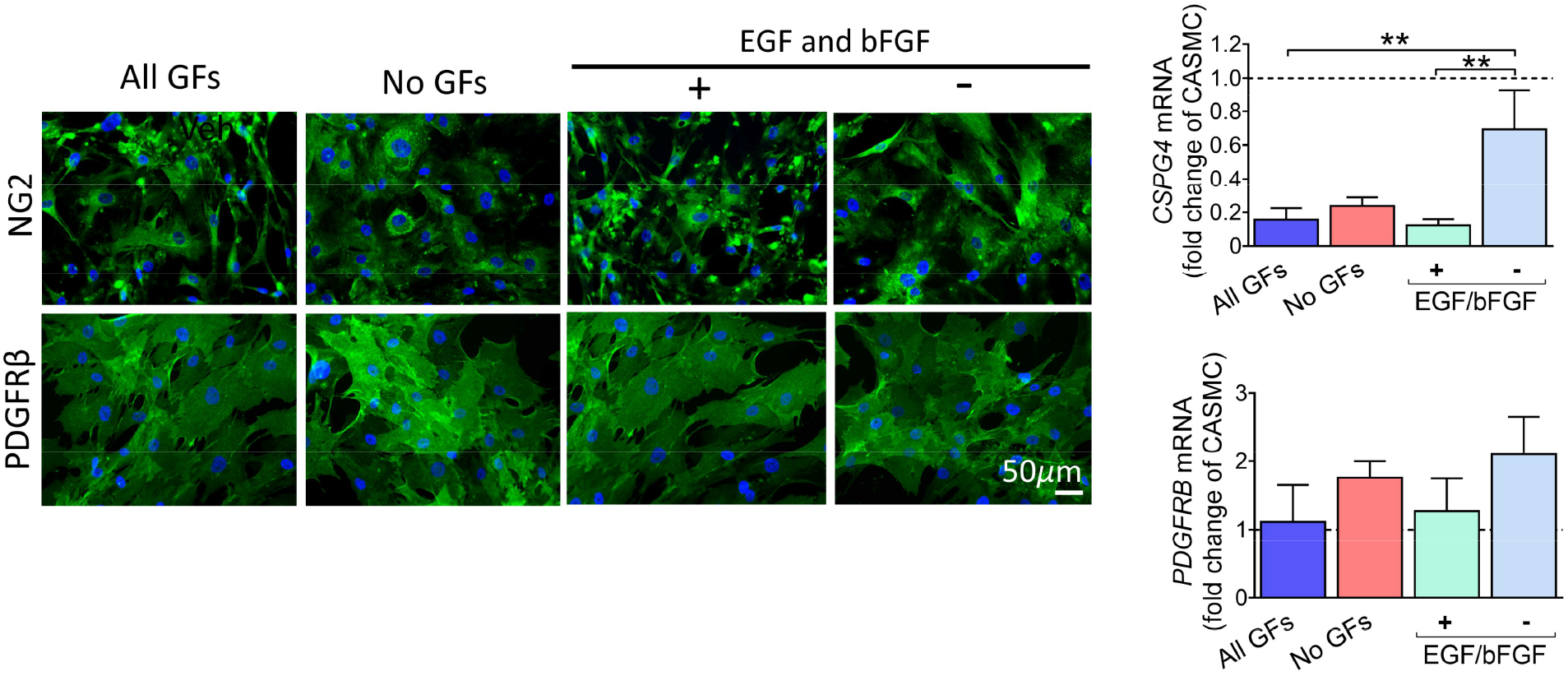
Differentiated human cardiac PCs retain the expression of common pericyte/VSMC genes. **(A)** Immunofluorescence images showing expression of NG2 and PDGFRβ by PCs and derived VSMCs. **(B)** Transcriptional analysis. Dashed line indicates coronary artery SMCs (CASMCs) used as reference cell line. Values are expressed as fold change of CASMCs. N=5 per group. ** P<0.01.

**Supplementary Figure 6.**
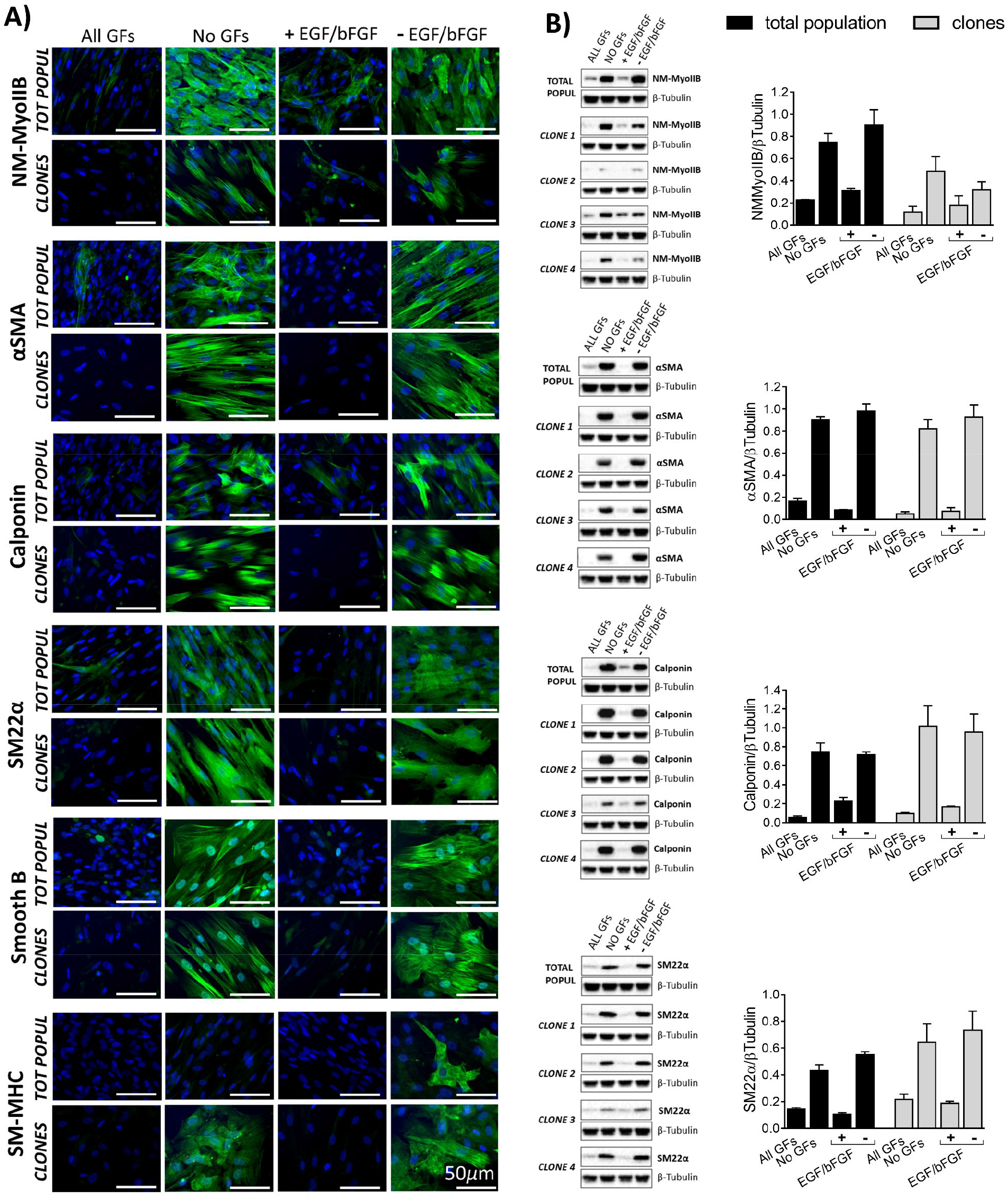
Clones of human PCs differentiate into VSMC in the absence of EGF and bFGF, similar to the total population. The VSMC differentiation capacity of clones (N=4) of one PC line was compared with that of the original population (N=3 different aliquots). No significant difference was observed in the expression of the proteins between the total population and its clones for each experimental condition. Cells were cultured for 10 days with respective media. **(A)** Immunofluorescence images showing VSMC protein expression. **(B)** Western blots and bar graphs showing blots quantification. For the total population, a representative blot (one of 3 aliquots of cells used) is shown. For the 4 clones, all blots are shown. N=3 total population, N=4 clones. Values are means ± SEM.

**Supplementary Figure 7.**
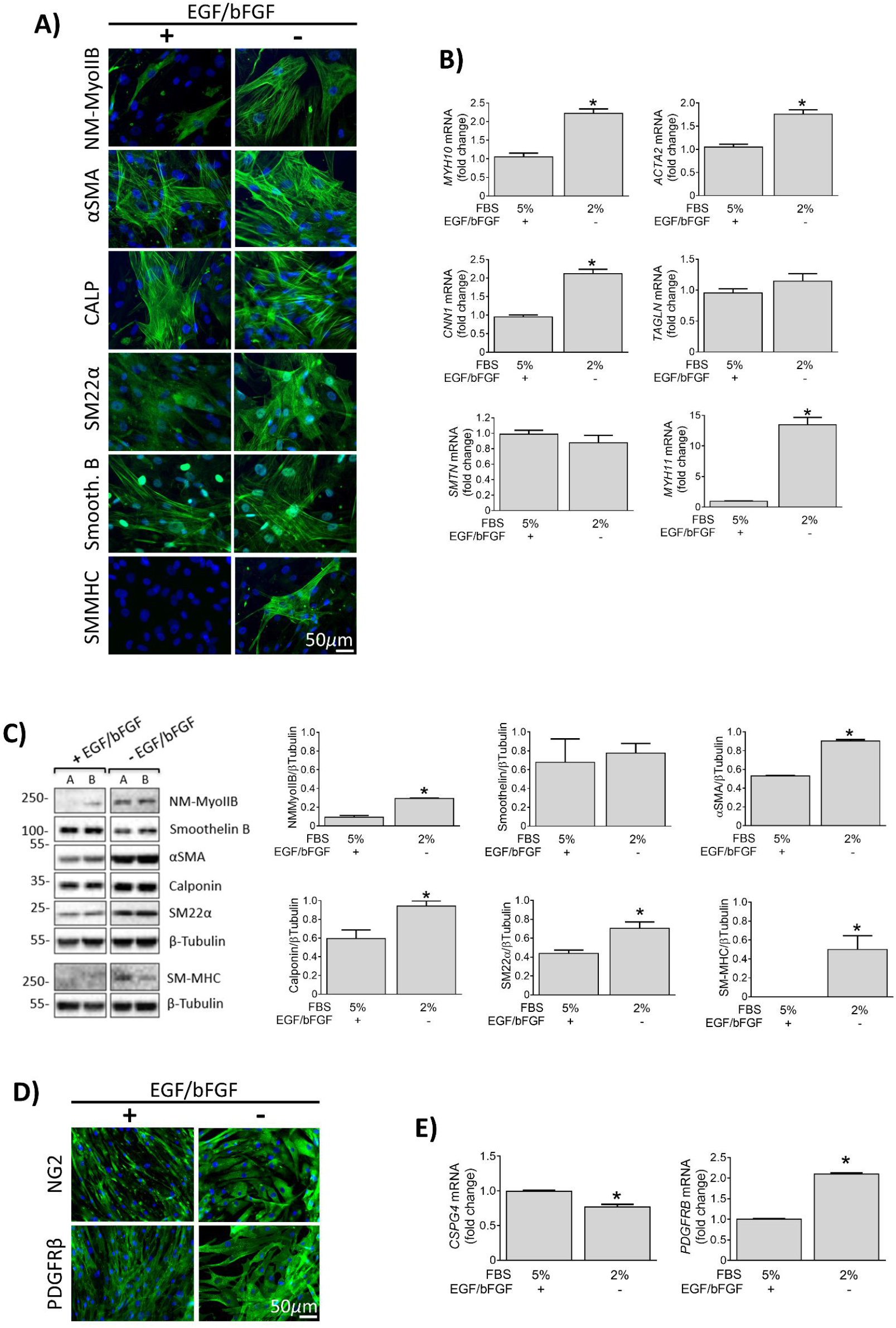
Human coronary artery SMCs (CASMCs) were used as a positive control for experiments with cardiac PCs. Cells were cultured for 10 days with their growth media in the presence or absence of GFs as indicated. Depletion of GFs and reduction of the % of FBS were applied to stimulate the differentiation of cells into a more contractile phenotype. **(A)** Immunofluorescence images showing expression of VSMC markers. **(B)** Transcriptional analysis of VSMC genes in control and differentiated cells. Values are expressed as fold change of control CASMCs. **(C)** Western blot analysis of VSMC proteins expression. A and B indicate two donors. **(D)** Immunofluorescence images showing expression of common pericyte/VSMC markers. **(E)** Transcriptional analysis of pericyte/VSMC genes in control and differentiated cells. Values are expressed as fold change of control CASMCs. In (B-C), N=4 per group (2 donors, 2 independent replicates each). In (D-E) N=3. All values are expressed as means ± SEM. * P<0.05 vs control cells.

**Supplementary Figure 8.**
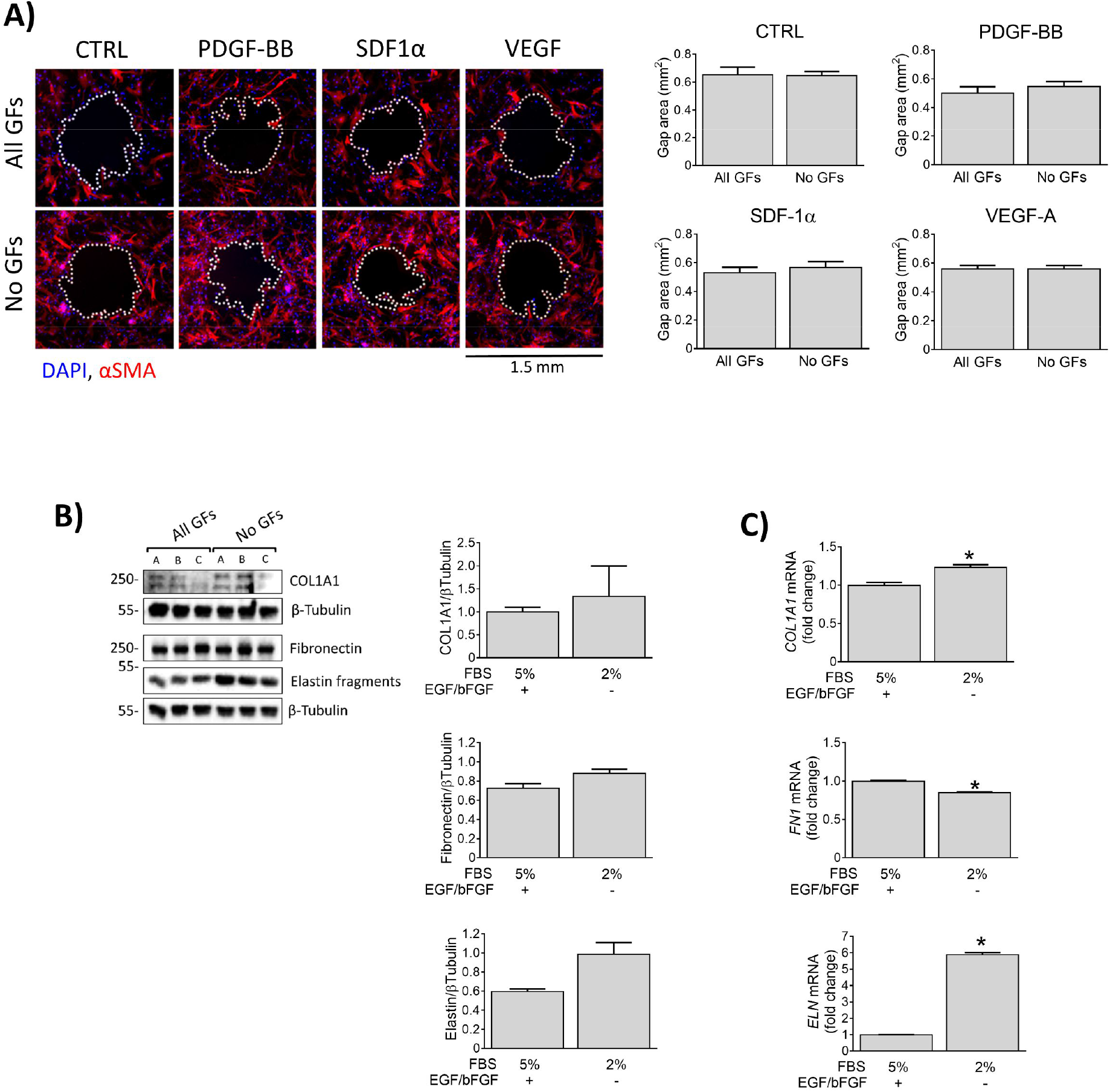
Human coronary artery SMCs (CASMCs) were used as a reference control in functional assays. **(A)** Gap closure migration assay. Migration time is 24h. Cells were differentiated for 10 days and then used in the migration assay. N=3. **(B-C) Expression of extracellular matrix proteins**. (C) Blots and respective quantification showing expression of Collagen I, Elastin and Fibronectin in control and differentiated CASMCs. N=3 per group. A, B, C indicate cells from different donors. (C) Transcriptional analysis of ECM markers. N=3. All values are displayed as means ± SEM.

**Supplementary Figure 9.**
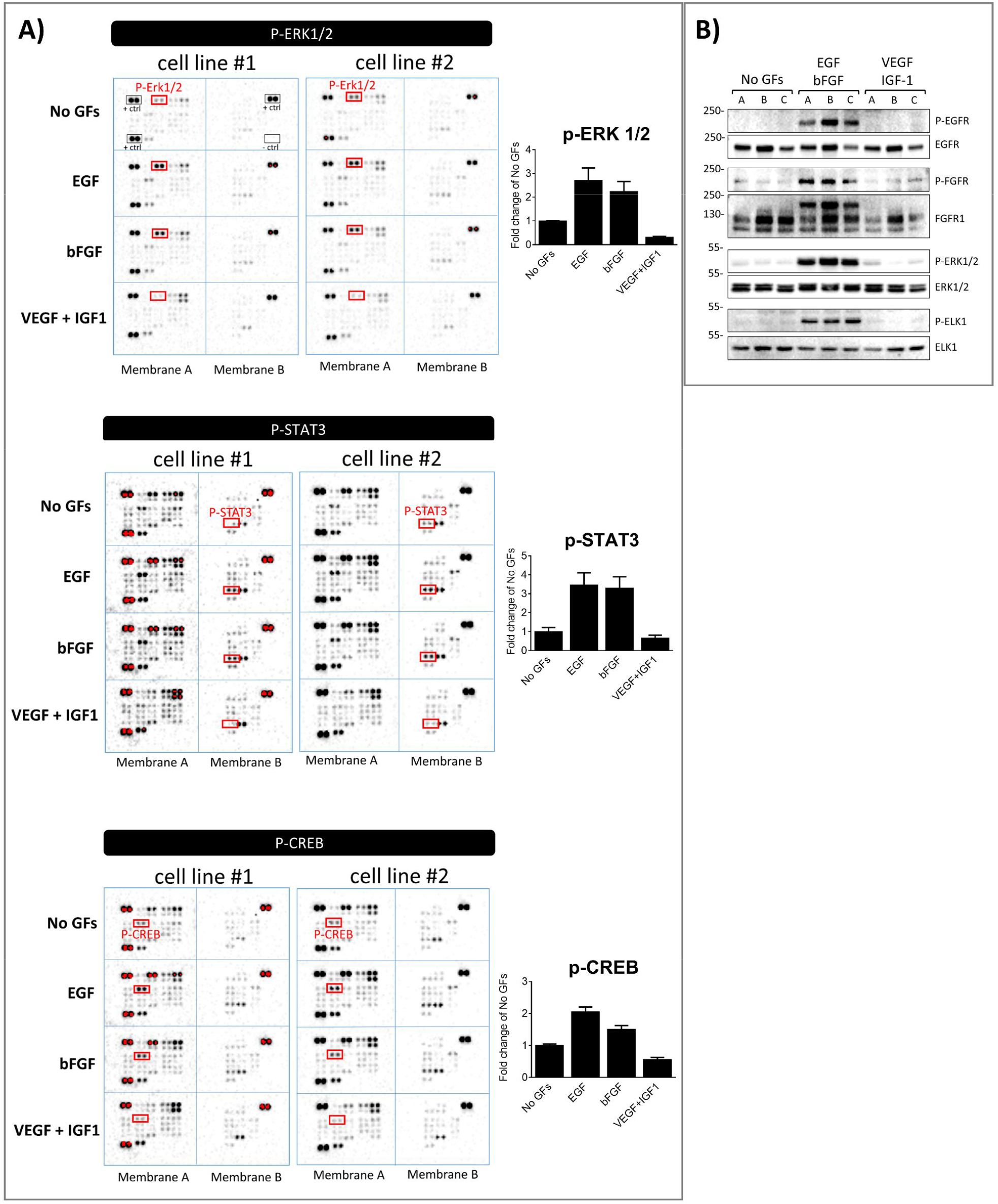
**A) Phospho-Kinase array with human cardiac PCs.** For a quick screening of the intracellular signalling activated by EGF and bFGF but not by VEGF and IGF1 in cardiac PCs, a human phospho-Kinase protein array was performed (N=2 PCs). The array allowed the detection of the phosphorylation of 43 kinases (R&D ARY003B). We report the main results of the screening, the membranes and densitometry graphs. **(B)** A subsequent western blot analysis confirmed the activation of EGFR-FGFR-ERK1/2 signalling by EGF and bFGF in cardiac PCs. N=3. A, B, C are different donors.

**Supplementary Figure 10.**
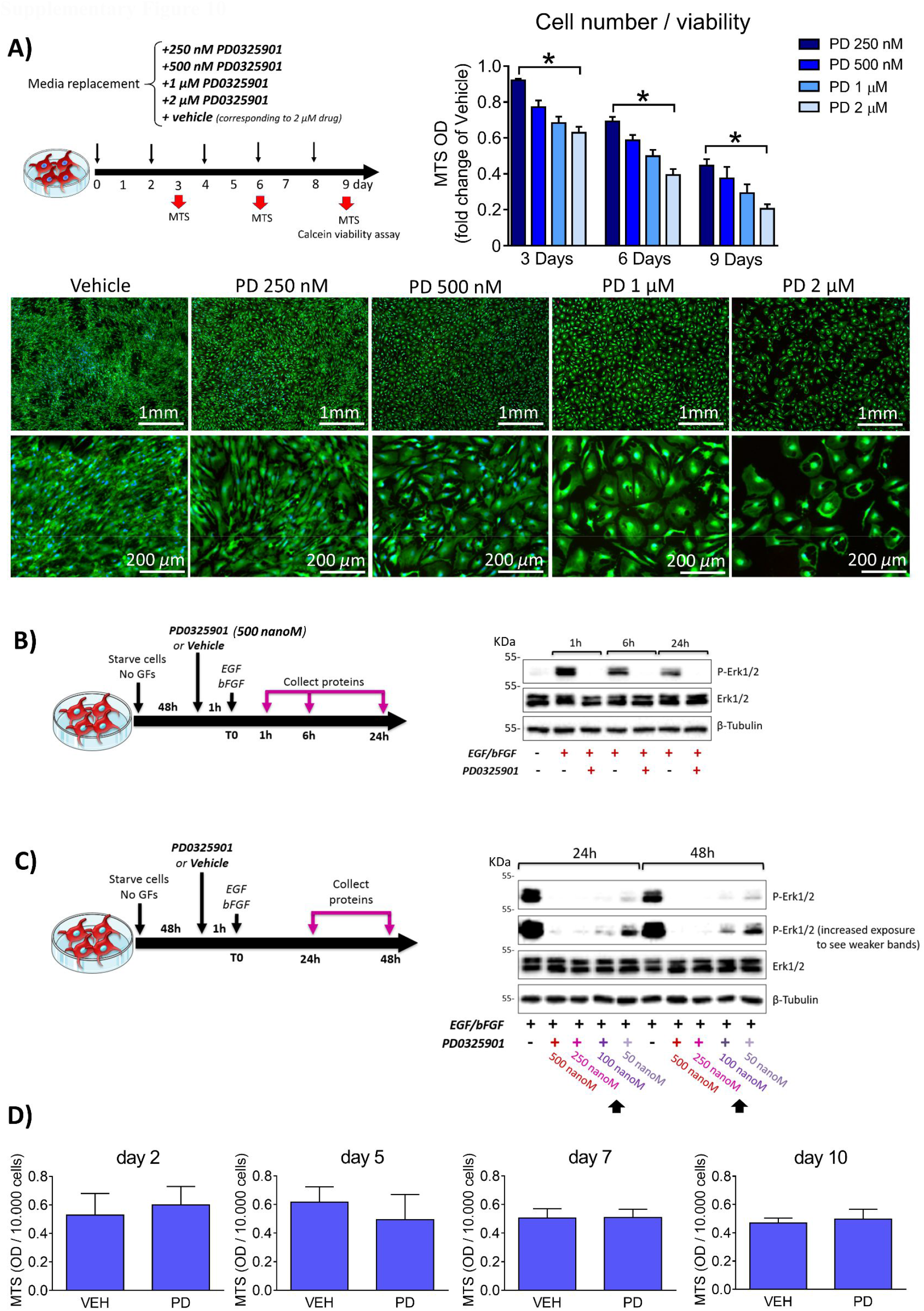
Dose-response study to assess PD0325901 efficacy and cytotoxicity in human cardiac PCs. **(A)** Four increasing doses of PD0325901 were tested in PCs up to 9 days. The DMSO corresponding to the higher concentration of the drug was used as control. Bar graphs report the results of the MTS assay. Results for each sample is reported as fold change of the respective vehicle. N=3 per group (4 replicates per each sample). Values are means ± SEM. **\****p*<0.05. The Calcein-AM/EthDIII viability assay (fluorescence microscopy images) was performed only at the day 9. Viable cells have a green cytoplasm, while dead cells are identified by red nuclei. HOECHST labels nuclei in blue (N=2 CPs). **(B)** The drug used at a concentration of 500 nanoM was able to prevent ERK1/2 phosphorylation in PCs (N=1) up to 24h. **(C)** Dose-response study to determine the minimum dose of the drug able to prevent ERK1/2 phosphorylation up to 48h. In (B&C), cells were initially starved for 48h and pre-treated with the drug/vehicle for 1h before stimulation with EGF and bFGF. **(D)** MTS assay showed that there is no significant difference in viability/metabolic activity between cells treated with the drug and its vehicle up to 10 days. MTS optical density (OD) values were normalised for the number of cells. N=3 per group. Values are means ± SEM. **Veh**: vehicle. **PD**: PD0325901.

**Supplementary Figure 11.**
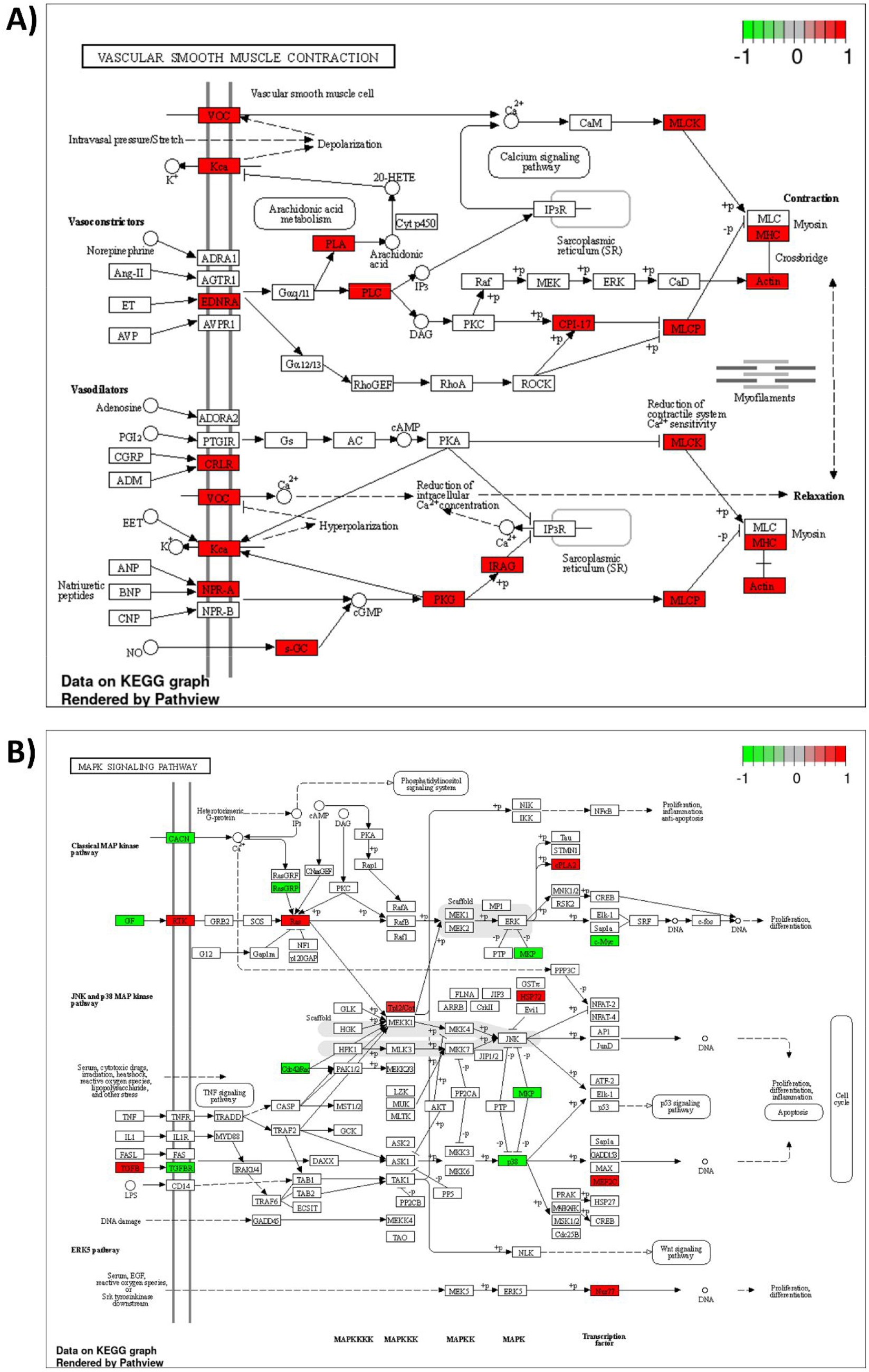
RNA-Seq of naïve and differentiated pericytes - effects of PD0325901. Schematic view of the main differentially regulated hits emerging from the pathways “*Vascular smooth muscle contraction*” **(A)** and “*MAPK signalling*” **(B)**.

**Supplementary Figure 12.**
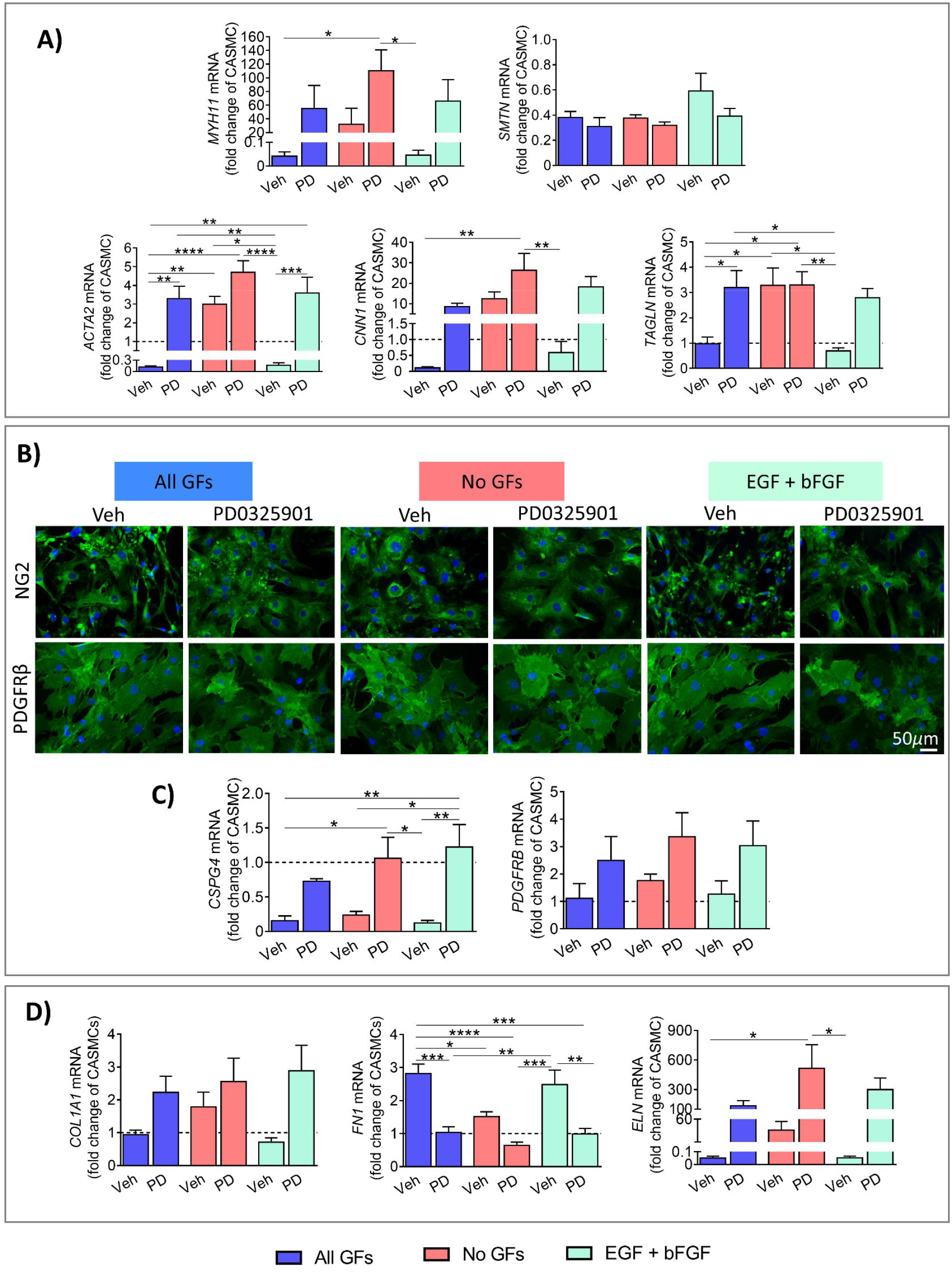
Expressional analyses in human cardiac PCs treated with PD0325901. PCs were cultured either with *All GFs* or with *No GFs* or with only *EGF + bFGF* for 10 days, in the presence of the MEK inhibitor (250 nanoM) or its vehicle DMSO, with complete media change every 48h. **(A)** VSMC gene expression analysis. The dashed line indicates coronary artery SMCs (CASMCs) employed as reference. **(B)** Representative immunofluorescence images showing common pericyte/VSMC markers at the end of the protocol. **(C-D)** Pericyte/VSMC (C) and ECM (D) gene expression analysis. The dashed line indicates coronary artery SMCs (CASMCs) employed as reference. Values are expressed as fold change of CASMCs. Values are means ± SEM. * *P*<0.05, ** *P*<0.01, *** *P*<0.001, **** *P*<0.0001. **Veh**: vehicle. **PD**: PD0325901.

**Supplementary Figure 13.**
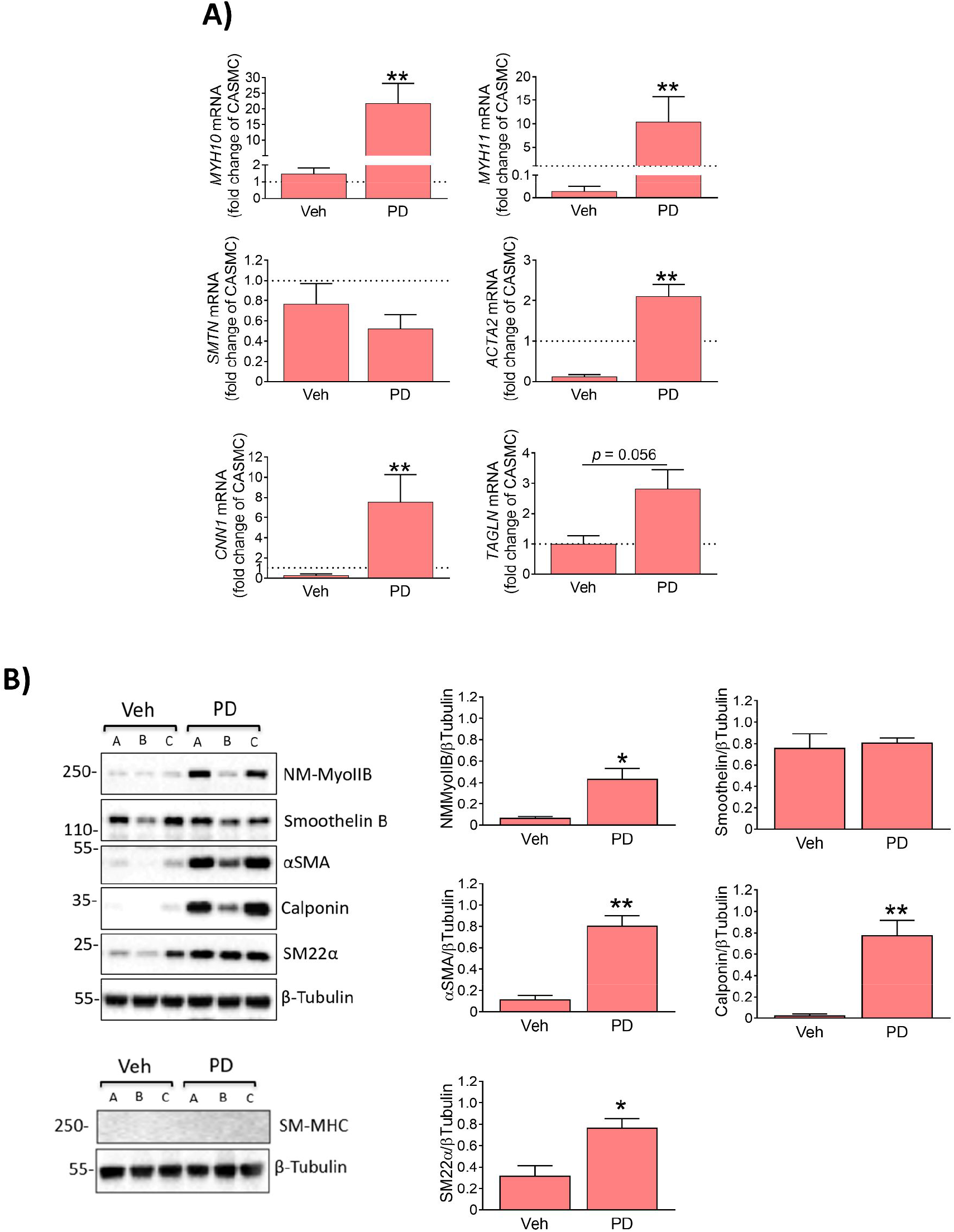
PD0325901 induces the human PC phenotypic switch into non-mature contractile VSMCs after 3 days. PCs were cultured for 3 days in the absence of GFs with either 250 nM PD0325901 or vehicle, with media replacement after 48h. Cells receiving the drug upregulated intermediate contractile VSMC markers, as showed by RNA **(A)** and protein **(B)** analyses. For RNA, dashed line indicates coronary artery SMCs (CASMCs) used as reference cell line. Values are expressed as fold change of CASMCs. N=5 per group. For blots, A, B, C are representative of cells from 3 different donors. Values are means ± SEM. \**P*<0.05, \*\**P*<0.01 vs Veh. **Veh**: vehicle. **PD**: PD0325901.

**Supplementary Figure 14.**
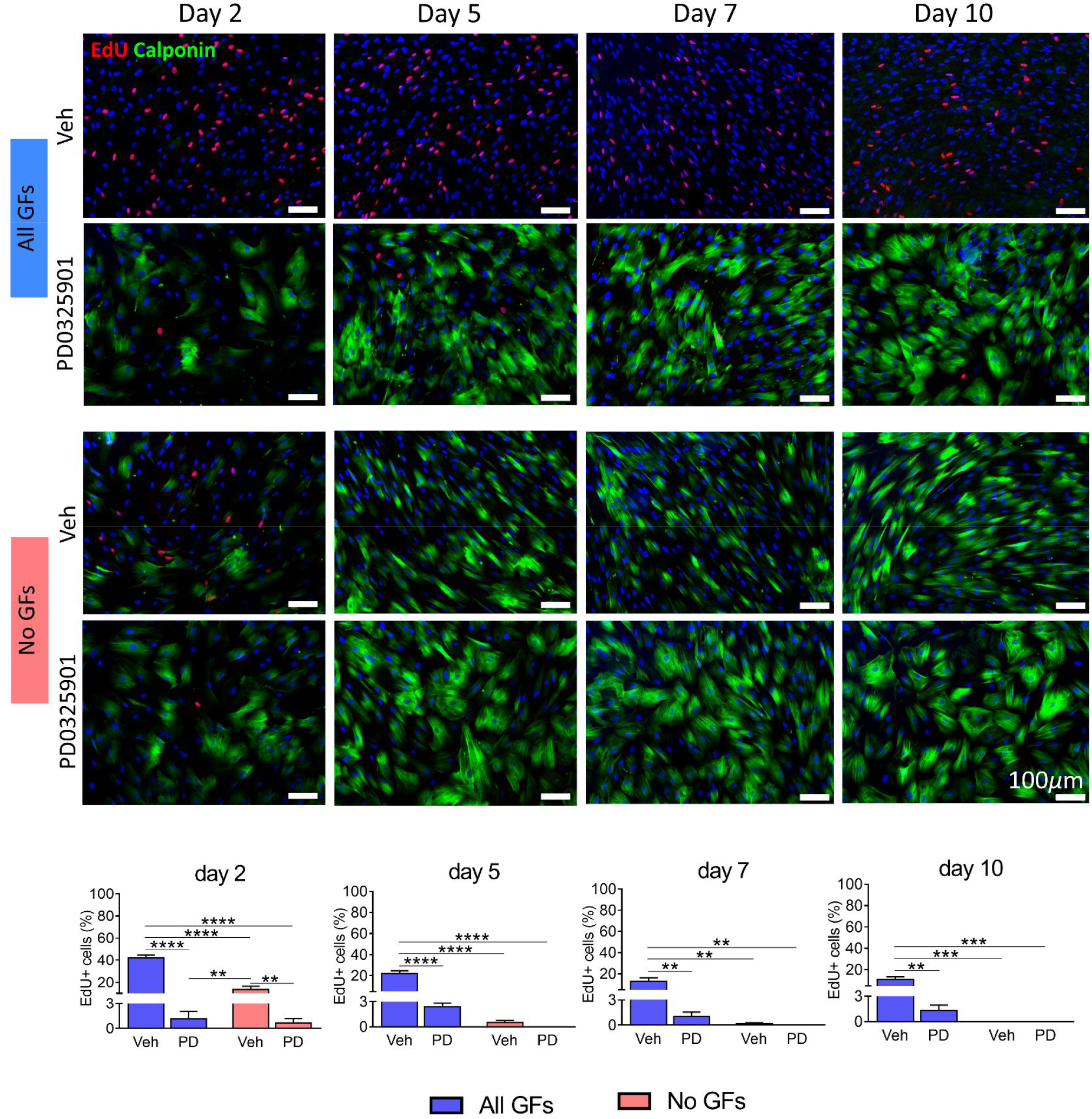
Proliferation assay with human cardiac PCs. PCs were cultured for 10 days with either *All GFs* or *No GFs*. Media were supplemented with 250 nanoM PD0325901 or its vehicle, with full replacement every 2 days. Cells were incubated with EdU for 24h at the times indicated. N=3 per each group. In the immunofluorescence images, EdU is shown in red, Calponin in green, and DAPI identifies nuclei. Values are means ± SEM. ** *P*<0.01, *** *P*<0.001, **** *P*<0.0001. **Veh**: vehicle. **PD**: PD0325901.

**Supplementary Figure 15.**
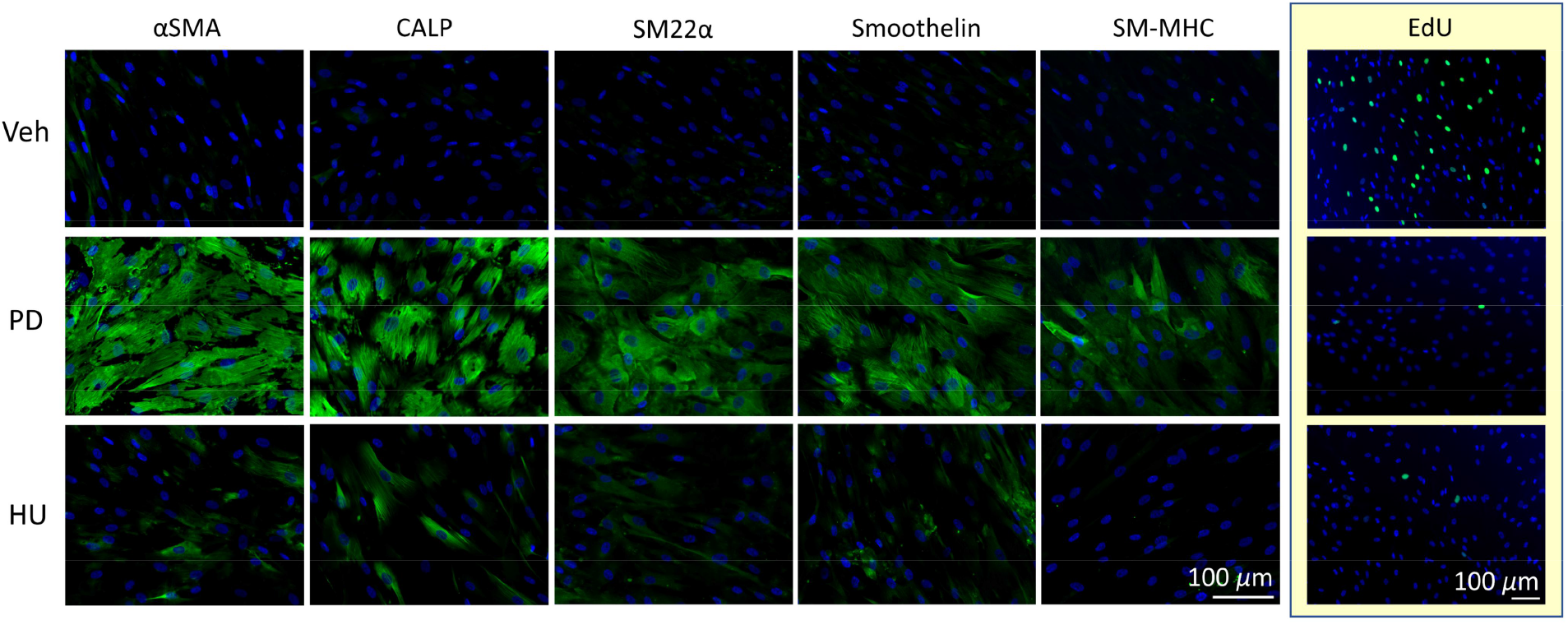
Test of the effects of cell cycle inhibition on human cardiac PC differentiation. PCs were cultured for 10 days with *All GFs*. Media were supplemented with either 250 nanoM PD0325901, or 2 mM hydroxyurea, or their vehicle (DMSO), with full replacement every 2 days. Cells were incubated with EdU for 24h at the end of the protocol. N=3 per each group. In the immunofluorescence images, VSMC markers and EdU are shown in green, DAPI identifies nuclei. **Veh**: vehicle. **PD**: PD0325901. **HU**: hydroxyurea.

**Supplementary Figure 16.**
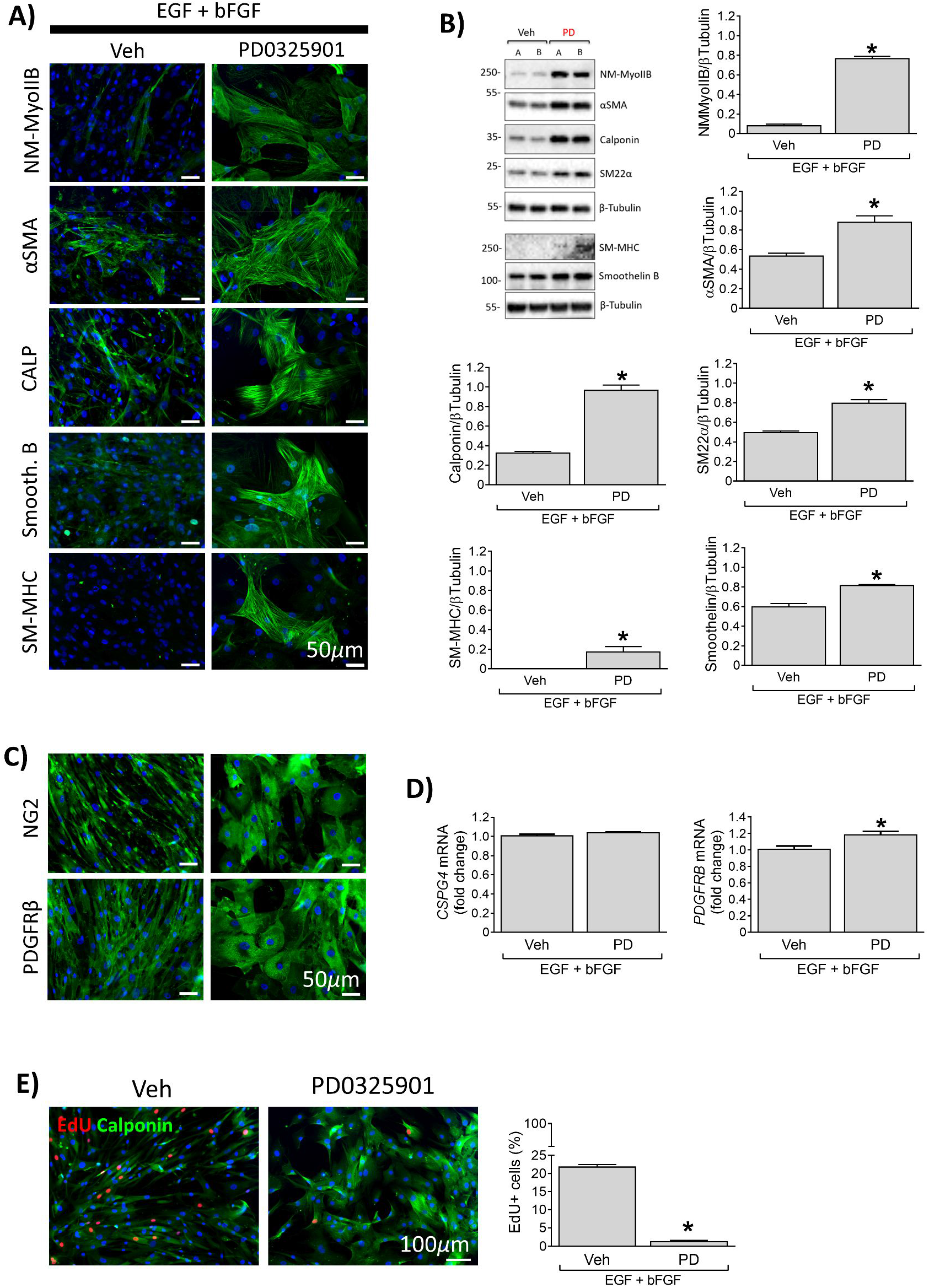
PD0325901 promotes the expression of contractile markers in human coronary artery SMCs (CASMCs). Cells were cultured for 10 days with the complete medium containing EGF and bFGF supplemented with either 250 nM PD0325901 or its vehicle, with media replacement every two days. Cells receiving the drug upregulated contractile VSMC markers, as showed by immunofluorescence **(A)** and western blotting **(B)** analyses. **(C-D)** Expression of pericyte/VSMC markers assessed by immunocytochemistry and qPCR. **(E)** Proliferation assay. Cells were incubated with EdU for 24h starting at day 3. EdU is shown in red, Calponin in green, and DAPI identifies nuclei in blue. In all experiments, N=4 per each group (N=2 donors, 2 replicates per each donor). For blots, A and B are representative of cells from 2 different donors. All values are means ± SEM. * *P*<0.05 vs Veh. **Veh**: vehicle. **PD**: PD0325901.

**Supplementary Figure 17.**
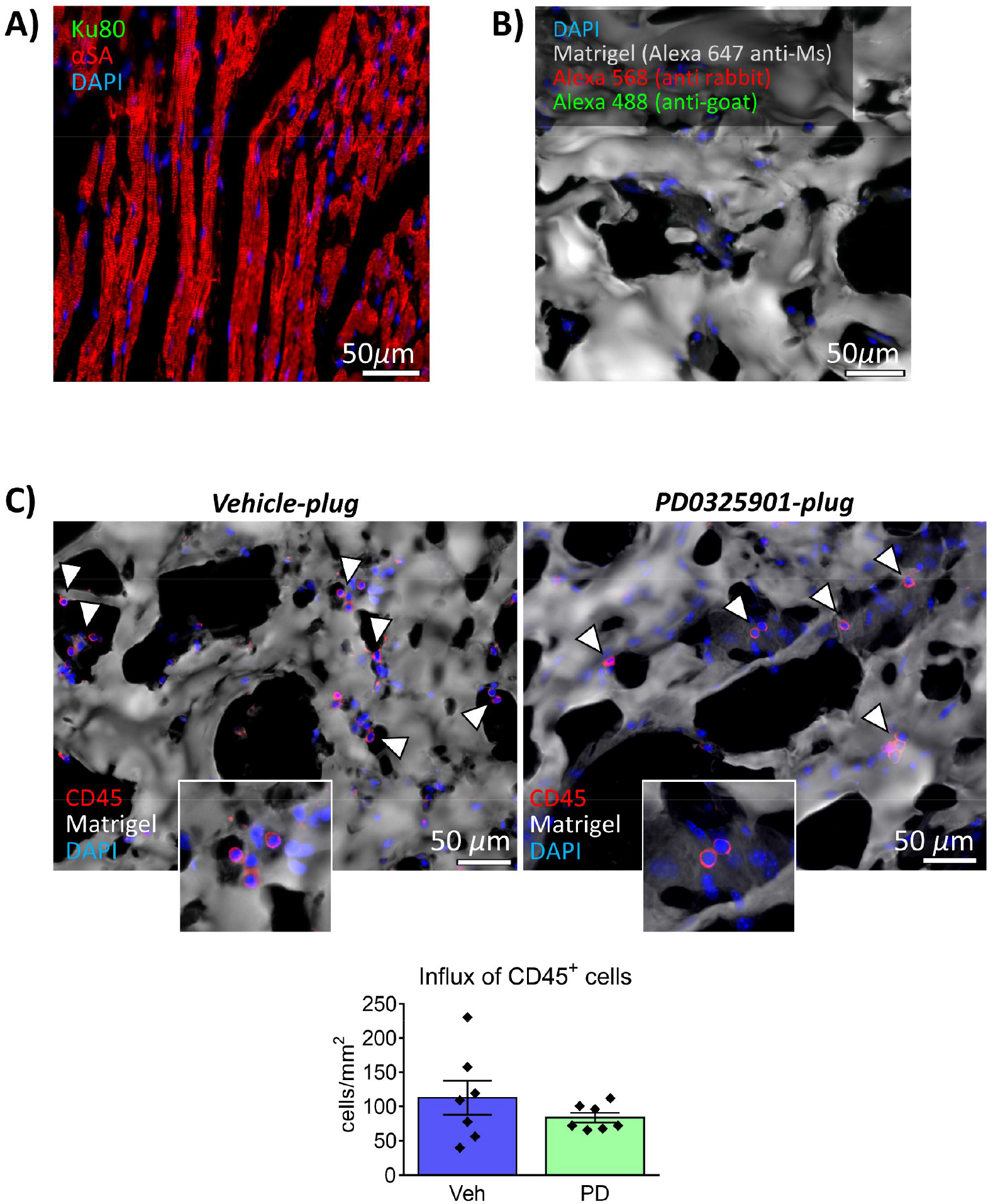
Matrigel plugs *in vivo* experiment. **(A)** Immunostaining for Ku80 in mice hearts ruled out the cross-reactivity of the antibody with mouse tissues. **(B)** Secondary antibody controls in Matrigel plugs. A secondary antibody Alexa 647 anti-mouse was used to evidence the structure of the matrix (Matrigel matrix from mouse sarcoma). **(C)** Influx of CD45+ immune/inflammatory cells in the Matrigel plugs. Immunofluorescence images showing CD45+ cells in red and the matrix in the pseudo-colour white. Graphs indicate the density of CD45+ cells in the plugs, expressed as number of cells per mm^2^ of area. N=7 per group. Values are means ± SEM. Results are not statistically significant. **Veh**: vehicle. **PD**: PD0325901.

**Supplementary Figure 18.**
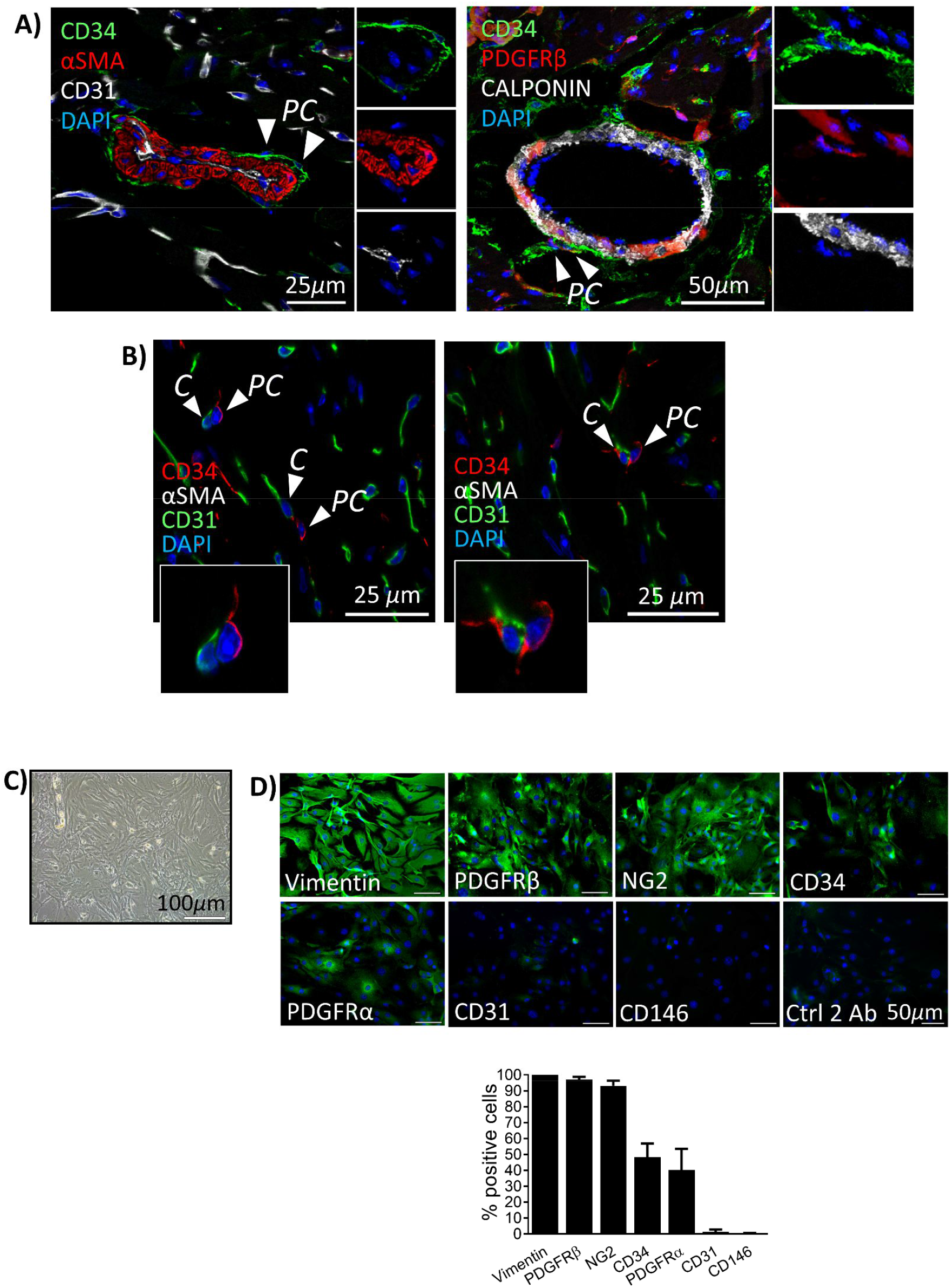
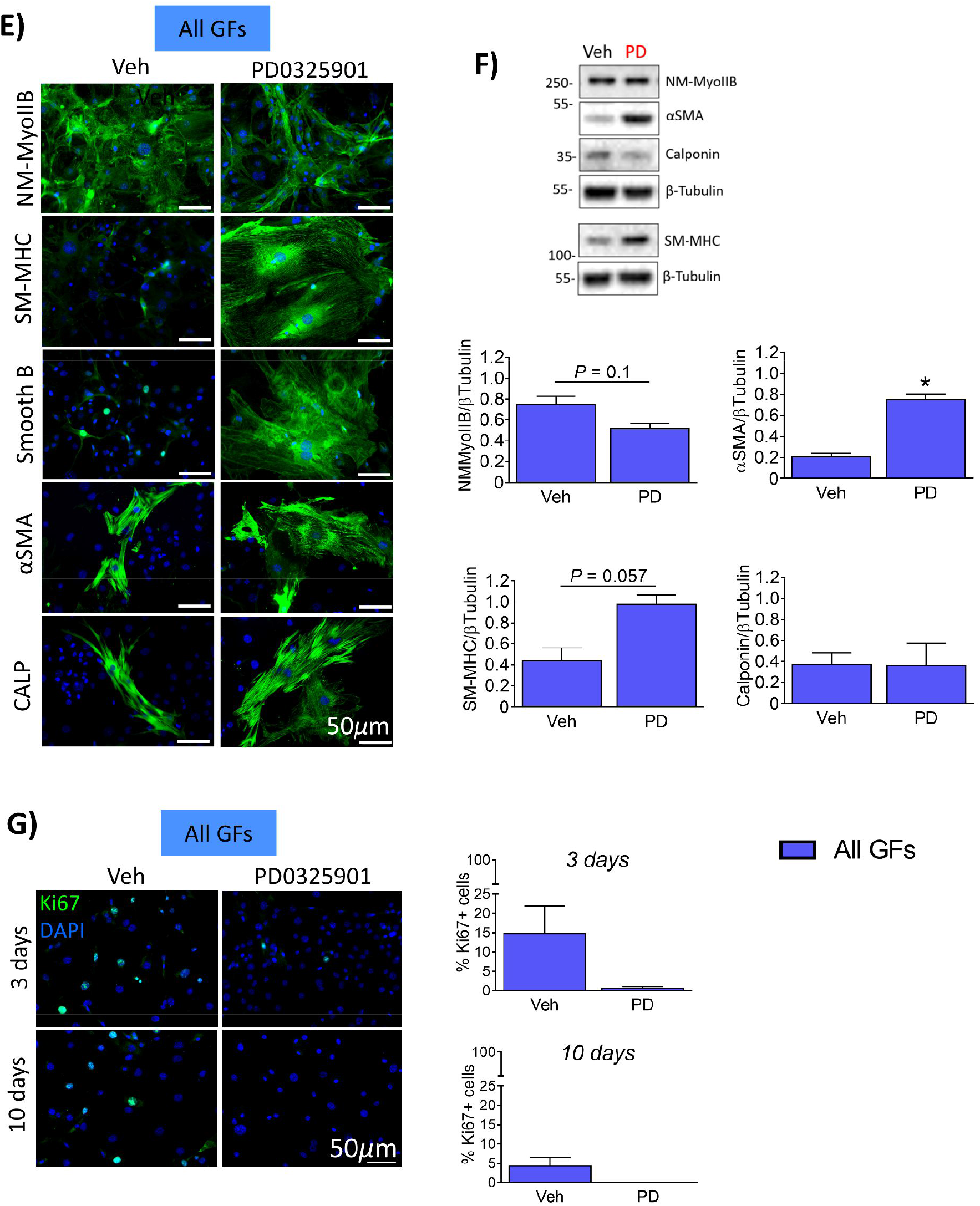
Effects of PD0325901 on murine cardiac PCs. **(A&B)** Confocal immunofluorescence images of mouse hearts. Arrows point to CD31neg αSMA/Calponin-neg CD34pos PDGFRβpos PCs around arterioles (A) and capillaries (B). C: capillary. PC: pericyte. **(C)** Brightfield microphotograph of expanded PCs at passage 4. **(D)** Antigenic profile of expanded PCs at passage 4. N=3. Values are means ± SEM. **(E&F)** PC differentiation into VSMCs using PD0325901. PCs were cultured with *All GFs* in the presence of PD0325901 or its vehicle for 10 days, with full media exchange every 2 days. (E) Immunofluorescence images show the expression of VSMC markers. (F) Blots showing a representative PC line and graphs showing results of blots densitometry. N=4. Values are means ± SEM. **\****p*<0.05 vs Veh. **(G)** PC proliferation after 3 and 10 days of PD0325901 treatment. Immunofluorescence images showing Ki67 in green fluorescence and graphs reporting quantitative analysis. N=3. Values are means ± SEM. **Veh**: vehicle. **PD**: PD0325901.

**Supplementary Figure 19.**
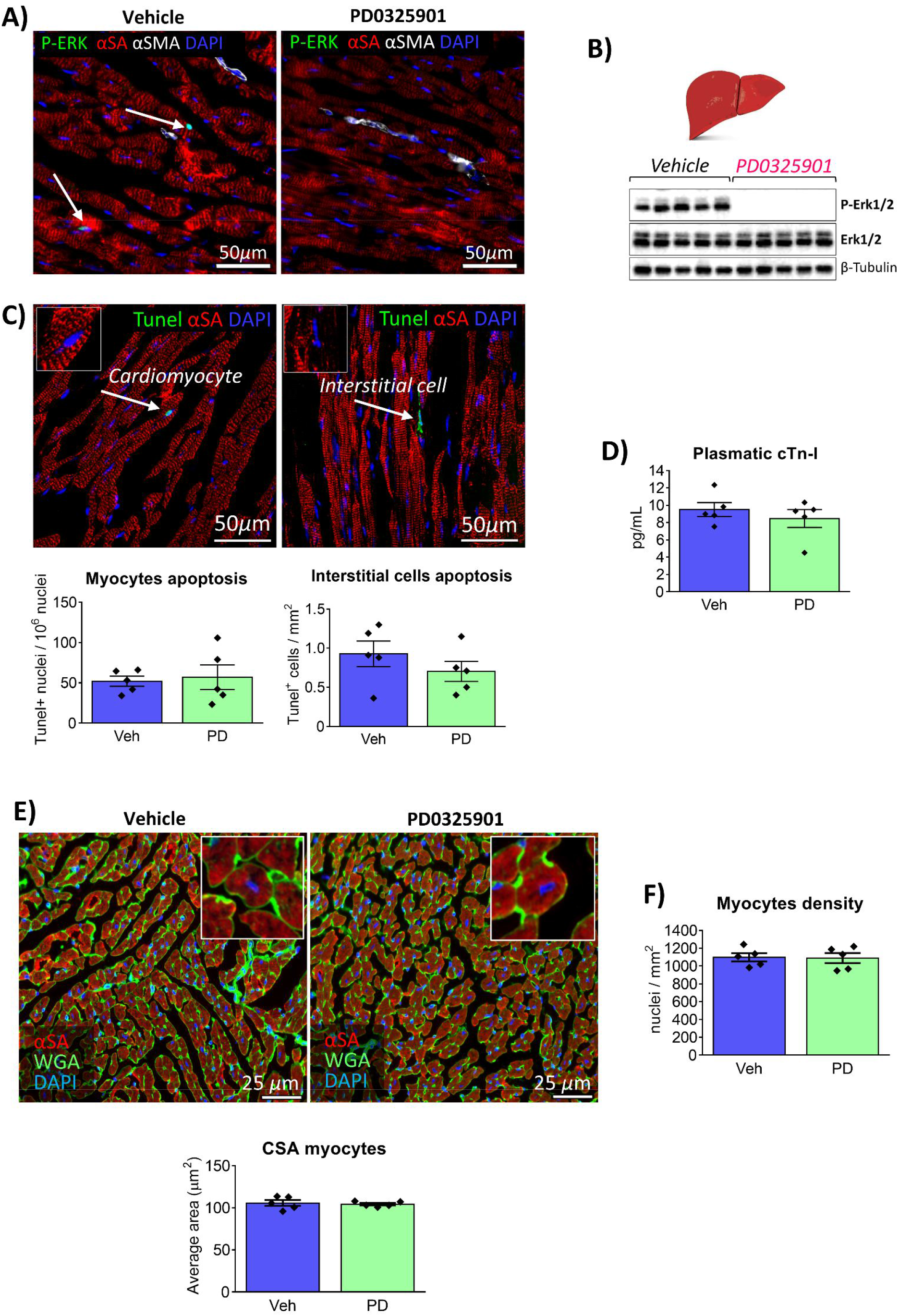
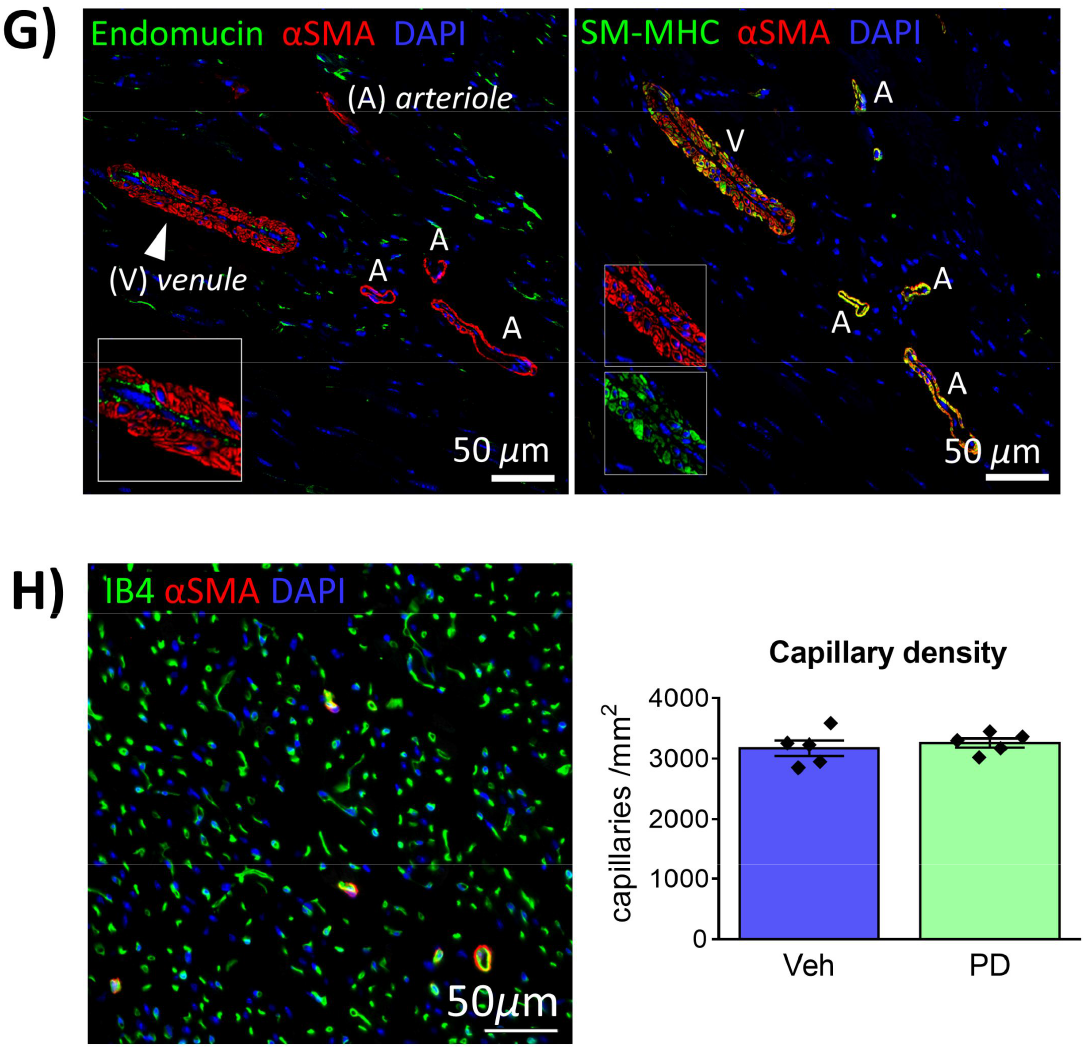
Efficacy and cardiac effects of PD0325901 treatment in healthy mice. **(A)** Absence of ERK1/2 phosphorylation in the hearts of PD-treated mice was confirmed by immunofluorescence staining. Arrows point to P-ERK-positive nuclei (green fluorescence) in the vehicle group. Cardiomyocytes are identified by the red fluorescence of α-Sarcomeric actin (α-SA). Nuclei in blue (DAPI). **(B)** Western blotting for P-ERK with total liver proteins showed systemic efficacy of the drug. **(C)** Apoptosis Tunel assay. Arrows point to apoptotic nuclei (green). Cardiomyocytes are identified by the red fluorescence of α-Sarcomeric actin (α-SA). Nuclei in blue (DAPI). Apoptotic cardiomyocytes are expressed as a fraction of total cardiomyocytes. Apoptotic α-SA-negative interstitial cells are expressed as number of cells per mm^2^ of myocardial area. N=5 per group. Values are plotted as single values and as means ± SEM. **(D)** ELISA for Cardiac Troponin T in mice plasma (n=5/group). According to guidelines for mice, healthy animals have cTn-I < 15 ng/mL. N=5 per group. Values are plotted as single values and as means ± SEM. **(E)** Analysis of the cardiomyocyte cross sectional area (CSA). Values are reported as average CSA per each mouse. Cardiomyocytes are identified by the red fluorescence of α-Sarcomeric actin (α-SA), Wheat Germ Agglutinin (WGA, green) was used to identify the cardiomyocytes borders for an accurate measurement of the cell area. Nuclei in blue (DAPI). CSA values are expressed in μm^2^. N=5 mice per group. Values are plotted as single values and as means ± SEM. **(F)** Cardiomyocyte density, expressed as the number of cardiomyocytes nuclei per mm^2^ of myocardial area. N=5 per group. Values are plotted as single values and as means ± SEM. **(G)** Example of venule identified for the luminal endothelium positivity to Endomucin. It is possible to observe that venules VSMCs are characterised by a higher expression of αSMA than SM-MHC. We only identified rare venules in each LV section. V: venule; A: arteriole. **(H)** Analysis of capillary density in the LV. The representative microphotograph shows capillaries in green (Isolectin-B4), arterioles in red (αSMA), and nuclei in blue (DAPI). Bar graphs show quantification of the capillary density. N=5 per group. Values are plotted as single values and as means ± SEM. **Veh**: vehicle, **PD**: PD0325901.

**Supplementary Figure 20.**
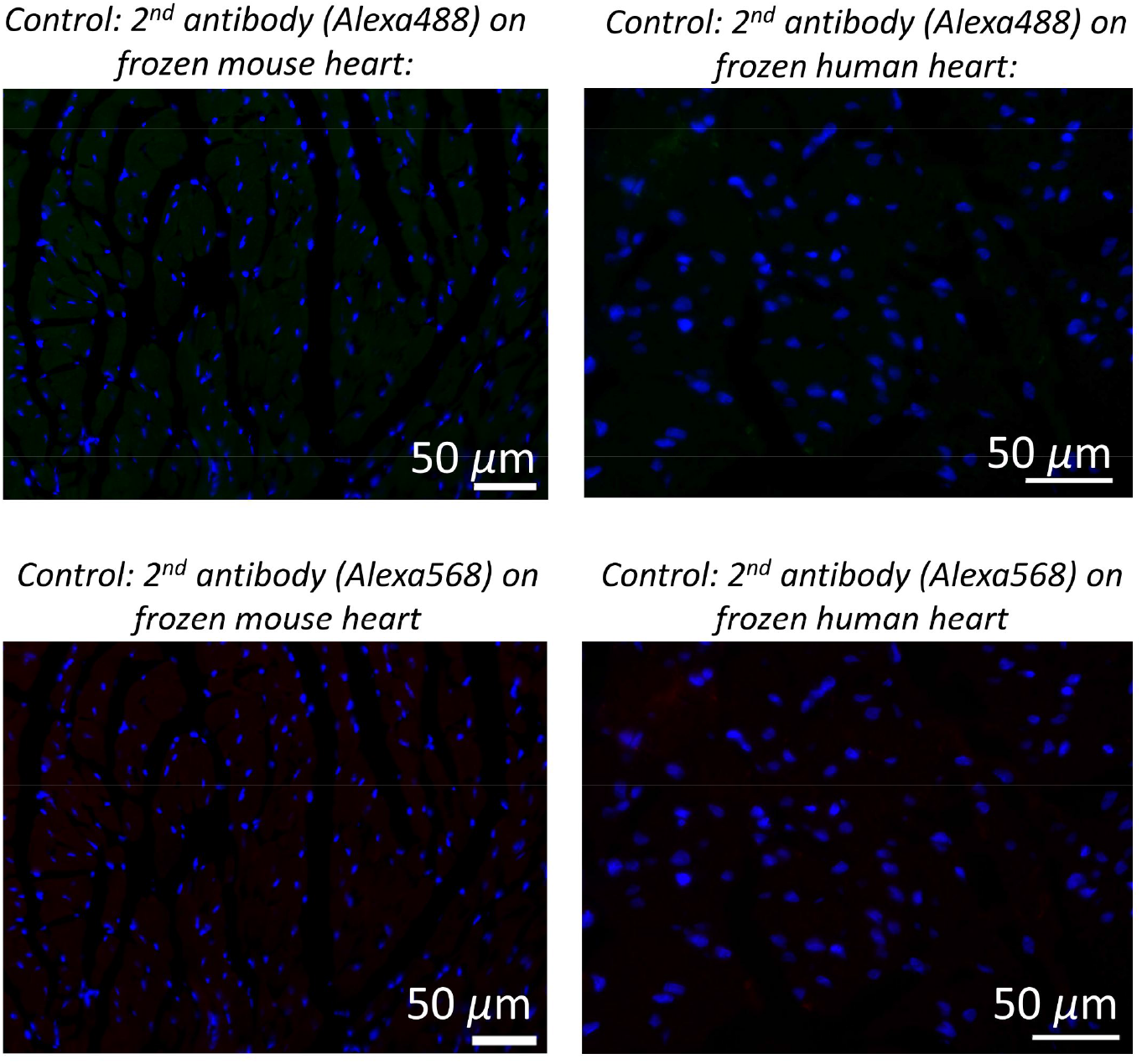
Controls for histological immunofluorescence analyses. Staining of mouse and human frozen hearts with the only secondary antibodies show the specificity of the secondary antibodies, as well as the low background autofluorescence of the tissues.

**Supplementary Table 1.** Congenital heart patients enrolled to the study.

| Cell line | Sample type | Sample weight (g) | Surgery | Patient age (years) | Experiments performed |
| --- | --- | --- | --- | --- | --- |
| PC 1 | RV | 0.100 | Pulmonary valve repair | 14 | *In vitro* |
| PC 2 | RV | 1.059 | Pulmonary valve repair | 15 | *In vitro, RNA-Seq* |
| PC 3 | RA | 0.190 | Atrial septal defect repair | 54 | *In vitro, RNA-Seq, In vivo* |
| PC 4 | RA | 0.085 | Atrial septal defect repair | 17 | *In vitro, RNA-Seq, In vivo* |
| PC 5 | RV | 0.120 | Tricuspid valve replacement + Pulmonary valve repair | 23 | *In vitro, In vivo* |
| PC 6 | RV | 2.700 | RV outflow tract resection | 56 | *In vitro, In vivo* |

RV right ventricle
RA right atrium

**Supplementary Table 2.** List of antibodies employed for analyses in human samples (including Matrigel plugs)

| ANTIGEN | HOST | COMPANY & CAT# | APPLICATION and DILUTION |  |  |
| --- | --- | --- | --- | --- | --- |
|  |  |  | IF (frozen) | ICC | WB |
| **CD34** | Sheep | R&D AF7227 | 1:100 |  |  |
| **CD34** | Mouse | Dako M7165 | 1:100 | 1:100 |  |
| **CD31** | Rabbit | Abcam Ab28364 | 1:50 | 1:20 |  |
| **CD31** | Mouse | R&D BBA7 | 1:100 |  |  |
| **CD146** | Rabbit | Abcam Ab75769 |  | 1:100 |  |
| **CD45** | Rat | NovusBio NB100-77417 | 1:500 |  |  |
| **NG2** | Rabbit | Millipore AB5320 | 1:50 | 1:50 |  |
| **PDGFRβ** | Goat | R&D AF385 | 1:50 | 1:50 |  |
| **PDGFRα** | Mouse | Santa Cruz sc-398206 |  | 1:100 |  |
| **Vimentin** | Rabbit | Abcam Ab92547 |  | 1:500 |  |
| **αSMA** | Mouse | Sigma, Cy3-conjugated C6198 | 1:400 | 1:200 |  |
| **αSMA** | Mouse | Dako, M0851 |  | 1:100 | 1:5000 |
| **CALPONIN** | Rabbit | Abcam Ab46794 | 1:100 | 1:100 | 1:10000 |
| **SMMHC (MYH11)** | Rabbit | Abcam Ab53219 | 1:100 | 1:50 |  |
| **SMMHC (MYH11)** | Rabbit | Invitrogen #702544 |  |  | 1:500 |
| **Smoothelin B** | Rabbit | Santa Cruz sc-28562 |  | 1:100 | 1:500 |
| **NMMyoIIb (MYH10)** | Rabbit | Abcam ab204358 |  | 1:200 | 1:1000 |
| **SM22α** | Mouse | Santa Cruz sc-271719 |  | 1:50 | 1:500 |
| **Ku80-XRCC5** | Goat | R&D AF5619-SP | 1:200 |  |  |
| **Elastin** | Mouse | Abcam ab77804 |  |  | 1:500 |
| **COL1A1** | Rabbit | Invitrogen #PA1-26204 |  |  | 1:1000 |
| **Fibronectin** | Rabbit | Abcam ab299 |  |  | 1:5000 |
| **P-ERK1/2** | Rabbit | Cell Signaling #4370 |  |  | 1:2000 |
| **ERK1/2** | Rabbit | Cell Signaling #4695 |  |  | 1:1000 |
| **P-EGFR** | Rabbit | Cell Signaling #3777 |  |  | 1:1000 |
| **EGFR** | Rabbit | Cell Signaling #4267 |  |  | 1:1000 |
| **P-FGFR** | Rabbit | Cell Signaling #3471 |  |  | 1:1000 |
| **FGFR1** | Rabbit | Cell Signaling #9740 |  |  | 1:1000 |
| **P-ELK1** | Rabbit | Cell Signaling #9181 |  |  | 1:1000 |
| **ELK1** | Rabbit | Cell Signaling #9182 |  |  | 1:1000 |
| **TCF21** | Rabbit | Abcam ab182134 |  |  | 1:1000 |
| **β-Tubulin** | Mouse | Cell Signaling #86298 |  |  | 1:5000 |

**Supplementary Table 3.**
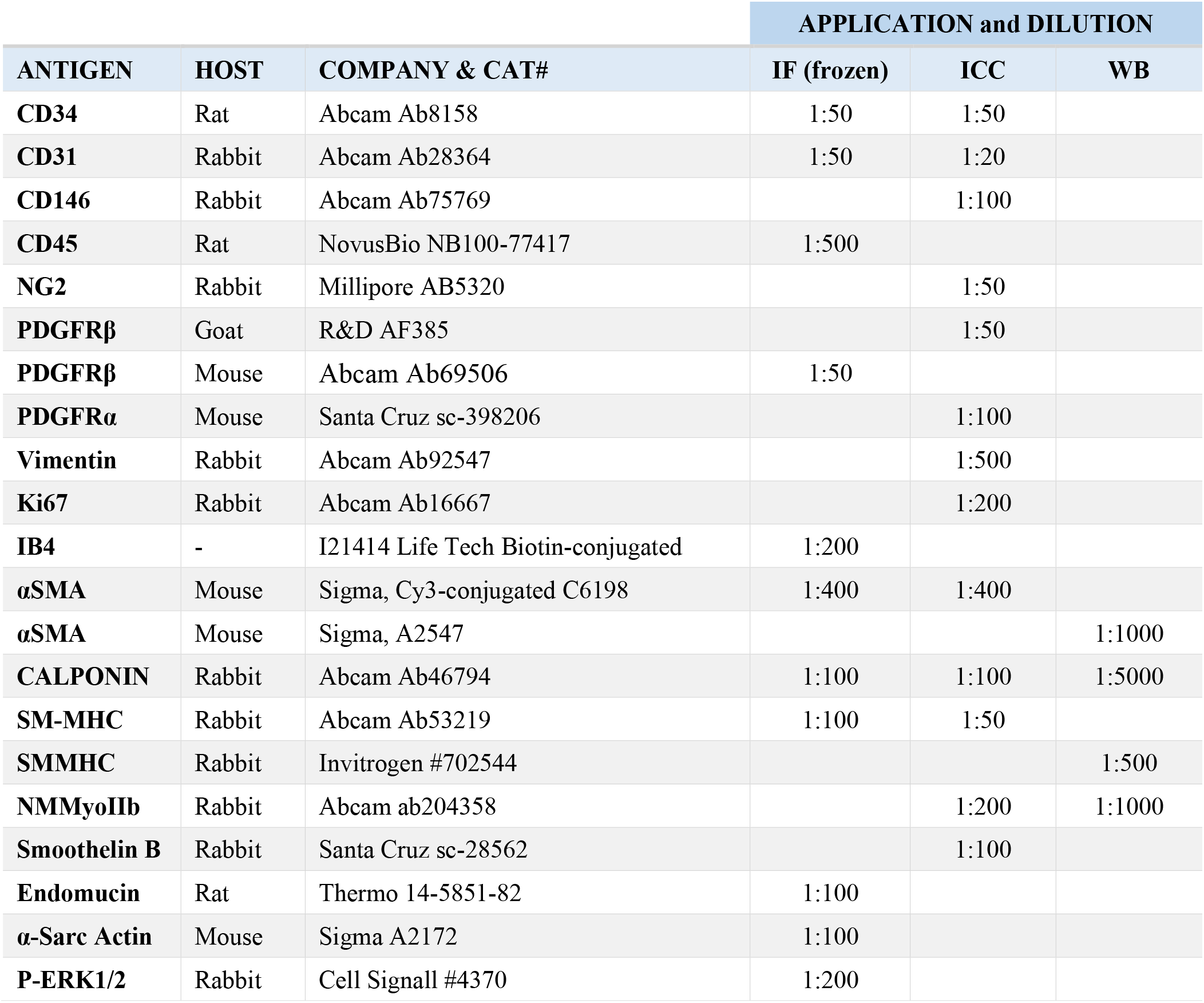
List of antibodies employed for analyses in mouse samples.

| ANTIGEN | HOST | COMPANY & CAT# | APPLICATION and DILUTION |  |  |
| --- | --- | --- | --- | --- | --- |
|  |  |  | IF (frozen) | ICC | WB |
| **CD34** | Rat | Abcam Ab8158 | 1:50 | 1:50 |  |
| **CD31** | Rabbit | Abcam Ab28364 | 1:50 | 1:20 |  |
| **CD146** | Rabbit | Abcam Ab75769 |  | 1:100 |  |
| **CD45** | Rat | NovusBio NB100-77417 | 1:500 |  |  |
| **NG2** | Rabbit | Millipore AB5320 |  | 1:50 |  |
| **PDGFRβ** | Goat | R&D AF385 |  | 1:50 |  |
| **PDGFRβ** | Mouse | Abcam Ab69506 | 1:50 |  |  |
| **PDGFRα** | Mouse | Santa Cruz sc-398206 |  | 1:100 |  |
| **Vimentin** | Rabbit | Abcam Ab92547 |  | 1:500 |  |
| **Ki67** | Rabbit | Abcam Ab16667 |  | 1:200 |  |
| **IB4** | - | I21414 Life Tech Biotin-conjugated | 1:200 |  |  |
| **αSMA** | Mouse | Sigma, Cy3-conjugated C6198 | 1:400 | 1:400 |  |
| **αSMA** | Mouse | Sigma, A2547 |  |  | 1:1000 |
| **CALPONIN** | Rabbit | Abcam Ab46794 | 1:100 | 1:100 | 1:5000 |
| **SM-MHC** | Rabbit | Abcam Ab53219 | 1:100 | 1:50 |  |
| **SMMHC** | Rabbit | Invitrogen #702544 |  |  | 1:500 |
| **NMMyoIIb** | Rabbit | Abcam ab204358 |  | 1:200 | 1:1000 |
| **Smoothelin B** | Rabbit | Santa Cruz sc-28562 |  | 1:100 |  |
| **Endomucin** | Rat | Thermo 14-5851-82 | 1:100 |  |  |
| **α-Sarc Actin** | Mouse | Sigma A2172 | 1:100 |  |  |
| **P-ERK1/2** | Rabbit | Cell Signall #4370 | 1:200 |  |  |

**Supplementary Table 4.**
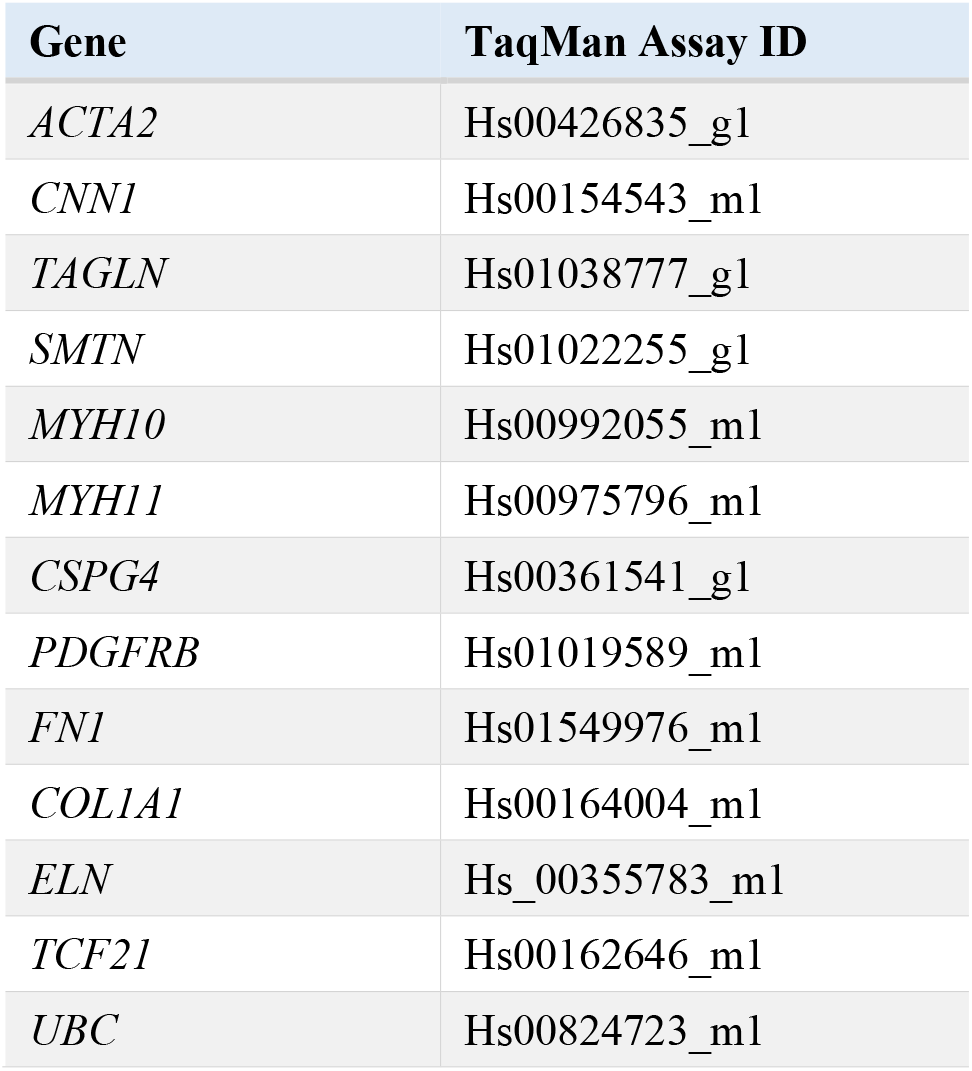
List of TaqMan probes used for expressional analyses in human cells.

| **Gene** | **TaqMan Assay ID** |
| --- | --- |
| *ACTA2* | Hs00426835_g1 |
| *CNN1* | Hs00154543_m1 |
| *TAGLN* | Hs01038777_g1 |
| *SMTN* | Hs01022255_g1 |
| *MYH10* | Hs00992055_m1 |
| *MYH11* | Hs00975796_m1 |
| *CSPG4* | Hs00361541_g1 |
| *PDGFRB* | Hs01019589_m1 |
| *FN1* | Hs01549976_m1 |
| *COL1A1* | Hs00164004_m1 |
| *ELN* | Hs_00355783_m1 |
| *TCF21* | Hs00162646_m1 |
| *UBC* | Hs00824723_m1 |

**Supplementary Table 5.** Main markers of synthetic/contractile VSMC phenotype investigated in the study.

| PROTEIN | GENE | FUNCTION | PHENOTYPE SPECIFICITY |
| --- | --- | --- | --- |
| **Non-muscle Myosin Heavy Chain II b (NM-MyoIIB)** | *MYH10* | Contractile filament<br>Actin-myosin force generation | *Synthetic* |
| **Smooth muscle α-actin (α-SMA)** | *ACTA2* | Contractile filament<br>Force generation | *Early-intermediate contractile* |
| **Transgelin (SM22α)** | *TAGLN* | Actin associated<br>Modulation of contraction | *Intermediate contractile*<br>Restricted to smooth muscle in adults |
| **Smooth muscle basic Calponin (CALP)** | *CNN1* | Actin associated<br>Regulation of contraction | *Intermediate contractile* |
| **Smoothelin B** | *SMTN* | Actin associated<br>Regulation of contraction | *Contractile*<br>Predominantly expressed in the vasculature of the adult. Specific to VSMC |
| **Smooth muscle myosin heavy chain (SM-MHC)** | *MYH11* | Contractile filament<br>Force generation | *Contractile*<br>The most specific marker of contractile VSMC. Rapidly lost by cultured VSMC |

**Supplementary Table 6.** Echocardiography indices (*In vivo* study 2 - Healthy mice) **Echocardiography indices in healthy mice given vehicle or treatment.** Echocardiography measurements were performed at baseline and after 14 days of treatment with vehicle or PD0325901. N=6/group. Values are means and SEM. *Statistics*: Two-way ANOVA was used to compare the effect of treatment and time and possible interaction. The two groups were balanced at baseline for all the measured parameters, except for CO and SV which required the use of analysis of covariance (ANCOVA). No significant differences attributable to treatment were observed.

|  | Vehicle |  | PD0325901 |  |
| --- | --- | --- | --- | --- |
|  | *Basal* | *Final* | *Basal* | *Final* |
| **E/A ratio** | 1.44 ± 0.11 | 1.73 ± 0.19 | 1.53 ± 0.11 | 1.41 ± 0.06 |
| **Heart rate (bpm)** | 441 ± 4 | 454 ± 4 | 442 ± 3 | 440 ± 5 |
| **Cardiac Output (mL/min)** | 12.7 ± 1.1 | 13.8 ± 0.9 | 13.5 ± 1.0 | 11.6 ± 0.9 |
| **LV Ejection Fraction (%)** | 67.9 ± 5.3 | 74.5 ± 2.7 | 69.2 ± 2.4 | 63.8 ± 3.0 |
| **Fractional Shortening (%)** | 37.8 ± 4.3 | 42.6 ± 2.6 | 38.1 ± 1.9 | 33.8 ± 2.2 |
| **Stroke volume (μL)** | 27.8 ± 2.5 | 30.8 ± 2.3 | 29.9 ± 2.1 | 24.8 ± 2.3 |
| **End Systolic Volume (μL)** | 15.9 ± 3.2 | 11.7 ± 0.5 | 14.7 ± 1.7 | 14.5 ± 2.4 |
| **End Diastolic Volume (μL)** | 41.1 ± 3.9 | 41.1 ± 1.6 | 47.5 ± 2.4 | 39.6 ± 4.2 |
| **LV ant wall thickness (μm)** | 0.72 ± 0.06 | 0.66 ± 0.02 | 0.80 ± 0.07 | 0.68 ± 0.06 |

**Supplementary Table 7.**
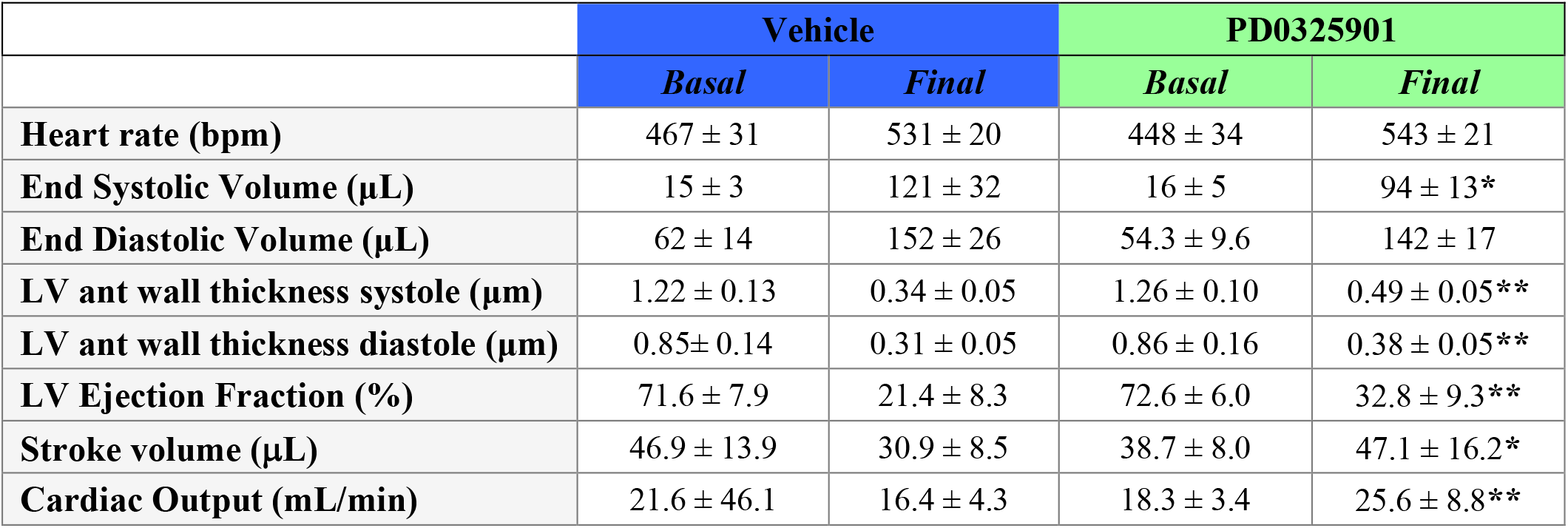
Echocardiography indices (*In vivo* study 3 - MI model) **Echocardiography indices in mice given vehicle or treatment after MI.** Echocardiography measurements were performed at baseline and after MI induction + 14 days of treatment with vehicle or PD0325901. N=8-12/Vehicle group and 11-12/PD group. Values are means and SD, **\*** *P*<0.05 and **\*\*** *P*<0.01 vs. Final vehicle. *Statistics*: The two groups were balanced at baseline for all the measured parameters.

|  | **Vehicle** |  | **PD0325901** |  |
| --- | --- | --- | --- | --- |
|  | *Basal* | *Final* | *Basal* | *Final* |
| **Heart rate (bpm)** | 467 ± 31 | 531 ± 20 | 448 ± 34 | 543 ± 21 |
| **End Systolic Volume (μL)** | 15 ± 3 | 121 ± 32 | 16 ± 5 | 94 ± 13* |
| **End Diastolic Volume (μL)** | 62 ± 14 | 152 ± 26 | 54.3 ± 9.6 | 142 ± 17 |
| **LV ant wall thickness systole (μm)** | 1.22 ± 0.13 | 0.34 ± 0.05 | 1.26 ± 0.10 | 0.49 ± 0.05** |
| **LV ant wall thickness diastole (μm)** | 0.85 ± 0.14 | 0.31 ± 0.05 | 0.86 ± 0.16 | 0.38 ± 0.05** |
| **LV Ejection Fraction (%)** | 71.6 ± 7.9 | 21.4 ± 8.3 | 72.6 ± 6.0 | 32.8 ± 9.3** |
| **Stroke volume (μL)** | 46.9 ± 13.9 | 30.9 ± 8.5 | 38.7 ± 8.0 | 47.1 ± 16.2* |
| **Cardiac Output (mL/min)** | 21.6 ± 46.1 | 16.4 ± 4.3 | 18.3 ± 3.4 | 25.6 ± 8.8** |

## Extended methods

Where not otherwise stated, all chemicals were purchased from Sigma-Aldrich.

## Frequently used abbreviations

Non-standard abbreviations and acronyms
bFGF: basic fibroblast growth factor
DAPI: 4′,6-diamidino-2-phenylindole
EGF: epidermal growth factor
FBS: Foetal bovine serum
IGF1: insulin-like growth factor 1
PBS: phosphate buffer saline
PC: pericyte
PFA: paraformaldehyde
VEGF: vascular endothelial growth factor
VSMC: vascular smooth muscle cell
ANGPT: angiopoietin
ANOVA: analysis of variance
ATP: adenosine triphosphate
CAECs: coronary artery endothelial cells
CALP: smooth muscle calponin
cTn-I: cardiac troponin I
DEGs: differentially expressed genes
DMSO: dimethyl sulfoxide
DPCs: PD0325901-differentiated PCs
ECGM2: growth factor enriched pericyte medium
ECM: extracellular matrix
EGFR: EGF Receptor
ELK1: E26 transformation-specific (ETS) Like-1 protein
ERK1/2: extracellular signal-regulated kinase 1/2
ET-1: endothelin-1
FGFR: FGF Receptor
GFs: growth factors
GO: gene ontology
HGF: hepatocyte growth factor
LAD: left anterior descending
LV: left ventricle
LVEF: LV ejection fraction
MEK1/2: dual specificity mitogen-activated protein kinase kinase
MEKi: MEK inhibitor
MI: myocardial infarction
NG2: neuron-glial antigen 2
NM-MyoIIB: non-muscle myosin IIB
PCs: pericytes
PDGF-BB: platelet-derived growth factor-BB
PDGFRα: platelet-derived growth factor receptor alpha
PDGFRβ: platelet-derived growth factor receptor beta
p-ERK1/2: phospho-ERK1/2
PPI: protein-protein interaction
SDF-1α: stromal derived factor-1 alpha
SM: smooth muscle
SM22α: smooth muscle protein 22-alpha
SMA-α: Smooth Muscle Actin-alpha
SM-MHC: smooth muscle myosin heavy chain
TCF21: transcription factor 21
TUNEL: terminal deoxynucleotidyl transferase dUTP nick end labelling
VSMCs: vascular smooth muscle cells

## Ethics

### Ethics regulating human studies

This study complies with the ethical guidelines of the Declaration of Helsinki. Human myocardial samples (right ventricle or atrium) were discarded material from surgical repair of congenital heart defects (ethical approval number 15/LO/1064 from the North Somerset and South Bristol Research Ethics Committee). Adult patients and paediatric patients’ custodians gave informed written consent. Donors and samples characteristics (n=6) are described in **Supplementary Table 1**.

### Ethics regulating animal studies

Procedures involving animals were performed under ethical licenses from the British Home Office (PPL numbers 30/3373 and PFF7D0506) and in compliance with the EU Directive 2010/63/EU, and following approval from Animal Ethics Committee at the University of Otago (AEC10/14), New Zealand. Procedures were carried out according to the principles stated in the Guide for the Care and Use of Laboratory Animals (The Institute of Laboratory Animal Resources, 1996).

Results are reported following the guidelines contained in the Animal Research Report of In Vivo Experiments (ARRIVE).

### Derivation of primary cultures of PCs

#### Human cardiac pericytes

Myocardial samples were collected in sterile, cold DMEM (Gibco) and processed within 24 h. The cell extraction protocol was as described previously^1^. Briefly, samples were finely minced using scissors and scalpel until nearly homogenous and digested with Liberase (Roche) for up to 1 hour at 37 C, with gentle rotation. The digest was passed through 70-, 40- and 30-μm strainers. Finally the cells were recovered and sorted using anti-human CD31 and anti-human CD34 microbeads (Miltenyi) to deplete the population of CD31+ endothelial cells and to select CD31-CD34+ cells. Cardiac PCs were cultured with full Endothelial Cell Growth Medium 2 (ECGM2, PromoCell) supplemented with recombinant human EGF (5 ng/mL), bFGF (10 ng/mL) and VEGF (0.5 ng/mL), R3-IGF1 (5 ng/mL), and 2% v/v FBS (all from Promocell) onto plastic plates. Cells were fed every 3 days and passaged at 90% confluence. Optimal seeding density is 8,000 cells/cm^2^. Frozen stocks of cells were preserved with 90% v/v FBS + 10% v/v DMSO. Centrifuge speed and time are 300g x 10 min. Unless otherwise stated, all experiments were performed between passage (P) 4 and 6. All cells tested negative for mycoplasma (assessed using a commercial PCR assay).

#### Mouse cardiac pericytes

C57BL/6 mice (n=5) were humanely killed under schedule 1 procedure and the heart harvested for cell extraction. Each heart was processed separately. Briefly, samples were finely minced using scissors and scalpel until nearly homogenous and digested with Liberase for up to 1 hour at 37 C, with gentle rotation. The digest was passed through 70-, 40- and 30-μm strainers. Final cells were recovered and sorted using anti-mouse CD31 microbeads (Miltenyi) and anti-mouse biotinylated-CD34 (Invitrogen) followed by anti-biotin microbeads (Miltenyi) to deplete CD31+ endothelial cells and select CD31-CD34+ cells. Cardiac PCs were cultured with basal ECBM2 medium (Promocell) supplemented with 10% v/v FBS, recombinant mouse EGF (5 ng/mL) and bFGF (10 ng/mL) (both from Peprotech) and Amphotericin B (Gibco, final concentration of 0.25 μg/mL), onto plastic plates pre-coated with collagen from bovine skin (final concentration of 30 μg/mL). Cells were fed every 3 days and passaged at 90% confluence. Optimal seeding density is 20,000 cells/cm^2^. Frozen stocks of cells were preserved with 90% v/v FBS + 10% v/v DMSO. Centrifuge speed and time are 300g x 5 min. Unless otherwise stated, all experiments were performed between passage 4 and 6.

### Cell biology studies

#### Culture of cardiac fibroblasts and ECs

Human cardiac fibroblasts were purchased from PromoCell and expanded in Fibroblast Growth Medium 2 (FGM2, PromoCell) according to manufacturer guidelines. Human coronary artery ECs (CAECs) were purchased from PromoCell and and expanded in full Endothelial Cell Growth MicroVascular medium 2 (ECGMV2, PromoCell) according to manufacturer guidelines. All cells were used between passage 4 and 6.

#### Immunocytochemistry analysis of human and mouse cells

Cells were rinsed with PBS and fixed with 4% w/v PFA in PBS for 15 min at 20°C. After washing with PBS, the cells were permeabilized with 0.1% v/v Triton-X100 in PBS for 10 min at 20°C as required. Cells were blocked with 10% v/v FBS and incubated with the antibodies as reported in **Supplementary tables 2&3**, for 16 h at 4°C. Secondary antibodies (conjugated with Alexa 488, Alexa 568, Alexa 647) were all purchased from ThermoFisher Scientific and used at a dilution of 1:200, for 1 h at 20°C, in the dark. Nuclei were counterstained using Hoechst (1:10,000 in PBS, 3 min at 20°C). Cells were mounted using Fluoromount.

#### 2D-Matrigel *in vitro* angiogenesis assay

Human coronary artery ECs (CAECs, purchased from PromoCell) were seeded on the top of Matrigel (Corning® Matrigel® Growth Factor Reduced Basement Membrane Matrix, cat# 356231) either alone (4,000 cells/well) or in co-culture with PCs (4,000 CAECs + 1,500 PC /well), using Angiogenesis μ-Slides (IBIDI, UK). Images were snapped after 5 hours, and the total tube length per imaging field was measured. To assess the interaction between PCs and CAECs, PCs were labelled with the red fluorescent tracker *Vybrant™ DiI Cell-Labeling Solution* (Invitrogen).

In experiments with only PCs, cells were seeded on the top of Matrigel, 4,000 cells/well. Images were snapped after 5 hours, and the total tube length per imaging field was measured.

#### Differentiation of human and mouse pericytes into VSMCs based on GFs

For experiments aiming to assess the role of the different GFs in the VSMC differentiation of PCs, species-specific GFs were added to, or removed from, the culture medium as indicated in the single experiments. For all experimental conditions, the concentration of FBS was never modified. Cells were cultured for a continuous period of 10 days with full media replacement every 48 h.

#### Differentiation of human coronary artery VSMCs based on GFs

Human coronary artery smooth muscle cells (CASMCs) were purchased from Promocell and cultured with the Smooth Muscle Cell Growth Medium 2 (SMCGM2) according to the vendor instructions. This medium is supplemented with recombinant human EGF and bFGF and 5% v/v FBS. For differentiation experiments, EGF and bFGF were depleted from the medium and the FBS concentration reduced to 2% (as per the pericytes). Cells were cultured for a continuous period of 10 days with full media replacement every 48h.

#### Pilot studies to determine PD0325901 efficacy and cytotoxicity

PD0325901 was purchased from Sigma-Aldrich and resuspended to a final stock concentration of 500 μM with DMSO. The compound was diluted in the culture medium as required, and DMSO used as vehicle control. Human PCs were cultured for ten days with growing concentrations of the drug (250 nM, 500 nM, 1 μM, 2 μM) to determine the effects on cell viability (Calcein-AM/EthDIII staining, from Biotium, and MTS assay, from Promega) and decide the maximum dose usable for the *in vitro* and *in vivo* cell transplantation experiments. Also, western blotting analysis of ERK1/2 phosphorylation was used to determine the minimum dose of the compound that effectively prevents ERK1/2 activation following incubation with stimulating GFs.

#### Differentiation of human and mouse cells into VSMCs with PD0325901

Cells were cultured with medium supplemented with either 250 nM PD0325901 or the same volume of DMSO (vehicle) for ten consecutive days, with complete medium exchange every 48 h. For shorter experiments, cells were cultured for only three days.

#### Effects of the cell cycle arrest on human PC differentiation

To investigate if the PC differentiation was driven by the arrest of cell proliferation, cells were cultured with full GFs medium supplemented with 2 mM hydroxyurea for ten consecutive days, with complete medium exchange every other day. The same volume of DMSO was used as control. The effective inhibition of cell proliferation was confirmed using an EdU assay (see below).

#### Single cell cloning assay with human PCs

Live PCs at P 3 were selected on their ability to exclude propidium iodide and sorted into cultures of single cells using a BD Bioscience Influx sorter (BD Biosciences). These sorted cells were cultured for up to 4 weeks in ECGM2 full medium in separate wells of a 96-well culture plate. Clones were passaged into bigger plates, with culturing continuing up to P6 before being used in the VSMC differentiation assay with/out GFs.

#### Contraction assay

A Cell Contraction Assay (CBA-201, from Cell Biolabs) was used to assess the capacity of differentiated PCs to contract upon stimulation with vasoconstrictors. At the end of the 10-days differentiation protocol, differentiated and control PCs were embedded in collagen gels following the manufacturer instructions. Endothelin-1 (ET1) was used at a concentration of 0.1 μM to stimulate cell contraction. Butanedione Monoxime (BDM) - contraction inhibitor provided with the kit - was added to the culture medium at a 1:100 dilution. Gels were released and the area of the gel measured at baseline and after 24h. Results are expressed as percentage of gel contraction.

#### Intracellular calcium flux assay

The Fluo-4 AM fluorescent calcium indicator (ThermoFisher Scientific) was employed to study the intracellular calcium flux in PCs and derived VSMCs following stimulation with a vasoconstrictor (Endothelin-1). Cell media was replaced with a buffer made of 20 mM HEPES pH=7.4, 137 mM NaCl, 5 mM KCl, 2 mM MgCl_2_, 1.8 mM CaCl_2_, 5.6 mM Glucose, 1 mg/mL bovine serum albumin, 0.5 mM NaH_2_PO_4_ and 1 mM NaHCO_3_. The Fluo-4 dye (2 μM) was loaded into the cells for 15 min. HOECHST was used to counterstain the nuclei. At the end of this incubation time, cells were stimulated with 0.1 μM ET-1 or its vehicle. Images were recorded every 20 sec for 10 min using an INCell Analyzer 2200 microscope (GE Healthcare) equipped with a 20 X objective. ET1 was injected in the cellular system after 120 sec. The intracellular calcium was measured as relative fluorescence units (RFU). Data analysis was carried out using the dedicated software In Cell Analyser Workstation 3.7.

#### Gap closure migration assay

The Radius^™^ 96-Well Cell Migration Assay (CBA-126 from Cell Biolabs) was used to assess the migratory properties of cardiac PCs and CASMCs at baseline and after differentiation. Stimuli used to induce cell migration were recombinant human VEGF-A (10 ng/mL), human SDF1α (100 ng/mL) and human PDGF-BB (20 ng/mL) (all from Peprotech). Migration time was 24h, in the presence of an inhibitor of cells proliferation (hydroxyurea, 2 mM). Absence of stimuli served as a control for migration. At the end of the protocol, cells were fixed with PFA and stained for VSMC markers (α-SMA or Calponin) and HOECHST (1:10,000 in PBS). The area of the gap not covered by cells was measured.

#### EdU proliferation assay

The Click-iT EdU Cell Proliferation Kit for imaging (C10337 - ThermoFisher Scientific) was used to assess cell proliferation. Cells were incubated with EdU for 24 h and then analyzed. Fluorescence staining for EdU was performed according to the manufacturer instructions.

### Molecular studies

#### Western blotting on human and mouse total cell lysates

Whole-cell protein lysates were collected on RIPA buffer supplemented with 1:50 proteases inhibitors cocktail and 1:100 phosphatases inhibitors. Cell lysates were centrifuged at 10,000 g, at 4°C, 15 min. After the assessment of protein concentration (BCA Protein Assay Kit, ThermoFisher Scientific), the supernatants were kept at −80°C. 10 to 30 μg of protein lysates were used as appropriate. Equal amounts of protein samples were prepared in Laemmli loading buffer, incubated for 8 min at 98°C, resolved on 8-12% SDS-PAGE and transferred onto 0.2 μm PVDF or nitrocellulose membranes (Bio-Rad). Membranes were blocked using 5% w/v BSA or 5% w/v non-fat dried milk (Bio-Rad) in Tris-buffered saline (TBS, BioRad) supplemented with 0.05% v/v Tween-20 for 2 h at 20°C. Primary antibodies (listed in **Supplementary Tables 2&3**) were incubated for 16 h at 4°C. Either β-tubulin or β-Actin were used as a loading control. Anti-rabbit IgG or anti-mouse IgG HRP-conjugated antibodies were employed as secondary antibodies (both 1:5000, GE Healthcare). Membrane development was performed by an enhanced chemiluminescence-based detection method (ECL™ Prime Western Blotting Detection Reagent, GE Healthcare) and observed using a ChemiDoc-MP system (Bio-Rad). Proteins with similar MW were assessed on different gels. No more than one stripping procedure was performed on an individual membrane (RestoreTM Plus Western Blot Stripping Buffer; Thermo Fisher Scientific). Western blot data were analyzed using the ImageJ software.

#### Western blotting on mouse liver samples

Freshly frozen liver samples were homogenized in RIPA buffer supplemented with proteases and phosphatases inhibitors using gentleMACS M tubes (Miltenyi). Tissue lysates were centrifuged at 10,000 g, at 4°C, 15 min and processed as described above.

#### Gene expression analysis by real-time qPCR in human PCs

Extracted total RNA was reverse-transcribed into single-stranded cDNA using a High Capacity RNA-to-cDNA Kit (ThermoFisher Scientific). The RT-PCR was performed using first-strand cDNA with TaqMan Fast Universal PCR Master Mix (ThermoFisher Scientific). The assay numbers for the endogenous controls and target transcripts are listed in **Supplementary Table 4**. TaqMan primer-probes were all obtained from ThermoFisher Scientific. Quantitative PCR was performed on a QuantStudio^™^ 5 System (ThermoFisher). All reactions were performed in a 10 μL volume in triplicate, using 7.5 ng cDNA per reaction. The mRNA expression levels were determined using the 2^-ΔΔCt^ method.

#### Signalling studies in human cardiac PCs

For the study of signalling activated by specific GFs, human PCs were cultured for 48-72 h with ECGM2 medium depleted of all GFs. Cells were then stimulated for 30 min or 1 h with specific combinations of GFs (EGF+bFGF, or VEGF+IGF1) or control. The whole-cell protein lysates were then collected for western blotting or protein arrays. A Human Phospho-kinase array kit (R&D Systems, catalogue number ARY003B) was used for quick screening of 43 kinase phosphorylation sites.

#### Enzyme-linked immunosorbent assay (ELISA)

For measurement of GFs in PC-conditioned medium, confluent cells were cultured for 48 h with basal ECBM2 without FBS and GFs. Media was then collected, centrifuged and used in the experiments. Quantikine ELISA Kit anti-human VEGF, Angiopoietin-1, Angiopoietin-2 and HGF were purchased from R&D Systems. The amount of secreted factors was normalized to the final number of cells and time of incubation. Cardiac troponin I was measured in mouse plasma using the Mouse cTn-I ELISA kit from Elabscience (E-EL-M1203).

#### Next-generation RNA-Sequencing

A whole-transcriptome analysis was carried out in 3 human PC lines differentiated for 10 days using PD0325901 and compared with the respective naïve controls. CASMCs (N=2) were used for reference. Strand specific RNA-sequencing (seq) was performed starting from total RNA (GENEWIZ, New Jersey, US).

RNA-Seq was carried out on an Illumina HiSeq platform, with a 2×150bp configuration, ∼20M reads per sample. Sequence reads were trimmed to remove possible adapter sequences and nucleotides with poor quality using Trimmomatic v.0.36. The trimmed reads were mapped to the Homo sapiens GRCh38 reference genome available on ENSEMBL using the STAR aligner v.2.5.2b, and BAM files generated. Unique gene hit counts were calculated by using featureCounts from the Subread package v.1.5.2. Only unique reads that fell within exon regions were counted. After extraction of gene hit counts, the gene hit counts table was used for downstream differential expression analysis. Using DESeq2, a comparison of gene expression between the groups of samples was performed. The Wald test was used to generate p-values and log2 fold changes. Genes with a false discovery rate (FDR) < 0.05 and absolute log2 fold change > 1 were called as differentially expressed genes (DEGs) for each comparison. A gene ontology (GO) analysis was performed on the statistically significant set of genes by implementing the software GeneSCF v.1.1-p2. The goa_human GO list was used to cluster the set of genes based on their biological processes and determine their statistical significance.

Extended gene annotations, hierarchical clusterings, MA-plots, pathway analyses, heatmaps were generated using the open-access bioinformatics softwares iDEP 9.1 (http://bioinformatics.sdstate.edu/idep/)^2^ and Morpheus (https://software.broadinstitute.org/morpheus). Venn diagrams were generated using InteractiVenn (http://www.interactivenn.net/index.html).^3^ Protein-protein interaction networks were generated using the STRING software (https://string-db.org/cgi/input?sessionId=bM1QcP9Hl6oN&input_page_show_search=on).

### *In vivo* studies

Three independent studies were conducted in mice, all following randomized controlled protocols.

#### Study 1: In vivo Matrigel plugs with human PCs

Cold Matrigel (Corning® Matrigel® Growth Factor Reduced Basement Membrane Matrix, cat# 356231) was mixed with human PCs (2×10^6^) and treated with PD0325901 (500 nM) or vehicle (DMSO). Twelve-week-old male C57BL6/J mice (Charles River) (n=8) were injected subcutaneously into both abdominal flanks with 400 μL Matrigel, under anaesthesia induced by isoflurane inhalation. Seven days later, the animals were sacrificed, and the Matrigel plugs were harvested, fixed in 4% PFA at 4°C overnight, immersed in 30% w/v sucrose in PBS overnight and embedded in OCT for histological analyses. 15-μm thick sections were cut for immunostainings.

#### Study 2: Administration of PD0325901 to healthy mice

Seven-week-old female C57BL6/J mice (Charles River) (n=22) were housed in individual cages, in an enriched environment within a bio-secure unit under a 12 h light/dark cycle, fed with EURodent Diet (5LF5, LabDiet) and given drinking water ad libitum. PD0325901 was given orally and voluntarily each day by inclusion of the compound in sugar-free strawberry-flavoured jelly, as previously described.^4, 5^

#### PD0325901-jelly preparation

The jelly was prepared every 2 days, stored at 4° C and used within 48 h to ensure the stability of the drug. Mice were conditioned to eat the jelly for 5 days (twice daily) before the start of the 14-day experimental protocol. PD0325901 was given to mice (n=11) at 10 mg/kg body weight once daily; it was dissolved in DMSO and incorporated within the jelly. The control group (n=11) received DMSO-jelly. 50 μL DMSO/drug were dissolved in a final volume of 4 mL jelly, so that the final dose of DMSO was not toxic for the animals. Mice were given jelly 8 μL/g body weight. Individual housing was necessary to observe jelly consumption.

All of the mice ate the entire jelly during the experiment; therefore none were excluded from the study. A cohort of mice (n=6/group) underwent echocardiography at baseline and at the end of the protocol. Cardiac dimensional and functional parameters were measured with using a high-frequency, high-resolution, 3D echocardiography system (Vevo3100, Fujifilm, VisualSonics) using an MX550D transducer, with mice under isoflurane anaesthesia to maintain the heart rate at 450 bpm. Animals were terminated 2-to-4 hours after the last treatment (jelly administration). Hearts (n=5/group) were stopped in diastole using KCl, perfusion-fixed (PBS-EDTA followed by 4% PFA) and harvested. Hearts were then weighed and fixed in 4% w/v PFA at 4°C overnight, immersed in 30% w/v sucrose in PBS overnight and embedded in OCT (Tissue-Tek® O.C.T. Compound, VWR) for cryopreservation and histological analyses. 5-μm thick sections were cut for immunostainings. Livers (n=5/group) were harvested and freshly frozen in liquid N2 for molecular biology. Blood (n=5/group) was collected with EDTA for plasma separation and analysis of circulating biomarkers.

#### Blood flow measurement

Left ventricle myocardial blood flow was assessed in the same mice that had undergone echocardiography (n=6 mice/group), according to previously published protocols.^6-8^ Briefly, carboxylate-modified green-fluorescent microspheres (cat# F8813, Invitrogen, 0.5μm diameter) were injected in the left ventricle cavity over one minute, for a total volume of 200 μL, and flushed with 150 μL of warmed saline. Reference blood samples were obtained from the descending aorta over a time of 2 min. The left ventricle, kidneys and reference blood were collected for subsequent recovery of microspheres and fluorescence reading. The weigth of the organ post-harvest was recorded. Regional blood flow was calculated as the absolute blood flow in ml/min/g of tissue using the formula described earlier.^7, 8^ The Kidneys were analyzed to confirm the uniform distribution of fluorescent microspheres in the systemic circulation.

#### Study 3: Administration of PD0325901 in a mouse model of myocardial infarction (MI)

This study was conducted in 8 weeks old C57/Bl6 mice (Hercus). Mice were acclimatized with sugar-free strawberry-flavoured jelly (prepared as above) for five days before surgery. MI was induced by permanent ligation of the left anterior descending coronary artery (LAD) as described before.^7^ In brief, following anaesthesia (2,2,2 tribromo ethanol, 0.3gm/kg, i.p.) and artificial ventilation, the chest cavity was opened and, after careful dissection of the pericardium, LAD was located and permanently ligated using a 7-0 silk suture. After confirming for the absence of bleeding, chest cavity was closed in layers. Animals were allowed to recover for at least 4 hours before returning to the housing unit. Mice were monitored twice a day for the first 5 days post-surgery and thereon once every day. All the mice received analgesic and antibiotic from just before the surgery for the next 3 days. Following 3 days of recovery post-MI, mice were randomized to receive the sugar-free strawberry-flavoured jelly with or without PD0325901 for next 14 days. Cardiac function was assessed using echocardiography measurement at baseline (before MI) and at 14 days post-MI. At the end of the treatment period and following echocardiography, and under anaesthesia, heart were stopped in diastole using KCl, and ventricular tissue was collected following perfusion-fixation with 4% PFA. The collected tissue were used to determine the effect of PD0325901 on angiogenesis and fibrotic remodelling following MI as described below.

### Histology on human and mouse tissues

#### Histological procedures

Human myocardial samples were fixed in 4% PFA at 4°C overnight, immersed in 30% w/v sucrose in PBS overnight and embedded in OCT for cryopreservation and histological analyses. 5-μm thick sections were cut for identification of cardiac PCs *in situ*. Mice samples were prepared as described above. Human and mouse frozen sections were post-fixed and permeabilized with ice-cold acetone (VWR) for 5 min at −20 °C, and let air dry for 30 min. Sections were rehydrated with PBS for 10 min. For P-ERK staining, the protocol required permeabilization with ice-cold methanol (VWR) for 10 min at −20 °C. Tissue sections were blocked with 5% v/v FBS or normal goat serum (as appropriate) and incubated with primary antibodies for 16h at 4°C.

To determine the capillary density, 5-μm thick cryosections were incubated with biotinylated isolectin B4 (Griffonia Simplicifolia Lectin I (GSL I) isolectin B4, Biotinylated, B-1205, Vector Laboratories) to stain endothelial cells overnight at 4°C. Antibodies are reported in the **Supplementary Tables 2&3**. Secondary antibodies (Alexa 488-, Alexa 568-, Alexa 647, streptavidin Alexa-488-conjugated) were all purchased from ThermoFisher Scientific and used at a dilution of 1:200, for 1h at 20°C except for streptavidin Alexa-488 to detect endothelial cells which was incubated for 4h at 20°C in the dark. To determine the arteriole density For smooth muscle cells identification to determine the arteriole density, 5-μm thick cryosections were incubated with Cy3 conjugated monoclonal anti-alpha smooth muscle actin for 4 hrs at room temperature along with the streptavidin Alexa-488. Slides were mounted using ProLong™ Gold Antifade Mountant with DAPI (ThermoFisher Scientific).

For the analysis of *apoptosis* in mice hearts, the ApopTag® Fluorescein In Situ Apoptosis Detection Kit was employed, following manufacturer instructions.

Alexa-488 conjugated wheat germ agglutinin (WGA) (ThermoFisher) (30 min, 1:200 dilution) was used to identify cardiomyocytes borders for morphometric analyses.

To identify the fibrotic tissue in the samples from the MI study, Azan Mallory staining was performed as previously described.^7, 9^

Whenever new antibodies, not previously employed and tested in our lab, were adopted, the specificity of the antibodies was tested on positive (and negative where available) control samples following the manufacturer’s guidelines. The specificity of the secondary antibodies Alexa-Fluor conjugated, as well as the background autofluorescence of the tissues, are shown in **Supplementary Figure 20**.

### *Histological analyses* of human and mouse tissues

#### Study 1 - PC-Matrigel plugs

- ***Expression of VSMC markers by human PCs***: The Ku80-XRCC5 antigen was utilized to recognize the human cells. Cytoplasmatic staining ruled out cell senescence.^10, 11^ Antibodies anti-αSMA, Calponin and SM-MHC were used to recognize VSMC proteins. Results are expressed as percentage of human PCs expressing the indicated VSMC marker. N=2 sections were analyzed per each sample.
- ***Infiltration of immune/inflammatory cells***: an antibody anti-CD45 was employed to assess the presence of immune cells within the plugs. Results are expressed as number of CD45+ cells per mm^2^ of plug area. N=2 sections were analyzed per each sample.

#### Study 2 - Healthy mice

- ***Arteriole density***: Small and large arterioles density was expressed as the number of arterioles per mm^2^. Arterioles were classified as small (< 20μm) and large (> 20μm) according to their lumen size. Venules were recognized for positivity to Endomucin and excluded from the analysis.^12^ All arterioles present in an entire section of LV was analyzed per each mouse. Considering we could detect only rare venous vessels with a very bright αSMA fluorescence like arteries, and this number was uniform across the two groups, the Endomucin marker was not used for all the following analyses. Normally, defining the microscope fluorescence settings to identify arteries, veins are not detected due to the remarkably low levels of fluorescence.
- ***Arterial VSMC maturation***: this was evaluated as the ratio between the areas occupied by SMMHC and αSMA. Briefly, RGB images, acquired keeping the microscope settings unvaried, were segmented into three grey-scale channels using ImageJ, the background subtracted, and an automatic threshold was calculated and applied for each channel using the ImageJ built-in algorithms to measure the area of the green and red signal for each vessel. Final data are expressed as a volume fraction between the areas occupied by SMMHC and αSMA. Ten random small and large arterioles were analyzed per each mouse, for a final N=50 total arterioles/group. Per each mouse, the average of all measurements was calculated.
- ***Coverage of arterioles with CD34***^***pos***^ ***cells***: this was measured as the ratio between the areas occupied by perivascular CD34 and αSMA of the arterial wall, with the same procedure described above. N=10 random small and large arterioles were analyzed per each mouse, for a final N=50 total arterioles/group. Per each mouse, the average of all measurements was calculated.
- ***Arterioles surrounded by CD34***^***pos***^ ***cells***: this was expressed as a percentage of total arterioles. All arterioles included in an entire section of LV were analyzed per each mouse.
- ***Capillary density***: this was evaluated in ten random fields (snapped using a 10x objective) in which capillaries were transversally oriented. Per each mouse, it was expressed as the number of capillaries per square millimetre of myocardial tissue.
- ***Cardiomyocytes cross-sectional area (CSA)***: the CSA was measured in cardiomyocytes in which the nucleus was centrally located within the cell. All suitable cardiomyocytes were evaluated in two entire LV sections. Data are reported as the average CSA per each mouse, and as a distribution of all individual CSA.
- ***Cardiomyocytes density:*** the number of cardiomyocytes nuclei was counted in 10 random fields, snapped at 100x, from two LV sections. Results are expressed as number of cardiomyocyte nuclei per mm^2^ of myocardial area.
- ***Cardiomyocytes apoptosis:*** Apoptotic cardiomyocytes were expressed as the number of Tunel+ cardiomyocytes per 100,000 cardiomyocytes. Two entire sections of the LV were analyzed.
- ***Interstitial cells apoptosis:*** Tunel+ αSA-negative interstitial cells were expressed as the number of cells per square millimetre of myocardial area.

#### Study 3 - Mouse MI model

- ***Capillary and arteriole density***: capillaries and arterioles were calculated in 6 fields at X200 magnification and the final data expressed as the number of capillaries or arterioles per square millimetre. Quantification was performed in the peri-infarct area and in the remote myocardium separately. Arterioles were categorised according to their luminal size.
- ***Infarct size***: Data were expressed as the percentage of LV occupied by the fibrotic scar (Azan Mallory staining). A whole LV section was analysed per each mouse.

#### Microscopy equipment and post-imaging analysis

Most of the analyses were carried out on photomicrographs obtained using an Axio Observer Z1 (Zeiss) equipped with 10 X and 20 X objectives. For confocal imaging, we used a Leica SP5-II AOBS multi-laser confocal laser scanning microscope attached to a Leica DM I6000 inverted epifluorescence microscope (Leica Microsystems), equipped with a 20 X, 40 X and 63 X immersion oil objectives. Images processing was performed using the dedicated ZEN PRO and Leica LAS X software, and Photoshop CC (Adobe). Images analyses were performed using the open-access ImageJ (NIH) software. A 3D reconstruction of multiple z-stacks acquisitions was performed during the Matrigel plug experiment.

### Statistical analyses

Continuous variables are presented as means ± standard error of the mean (SEM) or standard deviation (SD) of independent samples and/or as individual values. The D’Agostino-Pearson and Kolmogorov– Smirnov normality tests were used to check for normal distribution when applicable. Continuous variables normally distributed were compared using the Student’s t-test (two-group comparison) or one-way analysis of variance (ANOVA; for multiple group comparisons). Two-way ANOVA analyses were used to compare the mean differences between groups when appropriate. Non-parametric tests, including the Mann–Whitney U test (two-group comparison) and the Kruskal-Wallis test (multiple group comparison) were used to compare data not normally distributed. Post-hoc analyses included Tukey and Dunn tests, as appropriate. Echocardiography parameters (baseline and final assessed in the same animal) were compared using paired tests; for all other analyses, unpaired tests were applied. For *in vivo* studies, post-hoc analyses of outcomes were conducted according to the intention-to-treat principle. In Study 2, when baseline echo measurements were found to differ between groups, the analysis of covariance (ANCOVA) was used, as it provides the optimum statistical analysis in terms of bias, precision, and statistical power. In Study 3, where final values were missing due to the early animal mortality, we used a mixed-effects model to analyze repeated measures data with missing values (available in GraphPad Prism 8.0). Significance was assumed when p ≤ 0.05. Analyses were performed using GraphP
ad Prism 6.0 and 8.0.

